## supplementary materials for "Assessment of Polygenic Architecture and Risk Prediction based on Common Variants Across Fourteen Cancers"

**SUPPLEMENTARY INFORMATION**

**INDEX**

1.0 **Supplementary Tables**
2.0 **Supplementary Figures**
3.0 **Supplementary Note**
    3.1. **References**
    3.2. **Consortium-Specific Funding and Acknowledgments**
    3.3. **Consortium-Specific Collaborators**

**1.0 SUPPLEMENTARY TABLES**

**Supplementary Table 1.** Sample sizes for summary-level GWAS data used for analysis of 14 cancers.

| Group^a^ | Cancer site | Number of SNPs analyzed after filtering in the model | Number of cases | Number of controls | Chip imputation Information |
| --- | --- | --- | --- | --- | --- |
| 1 | CLL^b1^ | 1,068,036 | 3,100 | 7,667 | Infor score >0.3 |
|  | Esophageal^c2^ | 1,068,728 | 3,914 | 6,718 | Info score >0.4 |
|  | Testicular^3^ | 1,066,591 | 3,558 | 13,970 | Info score >0.3 |
|  | Oropharyngeal^4^ | 1,067,193 | 6,034 | 6,585 | Call rate >98% |
|  | Pancreas^5^ | 915,805 | 8,638 | 12,217 | Info >0.3 |
| 2 | Renal^6^ | 1,067,952 | 10,784 | 20,407 | Info >0.3 |
|  | Glioma^7^ | 1,067,960 | 12,488 | 18,169 | Info >0.4 |
|  | Melanoma^8^ | 1,052,042 | 12,874 | 23,203 | Quality R^2>0.95 |
|  | Colorectal^9^ | 1,058,067 | 17,050 | 19,529 | Call rate >95%, R^2 >0.7 |
|  | Endometrial^10^ | 1,068,132 | 12,906 | 108,979 | Info score>0.4 |
|  | Ovarian^11^ | 1,068,810 | 22,406 | 40,951 | Call rate >95% |
| 3 | Lung^12^ | 1,009,906 | 29,266 | 56,450 | R^2>0.3, info >0.4 |
|  | Prostate^13^ | 806,185 | 79,148 | 61,106 | R^2>0.8 |
|  | Breast^14^ | 1,067,502 | 108,067 | 88,386 | Info score >0.3 |

^a^Group 1, group 2, and group 3 include <10K, 10-25K, and >25K cases in the analysis, respectively. ^b^CLL = chronic lymphocytic leukemia. ^c^Includes Barrett’s esophagus and esophageal adenocarcinoma cases.

**Supplementary Table 2.** **Number of independent genome-wide significant SNPs (P<** $\boldsymbol{5\times}\boldsymbol{10}^{\boldsymbol{-8}}$**) and associated heritability explained in the current datasets and those projected based on the estimated effect-size distribution using sample sizes of the current datasets**. The number of independent SNPs reaching genome-wide significance in current studies were based on LD-clumping with an $r^{2}$-threhold of 0.1 and 1 MB window size. The heritability explained ($h_{e}^{2})$associated with these SNPs were calculated using the formula $h_{e}^{2}=\sum_{i} {(\hat{\beta}}_{i}^{2}-\tau_{i}^{2})$ where $\hat{\beta}_{i}$ is the estimate of log-odds ratio (in standardized scale) and $\tau_{i}$ is the corresponding standard error for the $i$-th SNP.

|  | Number of genome-wide significant SNPs observed | Observed $\boldsymbol{h}_{\boldsymbol{e}}^{\boldsymbol{2}}$ of the genome-wide significant SNPs | GENESIS projections of number of genome-wide significant SNPs (95%CI) | GENESIS projection of $\boldsymbol{h}_{\boldsymbol{e}}^{\boldsymbol{2}}$ of the genome-wide significant SNPs (95%CI) |
| --- | --- | --- | --- | --- |
| CLL^a^ | 20 | 0.559 | 16 (8, 29) | 0.53 (0.31, 0.83) |
| Esophageal | 1 | 0.012 | 0 (0, 5) | 0.00 (0.00, 0.03) |
| Testicular | 38 | 0.908 | 44 (28, 79) | 1.10 (0.81, 1.59) |
| Oropharyngeal | 0 | 0.000 | 0 (NA) | 0.00 (NA) |
| Pancreas | 11 | 0.128 | 11 (5, 25) | 0.13 (0.07, 0.24) |
| Renal | 16 | 0.116 | 12 (5, 28) | 0.10 (0.05, 0.17) |
| Glioma | 33 | 0.488 | 22 (13, 35) | 0.44 (0.33, 0.55) |
| Melanoma | 26 | 0.265 | 28 (16, 48) | 0.27 (0.20, 0.36) |
| Colorectal | 8 | 0.039 | 8 (3, 19) | 0.03 (0.01, 0.08) |
| Endometrial | 13 | 0.047 | 12 (5, 34) | 0.05 (0.03, 0.12) |
| Ovarian | 12 | 0.056 | 16 (7, 34) | 0.07 (0.04, 0.12) |
| Lung | 15 | 0.066 | 8 (3, 18) | 0.06 (0.04, 0.08) |
| Prostate | 144 | 0.392 | 127 (102, 169) | 0.37 (0.33, 0.41) |
| Breast | 169 | 0.278 | 149 (120, 192) | 0.27 (0.24, 0.29) |

^a^CLL = chronic lymphocytic leukemia.

**2.0. SUPPLEMENTARY FIGURES**

**Supplementary Fig. 1. Q-Q plots comparing observed distributions of GWAS summary statistics against those expected under the fitted GENESIS models across 8 different cancers.** Plots in upper and lower panels are generated under the two- and three-component models, respectively. The two-component model assumes the non-null effect sizes to follow a single normal distribution. The three-component model assumes the non-null effect sizes to follow a mixture of two normal distributions with two distinct variance components. Shaded regions mark 80% point-wise confidence intervals derived from 100 simulations. $\lambda_{obs}$ is the genomic control factor in the observed summary-level GWAS data; $\lambda_{fit}$ is the mean genomic control factor in simulated data over 100 replications. The ranges for all plots are restricted to be p-value < ${10}^{-10}$.

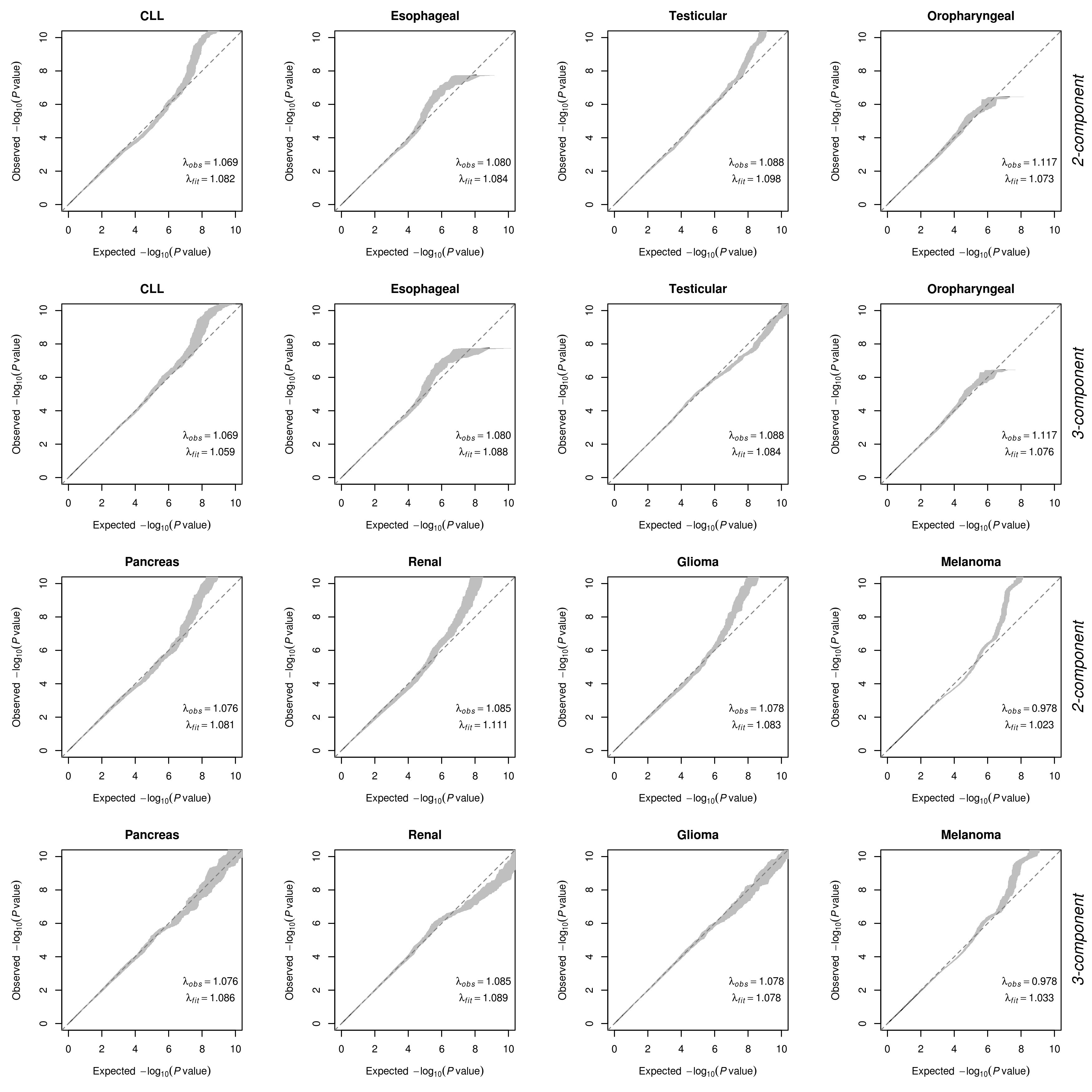

CLL = chronic lymphocytic leukemia.

**Supplementary Fig. 2.** **Q-Q plots comparing observed distributions of GWAS summary statistics) against those expected under the fitted GENESIS models across 6 different cancers**. Plots in upper and lower panels are generated under the two- and three-component models, respectively. The two-component model assumes the non-null effect sizes to follow a single normal distribution. The three-component model assumes the non-null effect sizes to follow a mixture of two normal distributions with two distinct variance components. Shaded regions mark 80% point-wise confidence intervals derived from 100 simulations. $\lambda_{obs}$ is the genomic control factor in the observed summary-level GWAS data; $\lambda_{fit}$ is the mean genomic control factor in simulated data over 100 replications. The ranges for all plots are restricted to be p-value < ${10}^{-10}$.

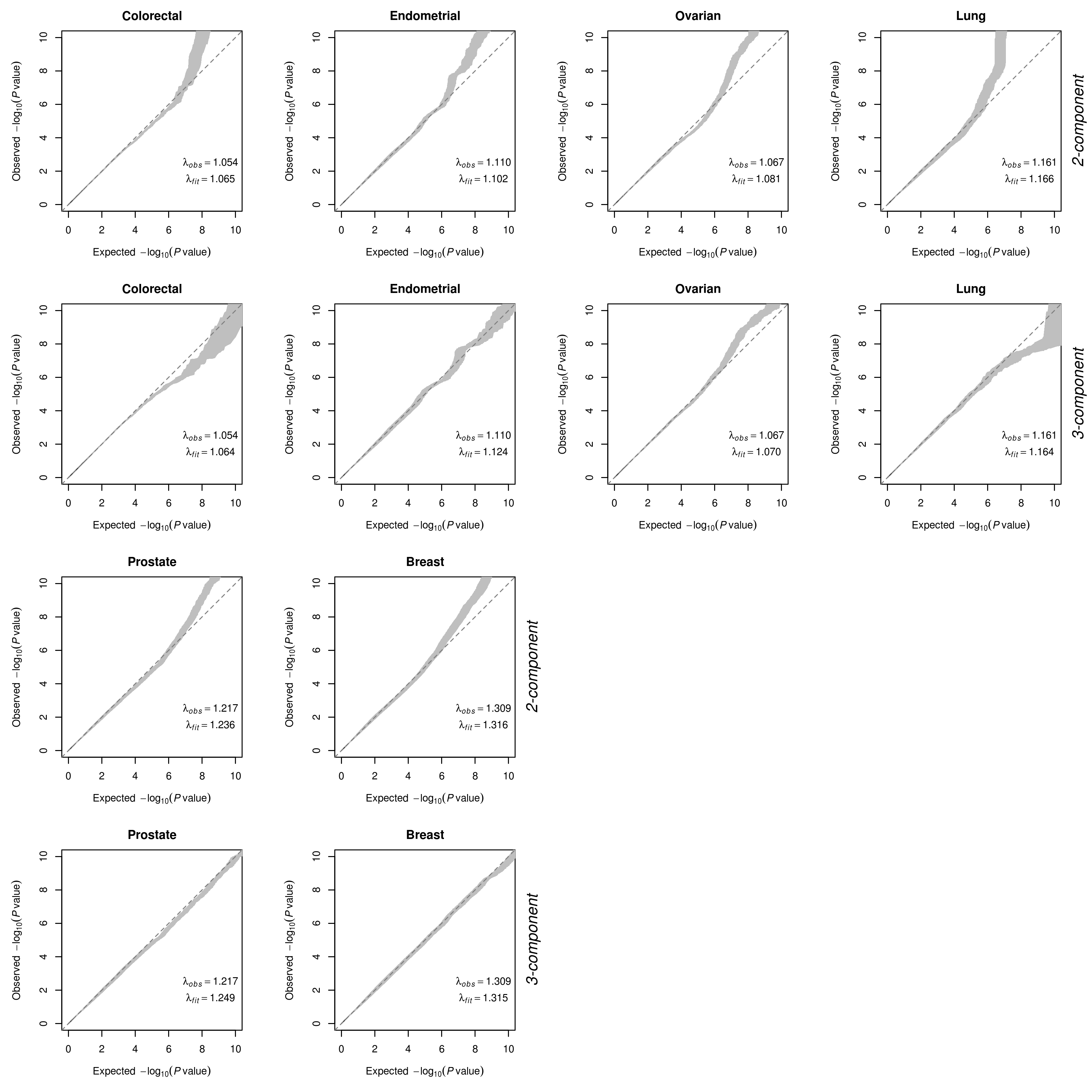

**Supplementary Fig. 3. Projections of area under the curve (AUC) characterizing predictive performance of PRS as sample size for GWAS increases.** Results are shown for PRS including SNPs at the optimized p-value threshold (solid curve) and at genome-wide significance (P<$5\times{10}^{-8}$) level (dashed curve). The dotted horizontal red line indicates the maximum AUC achievable according to the estimate of GWAS heritability. Colored dots correspond to sample size for largest published GWAS and those for doubled and quadruped sizes.

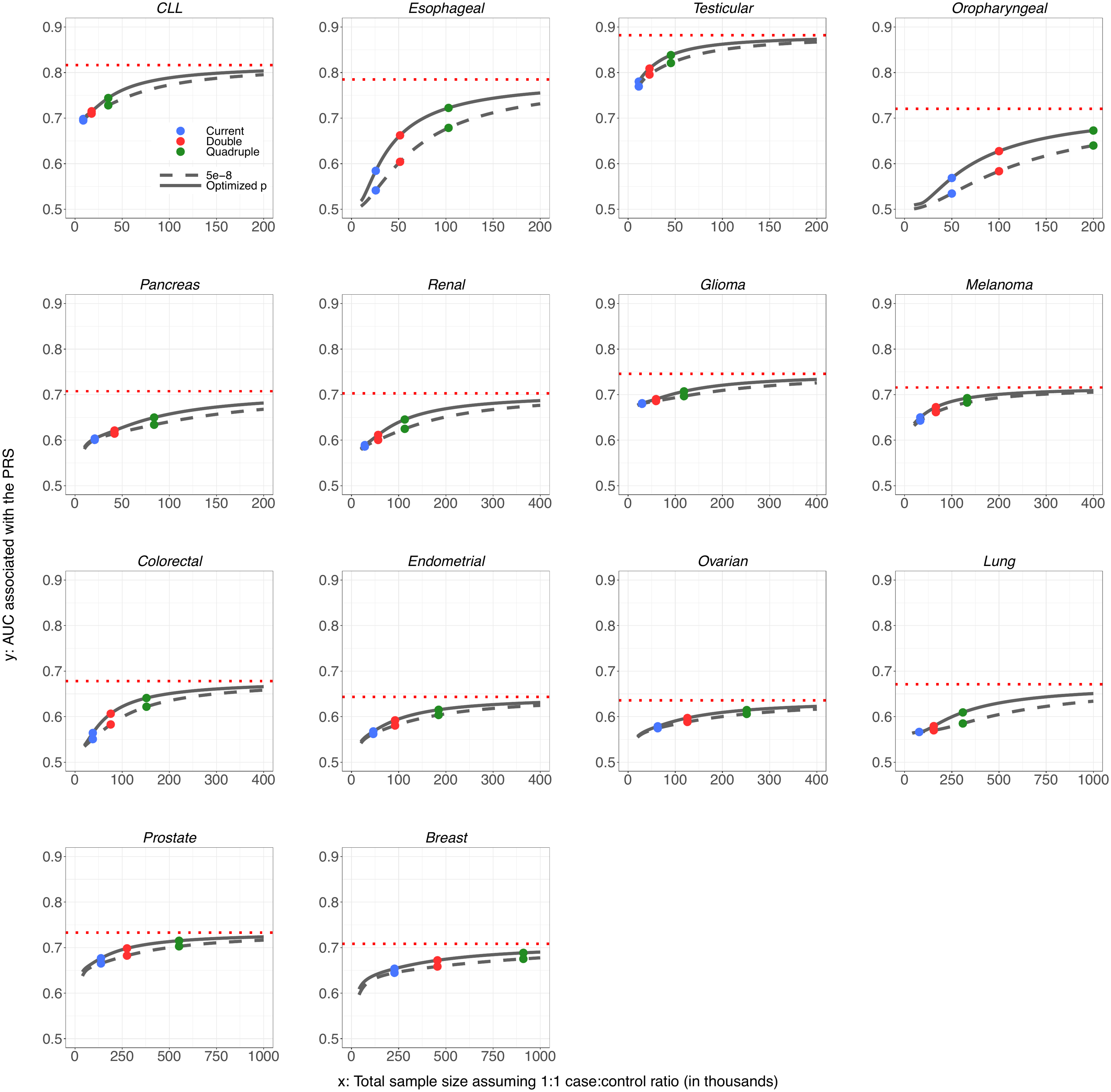

For oropharyngeal cancer, the projections at the “current sample size” are based on a sample size of 25K cases and 25K controls. For breast and esophageal cancer, the projections at the “current sample size” are based on the current largest GWAS sample sizes: 123K cases and 106K controls, and 10K cases and 17K controls, respectively. For all other cancer sites, the projections at the “current sample size” are based on the GWAS sample sizes in **Supplementary Table 1.** CLL = chronic lymphocytic leukemia.

**Supplementary Fig. 4. Projections of relative risks for individuals at or higher than 99^th^ percentile of PRS distribution (compared to average risk**) **as sample size for GWAS increases**. Results are shown for PRS including SNPs at the optimized p-value threshold (solid curve) and at genome-wide significance (P<$5\times{10}^{-8}$) level (dashed curve). The dotted horizontal red line indicates the maximum relative risk achievable according to estimate of GWAS heritability. Colored dots correspond to sample size for largest published GWAS and those for doubled and quadruped sizes.

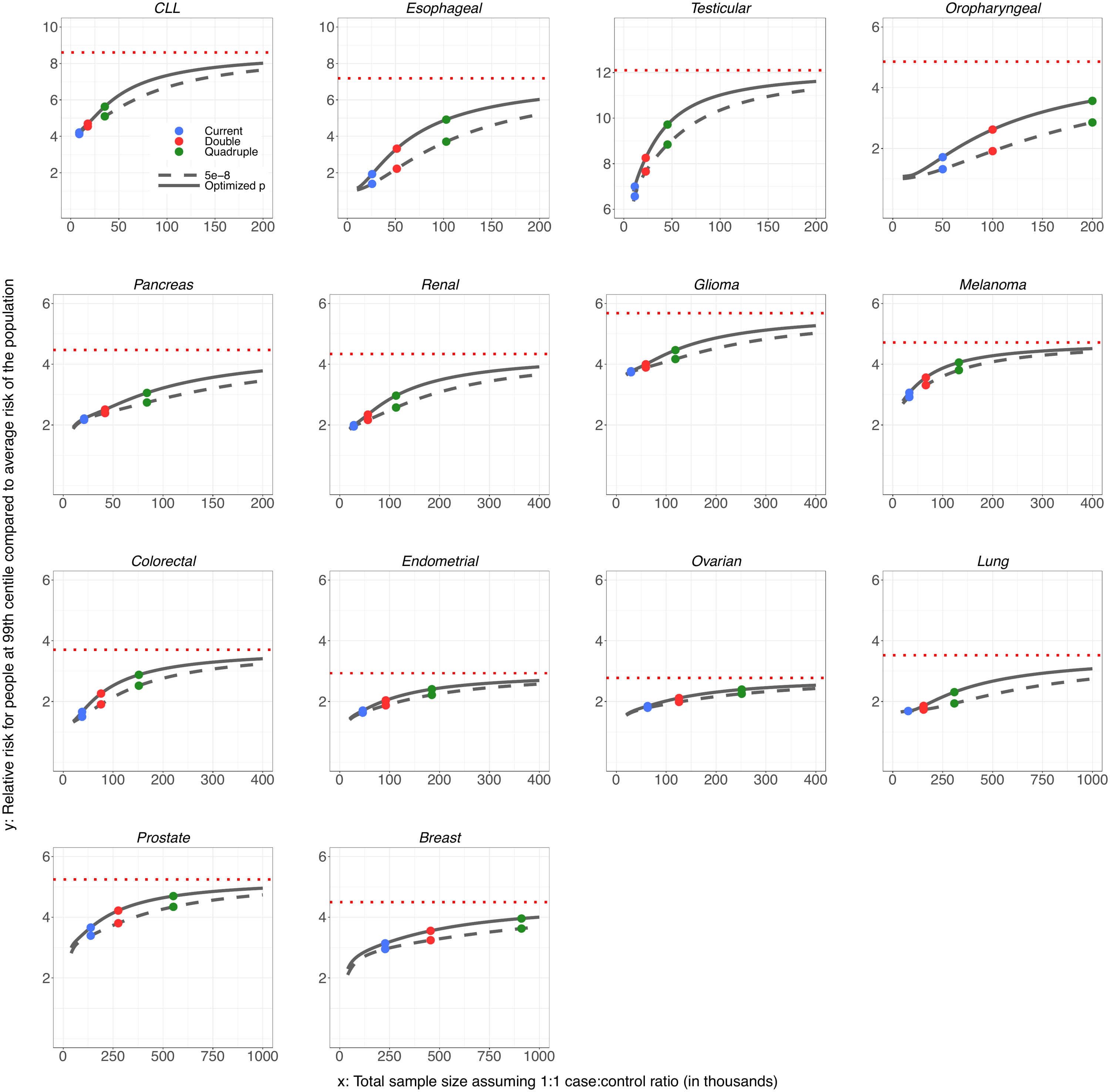

For oropharyngeal cancer, the projections at the “current sample size” are based on a sample size of 25K cases and 25K controls. For breast and esophageal cancer, the projections at the “current sample size” are based on the current largest GWAS sample sizes: 123K cases and 106K controls, and 10K cases and 17K controls, respectively. For all other cancer sites, the projections at the “current sample size” are based on the GWAS sample sizes in **Supplementary Table 1.** CLL = chronic lymphocytic leukemia.

**Supplementary Fig. 5. Projected distribution of age-stratified residual lifetime risk (up to age 75) in US Non-Hispanic Whites according to variation of polygenic risk scores (group 1 cancers).** Colored shades correspond to sample size for largest published GWAS and those for doubled, quadruped and infinite sizes.

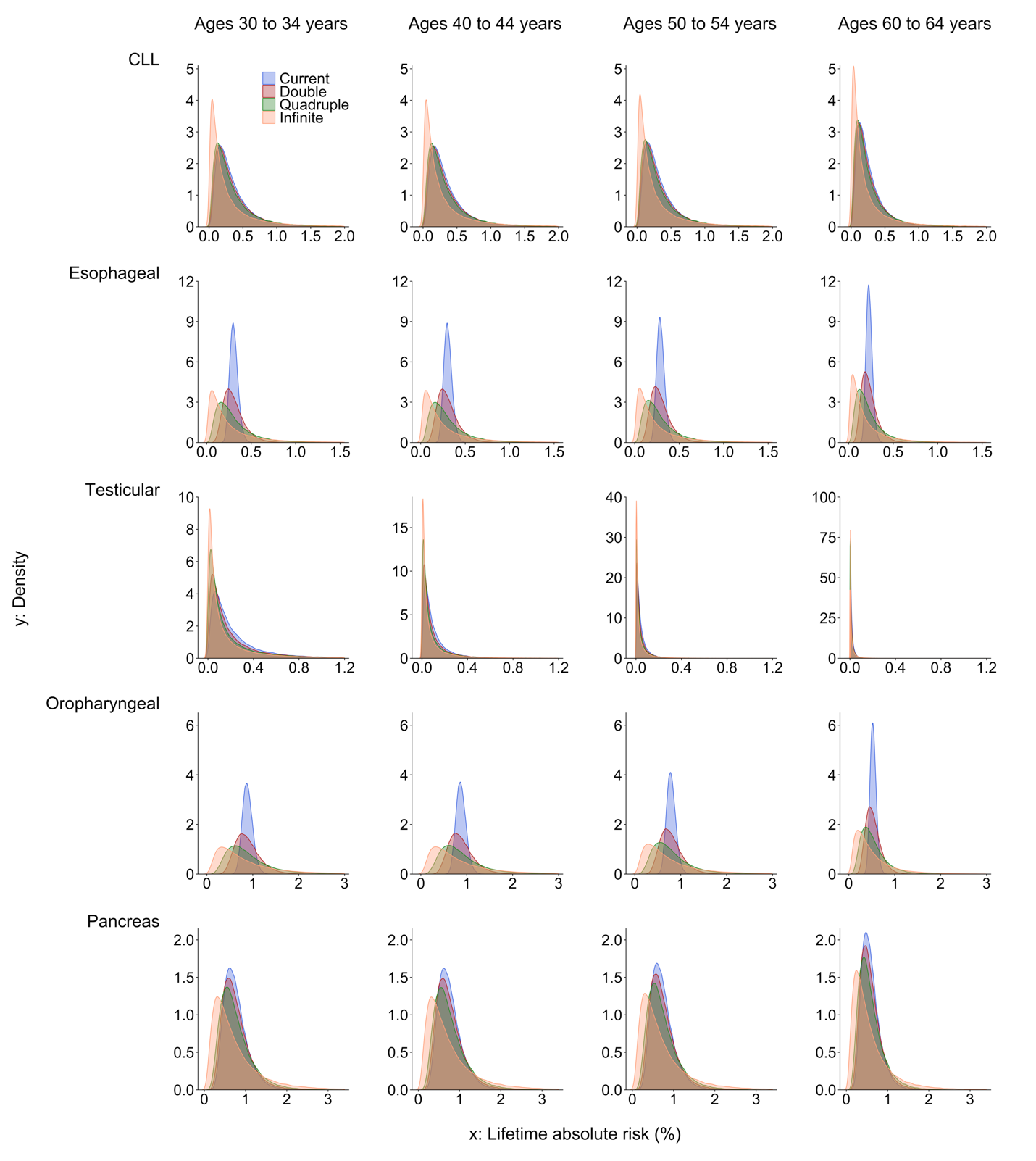

For oropharyngeal cancer, the projections at the “current sample size” are based on a sample size of 25K cases and 25K controls. For breast and esophageal cancer, the projections at the “current sample size” are based on the current largest GWAS sample sizes: 123K cases and 106K controls, and 10K cases and 17K controls, respectively. For all other cancer sites, the projections at the “current sample size” are based on the GWAS sample sizes in **Supplementary Table 1.** CLL = chronic lymphocytic leukemia.

**Supplementary Fig. 6. Projected distribution of age-stratified residual lifetime risk (up to age 75) in US Non-Hispanic Whites according to variation of polygenic risk scores (group 2 cancers).** Colored shades correspond to sample size for largest published GWAS and those for doubled, quadruped and infinite sizes.

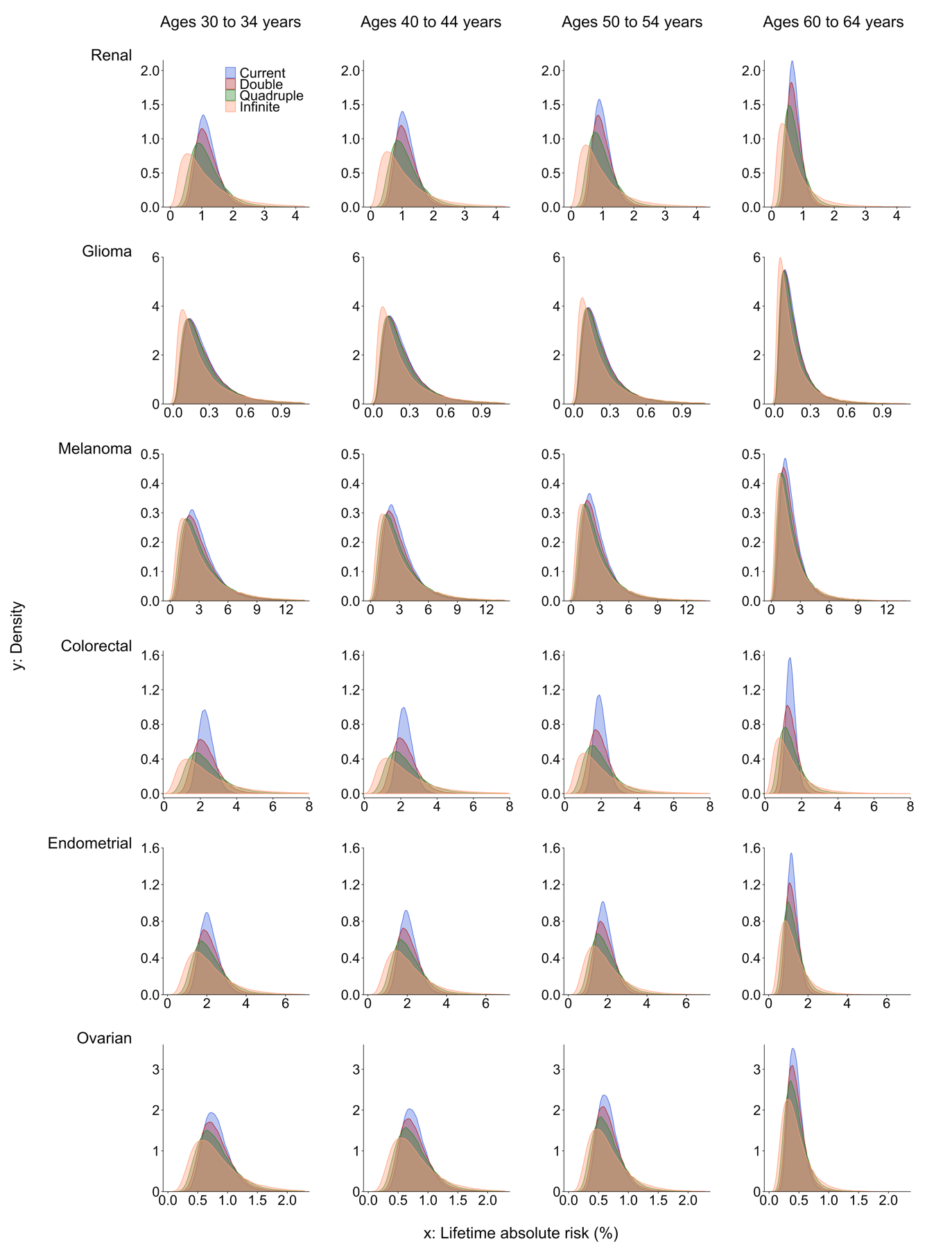

**Supplementary Fig. 7. Projected distribution of age-stratified residual lifetime risk (up to age 75) in US Non-Hispanic Whites according to variation of polygenic risk scores (group 3 cancers).** Colored shades correspond to sample size for largest published GWAS and those for doubled, quadruped and infinite sizes.

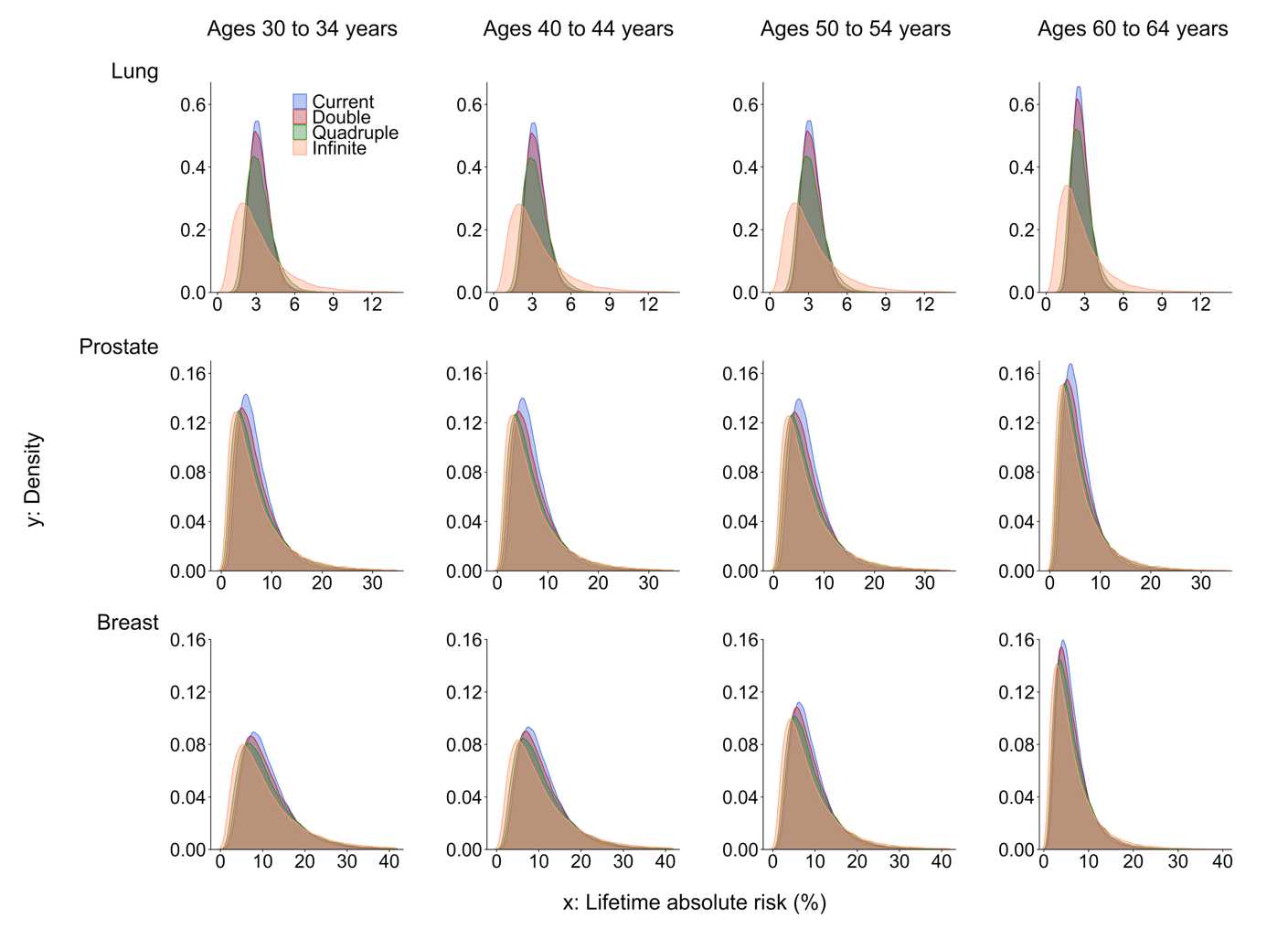

**3.0 SUPPLEMENTARY NOTE**

**3.2 CONSORTIUM-SPECIFIC FUNDING AND ACKNOWLEDGEMENTS**

**BCAC**

*Funding:*

BCAC is funded by Cancer Research UK [C1287/A16563, C1287/A10118], the European Union's Horizon 2020 Research and Innovation Programme (grant numbers 634935 and 633784 for BRIDGES and B-CAST respectively), and by the European Community´s Seventh Framework Programme under grant agreement number 223175 (grant number HEALTH-F2-2009-223175) (COGS). The EU Horizon 2020 Research and Innovation Programme funding source had no role in study design, data collection, data analysis, data interpretation or writing of the report.

Genotyping of the OncoArray was funded by the NIH Grant U19 CA148065, and Cancer UK Grant C1287/A16563 and the PERSPECTIVE project supported by the Government of Canada through Genome Canada and the Canadian Institutes of Health Research (grant GPH-129344) and, the Ministère de l’Économie, Science et Innovation du Québec through Genome Québec and the PSRSIIRI-701 grant, and the Quebec Breast Cancer Foundation. Funding for the iCOGS infrastructure came from: the European Community's Seventh Framework Programme under grant agreement n° 223175 (HEALTH-F2-2009-223175) (COGS), Cancer Research UK (C1287/A10118, C1287/A10710, C12292/A11174, C1281/A12014, C5047/A8384, C5047/A15007, C5047/A10692, C8197/A16565), the National Institutes of Health (CA128978) and Post-Cancer GWAS initiative (1U19 CA148537, 1U19 CA148065 and 1U19 CA148112 - the GAME-ON initiative), the Department of Defence (W81XWH-10-1-0341), the Canadian Institutes of Health Research (CIHR) for the CIHR Team in Familial Risks of Breast Cancer, and Komen Foundation for the Cure, the Breast Cancer Research Foundation, and the Ovarian Cancer Research Fund. The DRIVE Consortium was funded by U19 CA148065.

The **Australian Breast Cancer Family Study (ABCFS)**was supported by grant UM1 CA164920 from the National Cancer Institute (USA). The content of this manuscript does not necessarily reflect the views or policies of the National Cancer Institute or any of the collaborating centers in the Breast Cancer Family Registry (BCFR), nor does mention of trade names, commercial products, or organizations imply endorsement by the USA Government or the BCFR. The ABCFS was also supported by the National Health and Medical Research Council of Australia, the New South Wales Cancer Council, the Victorian Health Promotion Foundation (Australia) and the Victorian Breast Cancer Research Consortium. J.L.H. is a National Health and Medical Research Council (NHMRC) Senior Principal Research Fellow. M.C.S. is a NHMRC Senior Research Fellow. The **ABCS study** was supported by the Dutch Cancer Society [grants NKI 2007-3839; 2009 4363]. The **Australian Breast Cancer Tissue Bank (ABCTB)** was supported by the National Health and Medical Research Council of Australia, The Cancer Institute NSW and the National Breast Cancer Foundation. The **ACP study** is funded by the Breast Cancer Research Trust, UK. The work of the **BBCC** was partly funded by ELAN-Fond of the University Hospital of Erlangen. The **BBCS** is funded by Cancer Research UK and Breast Cancer Now and acknowledges NHS funding to the NIHR Biomedical Research Centre, and the National Cancer Research Network (NCRN). The **BCEES** was funded by the National Health and Medical Research Council, Australia and the Cancer Council Western Australia and acknowledges funding from the National Breast Cancer Foundation (JS). For the **BCFR-NY, BCFR-PA, BCFR-UT** this work was supported by grant UM1 CA164920 from the National Cancer Institute. The content of this manuscript does not necessarily reflect the views or policies of the National Cancer Institute or any of the collaborating centers in the Breast Cancer Family Registry (BCFR), nor does mention of trade names, commercial products, or organizations imply endorsement by the US Government or the BCFR. For BIGGS, ES is supported by NIHR Comprehensive Biomedical Research Centre, Guy's & St. Thomas' NHS Foundation Trust in partnership with King's College London, United Kingdom. IT is supported by the Oxford Biomedical Research Centre. BOCS is supported by funds from Cancer Research UK (C8620/A8372/A15106) and the Institute of Cancer Research (UK). **BOCS**acknowledges NHS funding to the Royal Marsden / Institute of Cancer Research NIHR Specialist Cancer Biomedical Research Centre. The **BREast Oncology GAlician Network (BREOGAN)** is funded by Acción Estratégica de Salud del Instituto de Salud Carlos III FIS PI12/02125/Cofinanciado FEDER; Acción Estratégica de Salud del Instituto de Salud Carlos III FIS Intrasalud (PI13/01136); Programa Grupos Emergentes, Cancer Genetics Unit, Instituto de Investigacion Biomedica Galicia Sur. Xerencia de Xestion Integrada de Vigo-SERGAS, Instituto de Salud Carlos III, Spain; Grant 10CSA012E, Consellería de Industria Programa Sectorial de Investigación Aplicada, PEME I + D e I + D Suma del Plan Gallego de Investigación, Desarrollo e Innovación Tecnológica de la Consellería de Industria de la Xunta de Galicia, Spain; Grant EC11-192. Fomento de la Investigación Clínica Independiente, Ministerio de Sanidad, Servicios Sociales e Igualdad, Spain; and Grant FEDER-Innterconecta. Ministerio de Economia y Competitividad, Xunta de Galicia, Spain. The **BSUCH study**was supported by the Dietmar-Hopp Foundation, the Helmholtz Society and the German Cancer Research Center (DKFZ). **CBCS** is funded by the Canadian Cancer Society (grant # 313404) and the Canadian Institutes of Health Research. **CCGP** is supported by funding from the University of Crete. The **CECILE study** was supported by Fondation de France, Institut National du Cancer (INCa), Ligue Nationale contre le Cancer, Agence Nationale de Sécurité Sanitaire, de l'Alimentation, de l'Environnement et du Travail (ANSES), Agence Nationale de la Recherche (ANR). The **CGPS** was supported by the Chief Physician Johan Boserup and Lise Boserup Fund, the Danish Medical Research Council, and Herlev and Gentofte Hospital. The American Cancer Society funds the creation, maintenance, and updating of the **CPS-II cohort**. The **CTS** was initially supported by the California Breast Cancer Act of 1993 and the California Breast Cancer Research Fund (contract 97-10500) and is currently funded through the National Institutes of Health (R01 CA77398, UM1 CA164917, and U01 CA199277). Collection of cancer incidence data was supported by the California Department of Public Health as part of the statewide cancer reporting program mandated by California Health and Safety Code Section 103885. HAC receives support from the Lon V Smith Foundation (LVS39420). The University of Westminster curates the **DietCompLyf** database funded by Against Breast Cancer Registered Charity No. 1121258 and the NCRN. The coordination of **EPIC** is financially supported by the European Commission (DG-SANCO) and the International Agency for Research on Cancer. The national cohorts are supported by: Ligue Contre le Cancer, Institut Gustave Roussy, Mutuelle Générale de l’Education Nationale, Institut National de la Santé et de la Recherche Médicale (INSERM) (France); German Cancer Aid, German Cancer Research Center (DKFZ), Federal Ministry of Education and Research (BMBF) (Germany); the Hellenic Health Foundation, the Stavros Niarchos Foundation (Greece); Associazione Italiana per la Ricerca sul Cancro-AIRC-Italy and National Research Council (Italy); Dutch Ministry of Public Health, Welfare and Sports (VWS), Netherlands Cancer Registry (NKR), LK Research Funds, Dutch Prevention Funds, Dutch ZON (Zorg Onderzoek Nederland), World Cancer Research Fund (WCRF), Statistics Netherlands (The Netherlands); Health Research Fund (FIS), PI13/00061 to Granada, PI13/01162 to EPIC-Murcia, Regional Governments of Andalucía, Asturias, Basque Country, Murcia and Navarra, ISCIII RETIC (RD06/0020) (Spain); Cancer Research UK (14136 to EPIC-Norfolk; C570/A16491 and C8221/A19170 to EPIC-Oxford), Medical Research Council (1000143 to EPIC-Norfolk, MR/M012190/1 to EPIC-Oxford) (United Kingdom). The **ESTHER study** was supported by a grant from the Baden Württemberg Ministry of Science, Research and Arts. Additional cases were recruited in the context of the VERDI study, which was supported by a grant from the German Cancer Aid (Deutsche Krebshilfe). FHRISK is funded from NIHR grant PGfAR 0707-10031. The **GC-HBOC (German Consortium of Hereditary Breast and Ovarian Cancer)** is supported by the German Cancer Aid (grant no 110837, coordinator: Rita K. Schmutzler, Cologne). This work was also funded by the European Regional Development Fund and Free State of Saxony, Germany (LIFE - Leipzig Research Centre for Civilization Diseases, project numbers 713-241202, 713-241202, 14505/2470, 14575/2470). The **GENICA** was funded by the Federal Ministry of Education and Research (BMBF) Germany grants 01KW9975/5, 01KW9976/8, 01KW9977/0 and 01KW0114, the Robert Bosch Foundation, Stuttgart, Deutsches Krebsforschungszentrum (DKFZ), Heidelberg, the Institute for Prevention and Occupational Medicine of the German Social Accident Insurance, Institute of the Ruhr University Bochum (IPA), Bochum, as well as the Department of Internal Medicine, Evangelische Kliniken Bonn gGmbH, Johanniter Krankenhaus, Bonn, Germany. The **GEPARSIXTO study** was conducted by the German Breast Group GmbH. The **GESBC** was supported by the Deutsche Krebshilfe e. V. [70492] and the German Cancer Research Center (DKFZ). The **HABCS study** was supported by the Claudia von Schilling Foundation for Breast Cancer Research, by the Lower Saxonian Cancer Society, and by the Rudolf Bartling Foundation. The**HEBCS** was financially supported by the Helsinki University Hospital Research Fund, the Finnish Cancer Society, and the Sigrid Juselius Foundation. The**HERPACC** was supported by MEXT Kakenhi (No. 170150181 and 26253041) from the Ministry of Education, Science, Sports, Culture and Technology of Japan, by a Grant-in-Aid for the Third Term Comprehensive 10-Year Strategy for Cancer Control from Ministry Health, Labour and Welfare of Japan, by Health and Labour Sciences Research Grants for Research on Applying Health Technology from Ministry Health, Labour and Welfare of Japan, by National Cancer Center Research and Development Fund, and "Practical Research for Innovative Cancer Control (15ck0106177h0001)" from Japan Agency for Medical Research and development, AMED, and Cancer Bio Bank Aichi. The **HMBCS**was supported by a grant from the Friends of Hannover Medical School and by the Rudolf Bartling Foundation. The **HUBCS**was supported by a grant from the German Federal Ministry of Research and Education (RUS08/017), and by the Russian Foundation for Basic Research and the Federal Agency for Scientific Organizations for support the Bioresource collections and RFBR grants 14-04-97088, 17-29-06014 and 17-44-020498. ICICLE was supported by Breast Cancer Now, CRUK and Biomedical Research Centre at Guy’s and St Thomas’ NHS Foundation Trust and King’s College London. Financial support for **KARBAC** was provided through the regional agreement on medical training and clinical research (ALF) between Stockholm County Council and Karolinska Institutet, the Swedish Cancer Society, The Gustav V Jubilee foundation and Bert von Kantzows foundation. The **KARMA study** was supported by Märit and Hans Rausings Initiative Against Breast Cancer. The **KBCP** was financially supported by the special Government Funding (EVO) of Kuopio University Hospital grants, Cancer Fund of North Savo, the Finnish Cancer Organizations, and by the strategic funding of the University of Eastern Finland. The **KOHBRA study** was partially supported by a grant from the Korea Health Technology R&D Project through the Korea Health Industry Development Institute (KHIDI), and the National R&D Program for Cancer Control, Ministry of Health & Welfare, Republic of Korea (HI16C1127; 1020350; 1420190). **LMBC** is supported by the 'Stichting tegen Kanker'. **DL** is supported by the FWO. The **MABCS study** is funded by the Research Centre for Genetic Engineering and Biotechnology "Georgi D. Efremov", MASA. The **MARIE study** was supported by the Deutsche Krebshilfe e.V. [70-2892-BR I, 106332, 108253, 108419, 110826, 110828], the Hamburg Cancer Society, the German Cancer Research Center (DKFZ) and the Federal Ministry of Education and Research (BMBF) Germany [01KH0402]. **MBCSG** is supported by grants from the Italian Association for Cancer Research (AIRC) and by funds from the Italian citizens who allocated the 5/1000 share of their tax payment in support of the Fondazione IRCCS Istituto Nazionale Tumori, according to Italian laws (INT-Institutional strategic projects “5x1000”). The **MCBCS** was supported by the NIH grants CA192393, CA116167, CA176785 an NIH Specialized Program of Research Excellence (SPORE) in Breast Cancer [CA116201], and the Breast Cancer Research Foundation and a generous gift from the David F. and Margaret T. Grohne Family Foundation. The **Melbourne Collaborative Cohort Study (MCCS)** cohort recruitment was funded by VicHealth and Cancer Council Victoria. The MCCS was further augmented by Australian National Health and Medical Research Council grants 209057, 396414 and 1074383 and by infrastructure provided by Cancer Council Victoria. Cases and their vital status were ascertained through the Victorian Cancer Registry and the Australian Institute of Health and Welfare, including the National Death Index and the Australian Cancer Database. The **MEC** was support by NIH grants CA63464, CA54281, CA098758, CA132839 and CA164973. The **MISS study** is supported by funding from ERC-2011-294576 Advanced grant, Swedish Cancer Society, Swedish Research Council, Local hospital funds, Berta Kamprad Foundation, Gunnar Nilsson. The **MMHS study** was supported by NIH grants CA97396, CA128931, CA116201, CA140286 and CA177150. **MSKCC** is supported by grants from the Breast Cancer Research Foundation and Robert and Kate Niehaus Clinical Cancer Genetics Initiative. The work of **MTLGEBCS** was supported by the Quebec Breast Cancer Foundation, the Canadian Institutes of Health Research for the “CIHR Team in Familial Risks of Breast Cancer” program – grant # CRN-87521 and the Ministry of Economic Development, Innovation and Export Trade – grant # PSR-SIIRI-701. **MYBRCA** is funded by research grants from the Malaysian Ministry of Higher Education (UM.C/HlR/MOHE/06) and Cancer Research Malaysia. The **NBCS** has received funding from the K.G. Jebsen Centre for Breast Cancer Research; the Research Council of Norway grant 193387/V50 (to A-L Børresen-Dale and V.N. Kristensen) and grant 193387/H10 (to A-L Børresen-Dale and V.N. Kristensen), South Eastern Norway Health Authority (grant 39346 to A-L Børresen-Dale) and the Norwegian Cancer Society (to A-L Børresen-Dale and V.N. Kristensen). The **NBHS** was supported by NIH grant R01CA100374. Biological sample preparation was conducted the Survey and Biospecimen Shared Resource, which is supported by P30 CA68485. The **Northern California Breast Cancer Family Registry (NC-BCFR)** and **Ontario Familial Breast Cancer Registry (OFBCR)** were supported by grant UM1 CA164920 from the National Cancer Institute (USA). The content of this manuscript does not necessarily reflect the views or policies of the National Cancer Institute or any of the collaborating centers in the Breast Cancer Family Registry (BCFR), nor does mention of trade names, commercial products, or organizations imply endorsement by the USA Government or the BCFR. The **NGOBCS** was supported by Grants-in-Aid for the Third Term Comprehensive Ten-Year Strategy for Cancer Control from the Ministry of Health, Labor and Welfare of Japan, and for Scientific Research on Priority Areas, 17015049 and for Scientific Research on Innovative Areas, 221S0001, from the Ministry of Education, Culture, Sports, Science, and Technology of Japan. The **NHS** was supported by NIH grants P01 CA87969, UM1 CA186107, and U19 CA148065. The **NHS2** was supported by NIH grants UM1 CA176726 and U19 CA148065. The **ORIGO study** was supported by the Dutch Cancer Society (RUL 1997-1505) and the Biobanking and Biomolecular Resources Research Infrastructure (BBMRI-NL CP16). The **PBCS** was funded by Intramural Research Funds of the National Cancer Institute, Department of Health and Human Services, USA. Genotyping for **PLCO** was supported by the Intramural Research Program of the National Institutes of Health, NCI, Division of Cancer Epidemiology and Genetics. The PLCO is supported by the Intramural Research Program of the Division of Cancer Epidemiology and Genetics and supported by contracts from the Division of Cancer Prevention, National Cancer Institute, National Institutes of Health. The **POSH study** is funded by Cancer Research UK (grants C1275/A11699, C1275/C22524, C1275/A19187, C1275/A15956 and Breast Cancer Campaign 2010PR62, 2013PR044. PROCAS is funded from NIHR grant PGfAR 0707-10031. The **RBCS** was funded by the Dutch Cancer Society (DDHK 2004-3124, DDHK 2009-4318). The SASBAC study was supported by funding from the Agency for Science, Technology and Research of Singapore (A*STAR), the US National Institute of Health (NIH) and the Susan G. Komen Breast Cancer Foundation. The **SBCGS** was supported primarily by NIH grants R01CA64277, R01CA148667, UMCA182910, and R37CA70867. Biological sample preparation was conducted the Survey and Biospecimen Shared Resource, which is supported by P30 CA68485. The scientific development and funding of this project were, in part, supported by the Genetic Associations and Mechanisms in Oncology (GAME-ON) Network U19 CA148065. **SEARCH** is funded by Cancer Research UK [C490/A10124, C490/A16561] and supported by the UK National Institute for Health Research Biomedical Research Centre at the University of Cambridge. The University of Cambridge has received salary support for **PDPP** from the NHS in the East of England through the Clinical Academic Reserve. **SEBCS** was supported by the BRL (Basic Research Laboratory) program through the National Research Foundation of Korea funded by the Ministry of Education, Science and Technology (2012-0000347). **SGBCC** is funded by the NUS start-up Grant, National University Cancer Institute Singapore (NCIS) Centre Grant and the NMRC Clinician Scientist Award. Additional controls were recruited by the Singapore Consortium of Cohort Studies-Multi-ethnic cohort (SCCS-MEC), which was funded by the Biomedical Research Council, grant number: 05/1/21/19/425. The **Sister Study (SISTER)** is supported by the Intramural Research Program of the NIH, National Institute of Environmental Health Sciences (Z01-ES044005 and Z01-ES049033). The **Two Sister Study (2SISTER)** was supported by the Intramural Research Program of the NIH, National Institute of Environmental Health Sciences (Z01-ES044005 and Z01-ES102245), and, also by a grant from Susan G. Komen for the Cure, grant FAS0703856. **SKKDKFZS** is supported by the DKFZ. The **SMC** is funded by the Swedish Cancer Foundation and the Swedish Research Council (VR 2017-00644) grant for the Swedish Infrastructure for Medical Population-based Life-course Environmental Research (SIMPLER). The **SZBCS** was supported by Grant PBZ_KBN_122/P05/2004. The **TNBCC** was supported by: a Specialized Program of Research Excellence (SPORE) in Breast Cancer (CA116201), a grant from the Breast Cancer Research Foundation, a generous gift from the David F. and Margaret T. Grohne Family Foundation. The **TWBCS** is supported by the Taiwan Biobank project of the Institute of Biomedical Sciences, Academia Sinica, Taiwan. The **UCIBCS** component of this research was supported by the NIH [CA58860, CA92044] and the Lon V Smith Foundation [LVS39420]. The **UKBGS** is funded by Breast Cancer Now and the Institute of Cancer Research (ICR), London. ICR acknowledges NHS funding to the NIHR Biomedical Research Centre. The **UKOPS** study was funded by The Eve Appeal (The Oak Foundation) and supported by the National Institute for Health Research University College London Hospitals Biomedical Research Centre. The US3SS study was supported by Massachusetts (K.M.E., R01CA47305), Wisconsin (P.A.N., R01 CA47147) and New Hampshire (L.T.-E., R01CA69664) centers, and Intramural Research Funds of the National Cancer Institute, Department of Health and Human Services, USA. The **WHI program** is funded by the National Heart, Lung, and Blood Institute, the US National Institutes of Health and the US Department of Health and Human Services (HHSN268201100046C, HHSN268201100001C, HHSN268201100002C, HHSN268201100003C, HHSN268201100004C and HHSN271201100004C). This work was also funded by NCI U19 CA148065-01.

*Acknowledgements:*

We thank all the individuals who took part in these studies and all the researchers, clinicians, technicians and administrative staff who have enabled this work to be carried out. The **COGS study** would not have been possible without the contributions of the following: Manjeet K. Bolla, Qin Wang (BCAC), Ali Amin Al Olama, Sara Benlloch (PRACTICAL), Georgia Chenevix-Trench, Antonis Antoniou, Lesley McGuffog, Fergus Couch and Ken Offit (CIMBA), Alison M. Dunning, Andrew Lee, and Ed Dicks, Craig Luccarini and the staff of the Centre for Genetic Epidemiology Laboratory, Javier Benitez, Anna Gonzalez-Neira and the staff of the CNIO genotyping unit, Daniel C. Tessier, Francois Bacot, Daniel Vincent, Sylvie LaBoissière and Frederic Robidoux and the staff of the McGill University and Génome Québec Innovation Centre, Stig E. Bojesen, Sune F. Nielsen, Borge G. Nordestgaard, and the staff of the Copenhagen DNA laboratory, and Julie M. Cunningham, Sharon A. Windebank, Christopher A. Hilker, Jeffrey Meyer and the staff of Mayo Clinic Genotyping Core Facility. **ABCFS** thank Maggie Angelakos, Judi Maskiell, Gillian Dite. **ABCS** thanks the Blood bank Sanquin, The Netherlands. **ABCTB** Investigators: Christine Clarke, Rosemary Balleine, Robert Baxter, Stephen Braye, Jane Carpenter, Jane Dahlstrom, John Forbes, Soon Lee, Debbie Marsh, Adrienne Morey, Nirmala Pathmanathan, Rodney Scott, Allan Spigelman, Nicholas Wilcken, Desmond Yip. Samples are made available to researchers on a non-exclusive basis. The **ACP study** wishes to thank the participants in the Thai Breast Cancer study. Special Thanks also go to the Thai Ministry of Public Health (MOPH), doctors and nurses who helped with the data collection process. Finally, the study would like to thank Dr Prat Boonyawongviroj, the former Permanent Secretary of MOPH and Dr Pornthep Siriwanarungsan, the Department Director-General of Disease Control who have supported the study throughout. **BBCS** thanks Eileen Williams, Elaine Ryder-Mills, Kara Sargus. **BCEES** thanks Allyson Thomson, Christobel Saunders, Terry Slevin, BreastScreen Western Australia, Elizabeth Wylie, Rachel Lloyd. The **BCINIS study** would not have been possible without the contributions of Dr. K. Landsman, Dr. N. Gronich, Dr. A. Flugelman, Dr. W. Saliba, Dr. E. Liani, Dr. I. Cohen, Dr. S. Kalet, Dr. V. Friedman, Dr. O. Barnet of the NICCC in Haifa, and all the contributing family medicine, surgery, pathology and oncology teams in all medical institutes in Northern Israel. BIGGS thanks Niall McInerney, Gabrielle Colleran, Andrew Rowan, Angela Jones. The **BREOGAN study** would not have been possible without the contributions of the following: Manuela Gago-Dominguez, Jose Esteban Castelao, Angel Carracedo, Victor Muñoz Garzón, Alejandro Novo Domínguez, Maria Elena Martinez, Sara Miranda Ponte, Carmen Redondo Marey, Maite Peña Fernández, Manuel Enguix Castelo, Maria Torres, Manuel Calaza (BREOGAN), José Antúnez, Máximo Fraga and the staff of the Department of Pathology and Biobank of the University Hospital Complex of Santiago-CHUS, Instituto de Investigación Sanitaria de Santiago, IDIS, Xerencia de Xestion Integrada de Santiago-SERGAS; Joaquín González-Carreró and the staff of the Department of Pathology and Biobank of University Hospital Complex of Vigo, Instituto de Investigacion Biomedica Galicia Sur, SERGAS, Vigo, Spain. **BSUCH** thanks Peter Bugert, Medical Faculty Mannheim. The **CAMA study** would like to recognize CONACyT for the financial support provided for this work and all physicians responsible for the project in the different participating hospitals: Dr. Germán Castelazo (IMSS, Ciudad de México, DF), Dr. Sinhué Barroso Bravo (IMSS, Ciudad de México, DF), Dr. Fernando Mainero Ratchelous (IMSS, Ciudad de México, DF), Dr. Joaquín Zarco Méndez (ISSSTE, Ciudad de México, DF), Dr. Edelmiro Pérez Rodríguez (Hospital Universitario, Monterrey, Nuevo León), Dr. Jesús Pablo Esparza Cano (IMSS, Monterrey, Nuevo León), Dr. Heriberto Fabela (IMSS, Monterrey, Nuevo León), Dr. Fausto Hernández Morales (ISSSTE, Veracruz, Veracruz), Dr. Pedro Coronel Brizio (CECAN SS, Xalapa, Veracruz) and Dr. Vicente A. Saldaña Quiroz (IMSS, Veracruz, Veracruz). **CBCS** thanks study participants, co-investigators, collaborators and staff of the Canadian Breast Cancer Study, and project coordinators Agnes Lai and Celine Morissette. **CCGP** thanks Styliani Apostolaki, Anna Margiolaki, Georgios Nintos, Maria Perraki, Georgia Saloustrou, Georgia Sevastaki, Konstantinos Pompodakis. **CGPS** thanks staff and participants of the Copenhagen General Population Study. For the excellent technical assistance: Dorthe Uldall Andersen, Maria Birna Arnadottir, Anne Bank, Dorthe Kjeldgård Hansen. The Danish Cancer Biobank is acknowledged for providing infrastructure for the collection of blood samples for the cases. CNIO-BCS thanks Guillermo Pita, Charo Alonso, Nuria Álvarez, Pilar Zamora, Primitiva Menendez, the Human Genotyping-CEGEN Unit (CNIO). **COLBCCC** thanks all patients, the physicians Justo G. Olaya, Mauricio Tawil, Lilian Torregrosa, Elias Quintero, Sebastian Quintero, Claudia Ramírez, José J. Caicedo, and Jose F. Robledo, the researchers Diana Torres, Ignacio Briceno, Fabian Gil, Angela Umana, Angela Beltran and Viviana Ariza, and the technician Michael Gilbert for their contributions and commitment to this study. Investigators from the **CPS-II cohort** thank the participants and Study Management Group for their invaluable contributions to this research. They also acknowledge the contribution to this study from central cancer registries supported through the Centers for Disease Control and Prevention National Program of Cancer Registries, as well as cancer registries supported by the National Cancer Institute Surveillance Epidemiology and End Results program. The **CTS** Steering Committee includes Leslie Bernstein, Susan Neuhausen, James Lacey, Sophia Wang, Huiyan Ma, and Jessica Clague DeHart at the Beckman Research Institute of City of Hope, Dennis Deapen, Rich Pinder, and Eunjung Lee at the University of Southern California, Pam Horn-Ross, Peggy Reynolds, Christina Clarke Dur and David Nelson at the Cancer Prevention Institute of California, Hoda Anton-Culver, Argyrios Ziogas, and Hannah Park at the University of California Irvine, and Fred Schumacher at Case Western University. **DIETCOMPLYF** thanks the patients, nurses and clinical staff involved in the study. The DietCompLyf study was funded by the charity Against Breast Cancer (Registered Charity Number 1121258) and the NCRN. We thank the participants and the investigators of **EPIC (European Prospective Investigation into Cancer and Nutrition)**. **ESTHER** thanks Hartwig Ziegler, Sonja Wolf, Volker Hermann, Christa Stegmaier, Katja Butterbach. **FHRISK** thanks NIHR for funding. **GC-HBOC** thanks Stefanie Engert, Heide Hellebrand, Sandra Kröber and LIFE - Leipzig Research Centre for Civilization Diseases (Markus Loeffler, Joachim Thiery, Matthias Nüchter, Ronny Baber). The **GENICA Network**: Dr. Margarete Fischer-Bosch-Institute of Clinical Pharmacology, Stuttgart, and University of Tübingen, Germany [HB, Wing-Yee Lo], German Cancer Consortium (DKTK) and German Cancer Research Center (DKFZ) [HB], Deutsche Forschungsgemeinschaft (DFG, German Research Foundation) under Germany's Excellence Strategy - EXC 2180 - 390900677 [HB], Department of Internal Medicine, Evangelische Kliniken Bonn gGmbH, Johanniter Krankenhaus, Bonn, Germany [Yon-Dschun Ko, Christian Baisch], Institute of Pathology, University of Bonn, Germany [Hans-Peter Fischer], Molecular Genetics of Breast Cancer, Deutsches Krebsforschungszentrum (DKFZ), Heidelberg, Germany [Ute Hamann], Institute for Prevention and Occupational Medicine of the German Social Accident Insurance, Institute of the Ruhr University Bochum (IPA), Bochum, Germany [Thomas Brüning, Beate Pesch, Sylvia Rabstein, Anne Lotz]; and Institute of Occupational Medicine and Maritime Medicine, University Medical Center Hamburg-Eppendorf, Germany [Volker Harth]. **GLACIER** thanks Kelly Kohut, Patricia Gorman, Maria Troy. **HABCS** thanks Michael Bremer. **HEBCS** thanks Sofia Khan, Johanna Kiiski, Carl Blomqvist, Kristiina Aittomäki, Rainer Fagerholm, Kirsimari Aaltonen, Karl von Smitten, Irja Erkkilä. **HKBCS** thanks Hong Kong Sanatorium and Hospital, Dr Ellen Li Charitable Foundation, The Kerry Group Kuok Foundation, National Institute of Health 1R03CA130065 and the North California Cancer Center for support. **HMBCS** thanks Peter Hillemanns, Hans Christiansen and Johann H. Karstens. **HUBCS** thanks Shamil Gantsev. ICICLE thanks Kelly Kohut, Michele Caneppele, Maria Troy. **KARMA** and **SASBAC** thank the Swedish Medical Research Counsel. **KBCP** thanks Eija Myöhänen, Helena Kemiläinen. **KConFab/AOCS** wish to thank Heather Thorne, Eveline Niedermayr, all the kConFab research nurses and staff, the heads and staff of the Family Cancer Clinics, and the Clinical Follow Up Study (which has received funding from the NHMRC, the National Breast Cancer Foundation, Cancer Australia, and the National Institute of Health (USA)) for their contributions to this resource, and the many families who contribute to kConFab. We thank all investigators of the **KOHBRA (Korean Hereditary Breast Cancer) Study**. **LAABC** thanks all the study participants and the entire data collection team, especially Annie Fung and June Yashiki. **LMBC** thanks Gilian Peuteman, Thomas Van Brussel, EvyVanderheyden and Kathleen Corthouts. **MABCS** thanks Milena Jakimovska (RCGEB “Georgi D. Efremov”), Emilija Lazarova (University Clinic of Radiotherapy and Oncology), Katerina Kubelka-Sabit, Mitko Karadjozov (Adzibadem-Sistina Hospital), Andrej Arsovski and Liljana Stojanovska (Re-Medika Hospital) for their contributions and commitment to this study. **MARIE** thanks Petra Seibold, Dieter Flesch-Janys, Judith Heinz, Nadia Obi, Alina Vrieling, Sabine Behrens, Ursula Eilber, Muhabbet Celik, Til Olchers and Stefan Nickels. **MBCSG (Milan Breast Cancer Study Group)**: Paolo Radice, Paolo Peterlongo, Siranoush Manoukian, Bernard Peissel, Roberto Villa, Cristina Zanzottera, Bernardo Bonanni, Irene Feroce, and the personnel of the Cogentech Cancer Genetic Test Laboratory. The **MCCS** was made possible by the contribution of many people, including the original investigators, the teams that recruited the participants and continue working on follow-up, and the many thousands of Melbourne residents who continue to participate in the study. We thank the coordinators, the research staff and especially the **MMHS** participants for their continued collaboration on research studies in breast cancer. **MSKCC** thanks Marina Corines, Lauren Jacobs. **MTLGEBCS** would like to thank Martine Tranchant (CHU de Québec – Université Laval Research Center), Marie-France Valois, Annie Turgeon and Lea Heguy (McGill University Health Center, Royal Victoria Hospital; McGill University) for DNA extraction, sample management and skilful technical assistance. J.S. is Chair holder of the Canada Research Chair in Oncogenetics. **MYBRCA** thanks study participants and research staff (particularly Patsy Ng, Nurhidayu Hassan, Yoon Sook-Yee, Daphne Lee, Lee Sheau Yee, Phuah Sze Yee and Norhashimah Hassan) for their contributions and commitment to this study. The following are **NBCS** Collaborators: Kristine K. Sahlberg (PhD), Lars Ottestad (MD), Rolf Kåresen (Prof. Em.) Dr. Ellen Schlichting (MD), Marit Muri Holmen (MD), Toril Sauer (MD), Vilde Haakensen (MD), Olav Engebråten (MD), Bjørn Naume (MD), Alexander Fosså (MD), Cecile E. Kiserud (MD), Kristin V. Reinertsen (MD), Åslaug Helland (MD), Margit Riis (MD), Jürgen Geisler (MD), OSBREAC and Grethe I. Grenaker Alnæs (MSc). **NBHS** and **SBCGS** thank study participants and research staff for their contributions and commitment to the studies. For **NHS** and **NHS2** the study protocol was approved by the institutional review boards of the Brigham and Women’s Hospital and Harvard T.H. Chan School of Public Health, and those of participating registries as required. We would like to thank the participants and staff of the NHS and NHS2 for their valuable contributions as well as the following state cancer registries for their help: AL, AZ, AR, CA, CO, CT, DE, FL, GA, ID, IL, IN, IA, KY, LA, ME, MD, MA, MI, NE, NH, NJ, NY, NC, ND, OH, OK, OR, PA, RI, SC, TN, TX, VA, WA, WY. The authors assume full responsibility for analyses and interpretation of these data. OBCS thanks Arja Jukkola-Vuorinen, Mervi Grip, Saila Kauppila, Meeri Otsukka, Leena Keskitalo and Kari Mononen for their contributions to this study. **OFBCR** thanks Teresa Selander, Nayana Weerasooriya. **ORIGO** thanks E. Krol-Warmerdam, and J. Blom for patient accrual, administering questionnaires, and managing clinical information. The LUMC survival data were retrieved from the Leiden hospital-based cancer registry system (ONCDOC) with the help of Dr. J. Molenaar. **PBCS** thanks Louise Brinton, Mark Sherman, Neonila Szeszenia-Dabrowska, Beata Peplonska, Witold Zatonski, Pei Chao, Michael Stagner. The ethical approval for the **POSH study** is MREC /00/6/69, UKCRN ID: 1137. We thank staff in the Experimental Cancer Medicine Centre (ECMC) supported Faculty of Medicine Tissue Bank and the Faculty of Medicine DNA Banking resource. **PREFACE** thanks Sonja Oeser and Silke Landrith. **PROCAS** thanks NIHR for funding. **RBCS** thanks Jannet Blom, Saskia Pelders, Annette Heemskerk and the Erasmus MC Family Cancer Clinic. **SBCS** thanks Sue Higham, Helen Cramp, Dan Connley, Ian Brock, Sabapathy Balasubramanian and Malcolm W.R. Reed. We thank the **SEARCH** and **EPIC** teams. **SGBCC** thanks the participants and research coordinator Ms Tan Siew Li. **SKKDKFZS** thanks all study participants, clinicians, family doctors, researchers and technicians for their contributions and commitment to this study. We thank the **SUCCESS Study** teams in Munich, Duessldorf, Erlangen and Ulm. **SZBCS** thanks Ewa Putresza. **UCIBCS** thanks Irene Masunaka. **UKBGS** thanks Breast Cancer Now and the Institute of Cancer Research for support and funding of the Breakthrough Generations Study, and the study participants, study staff, and the doctors, nurses and other health care providers and health information sources who have contributed to the study. We acknowledge NHS funding to the Royal Marsden/ICR NIHR Biomedical Research Centre. We acknowledge funding to the Manchester NIHR Biomedical Research Centre (IS-BRC-1215-20007). The authors thank the **WHI** investigators and staff for their dedication and the study participants for making the program possible.

**BEACON**

The MD Anderson controls were drawn from dbGaP (study accession: phs000187.v1.p1). Genotyping of these controls were done through the University of Texas MD Anderson Cancer Center (UTMDACC) and the Johns Hopkins University Center for Inherited Disease Research (CIDR). We acknowledge the principal investigators of this study: Christopher Amos, Qingyi Wei, and Jeff rey E Lee. Controls from the Genome-Wide Association Study of Parkinson Disease were obtained from dbGaP (study accession: phs000196.v2.p1).

Controls from the Chronic Renal Insufficiency Cohort (CRIC) were drawn from dbGaP (study accession: phs000524.v1.p1). The CRIC study was done by the CRIC investigators and supported by the National Institute of Diabetes and Digestive and Kidney Diseases (NIDDK). Data and samples from CRIC reported here were supplied by NIDDK Central Repositories. This report was not prepared in collaboration with investigators of the CRIC study and does not necessarily reflect the opinions or views of the CRIC study, the NIDDK Central Repositories, or the NIDDK.

**ECAC**

The authors thank the many individuals who participated in this study and the numerous institutions and their staff who supported recruitment.

The iCOGS and OncoArray endometrial cancer analysis were supported by NHMRC project grants [ID#1031333 & ID#1109286] Funding for the iCOGS infrastructure came from: the European Community's Seventh Framework Programme under grant agreement no 223175 [HEALTH-F2-2009-223175] [COGS], Cancer Research UK [C1287/A10118, C1287/A 10710, C12292/A11174, C1281/A12014, C5047/A8384, C5047/A15007, C5047/A10692, C8197/A16565], the National Institutes of Health [CA128978] and Post-Cancer GWAS initiative [1U19 CA148537, 1U19 CA148065 and 1U19 CA148112 - the GAME-ON initiative], the Department of Defence [W81XWH-10-1-0341], the Canadian Institutes of Health Research [CIHR] for the CIHR Team in Familial Risks of Breast Cancer, Komen Foundation for the Cure, the Breast Cancer Research Foundation, and the Ovarian Cancer Research Fund. OncoArray genotyping of ECAC cases was performed with the generous assistance of the Ovarian Cancer Association Consortium (OCAC). We particularly thank the efforts of Cathy Phelan. The OCAC OncoArray genotyping project was funded through grants from the US National Institutes of Health (CA1X01HG007491-01, U19-CA148112, R01-CA149429 and R01-CA058598); Canadian Institutes of Health Research (MOP-86727); and the Ovarian Cancer Research Fund. CIDR genotyping for the Oncoarray was conducted under contract 268201200008I. OncoArray genotyping of the BCAC controls was funded by Genome Canada Grant GPH-129344, NIH Grant U19 CA148065, and Cancer UK Grant C1287/A16563.

Stage 1 and stage 2 case genotyping was supported by the NHMRC [ID#552402, ID#1031333]. Control data were generated by the Wellcome Trust Case Control Consortium (WTCCC), and a full list of the investigators who contributed to the generation of the data is available from the WTCCC website. We acknowledge use of DNA from the British 1958 Birth Cohort collection, funded by the Medical Research Council grant G0000934 and the Wellcome Trust grant 068545/Z/02 - funding for this project was provided by the Wellcome Trust under award 085475. NSECG was supported by the EU FP7 CHIBCHA grant, Wellcome Trust Centre for Human Genetics Core Grant 090532/Z/09Z, and CORGI was funded by Cancer Research UK. We thank Nick Martin, Dale Nyholt and Anjali Henders for access to GWAS data from QIMR Controls. Recruitment of the QIMR controls was supported by the NHMRC. The University of Newcastle, the Gladys M Brawn Senior Research Fellowship scheme, The Vincent Fairfax Family Foundation, the Hunter Medical Research Institute and the Hunter Area Pathology Service all contributed towards the costs of establishing the Hunter Community Study. The WHI program is funded by the National Heart, Lung, and Blood Institute, the US National Institutes of Health and the US Department of Health and Human Services (HHSN268201100046C, HHSN268201100001C, HHSN268201100002C, HHSN268201100003C, HHSN268201100004C and HHSN271201100004C). This work was also funded by NCI U19 CA148065-01. This research has been conducted using the UK Biobank Resource under applications 5122 and 9797.

**ANECS** recruitment was supported by project grants from the NHMRC [ID#339435], The Cancer Council Queensland [ID#4196615] and Cancer Council Tasmania [ID#403031 and ID#457636]. SEARCH recruitment was funded by a programme grant from Cancer Research UK [C490/A10124]. The Bavarian Endometrial Cancer Study (BECS) was partly funded by the ELAN fund of the University of Erlangen. The Hannover-Jena Endometrial Cancer Study was partly supported by the Rudolf Bartling Foundation. The Leuven Endometrium Study (LES) was supported by the Verelst Foundation for endometrial cancer. The Mayo Endometrial Cancer Study (MECS) and Mayo controls (MAY) were supported by grants from the National Cancer Institute of United States Public Health Service [R01 CA122443, P30 CA15083, P50 CA136393, and GAME-ON the NCI Cancer Post-GWAS Initiative U19 CA148112], the Fred C and Katherine B Andersen Foundation, the Mayo Foundation, and the Ovarian Cancer Research Fund with support of the Smith family, in memory of Kathryn Sladek Smith. MoMaTEC received financial support from a Helse Vest Grant, the University of Bergen, Melzer Foundation, The Norwegian Cancer Society (Harald Andersens legat), The Research Council of Norway and Haukeland University Hospital. The Newcastle Endometrial Cancer Study (NECS) acknowledges contributions from the University of Newcastle, The NBN Children’s Cancer Research Group, Ms Jennie Thomas and the Hunter Medical Research Institute. RENDOCAS was supported through the regional agreement on medical training and clinical research (ALF) between Stockholm County Council and Karolinska Institutet [numbers: 20110222, 20110483, 20110141 and DF 07015], The Swedish Labor Market Insurance [number 100069] and The Swedish Cancer Society [number 11 0439]. The Cancer Hormone Replacement Epidemiology in Sweden Study (CAHRES, formerly called The Singapore and Swedish Breast/Endometrial Cancer Study; SASBAC) was supported by funding from the Agency for Science, Technology and Research of Singapore (A*STAR), the US National Institutes of Health and the Susan G. Komen Breast Cancer Foundation.

The **Nurses’ Health Study** (NHS) is supported by the NCI, NIH Grants Number UM1 CA186107, P01 CA087969, R01 CA49449, 1R01 CA134958, and 2R01 CA082838. The authors thank the participants and staff of the Nurses’ Health Study for their valuable contributions as well as the following state cancer registries for their help: AL, AZ, AR, CA, CO, CT, DE, FL, GA, ID, IL, IN, IA, KY, LA, ME, MD, MA, MI, NE, NH, NJ, NY, NC, ND, OH, OK, OR, PA, RI, SC, TN, TX, VA, WA, WY. The authors assume full responsibility for analyses and interpretation of these data. The authors also thank Channing Division of Network Medicine, Department of Medicine, Brigham and Women’s Hospital and Harvard Medical School. Finally, the authors also acknowledge Pati Soule and Hardeep Ranu for their laboratory assistance. The Connecticut Endometrial Cancer Study was supported by NCI, NIH Grant Number RO1CA98346. The Fred Hutchinson Cancer Research Center (FHCRC) is supported by NCI, NIH Grant Number NIH RO1 CA105212, RO1 CA 87538, RO1 CA75977, RO3 CA80636, NO1 HD23166, R35 CA39779, KO5 CA92002 and funds from the Fred Hutchinson Cancer Research Center. The Multiethnic Cohort Study (MEC) is supported by the NCI, NHI Grants Number CA54281, CA128008 and 2R01 CA082838. The California Teachers Study (CTS) is supported by NCI, NIH Grant Number 2R01 CA082838, R01 CA91019 and R01 CA77398, and contract 97-10500 from the California Breast Cancer Research Fund. The Polish Endometrial Cancer Study (PECS) is supported by the Intramural Research Program of the NCI. The Prostate, Lung, Colorectal, and Ovarian Cancer Screening Trial (PLCO) is supported by the Extramural and the Intramural Research Programs of the NCI.

**GECCO, CORECT, CCFR**

*Funding:*

**Genetics and Epidemiology of Colorectal Cancer Consortium (GECCO):** National Cancer Institute, National Institutes of Health, U.S. Department of Health and Human Services (U01 CA137088, U01 CA164930, R01201407). Genotyping/Sequencing services were provided by the Center for Inherited Disease Research (CIDR) (X01-HG008596 and X-01-HG007585). CIDR is fully funded through a federal contract from the National Institutes of Health to The Johns Hopkins University, contract number HHSN268201200008I. This research was funded in part through the NIH/NCI Cancer Center Support Grant P30 CA015704. **ASTERISK:** a Hospital Clinical Research Program (PHRC-BRD09/C) from the University Hospital Center of Nantes (CHU de Nantes) and supported by the Regional Council of Pays de la Loire, the Groupement des Entreprises Françaises dans la Lutte contre le Cancer (GEFLUC), the Association Anne de Bretagne Génétique and the Ligue Régionale Contre le Cancer (LRCC). **COLO2&3:** National Institutes of Health (R01 CA60987). **The Colon Cancer Family Registry (CFR)** Illumina GWAS was supported by funding from the National Cancer Institute, National Institutes of Health (grant numbers U01 CA122839, R01 CA143247 to G Casey). The Colon CFR/CORECT Affymetrix Axiom GWAS and OncoArray GWAS were supported by funding from National Cancer Institute, National Institutes of Health (grant number U19 CA148107 to S Gruber). The Colon CFR participant recruitment and collection of data and biospecimens used in this study were supported by the National Cancer Institute, National Institutes of Health (grant number U01 CA167551) and through cooperative agreements with the following Colon CFR centers: Australasian Colorectal Cancer Family Registry (NCI/NIH grant numbers U01 CA074778 and U01/U24 CA097735), USC Consortium Colorectal Cancer Family Registry (NCI/NIH grant numbers U01/U24 CA074799), Mayo Clinic Cooperative Family Registry for Colon Cancer Studies (NCI/NIH grant number U01/U24 CA074800), Ontario Familial Colorectal Cancer Registry (NCI/NIH grant number U01/U24 CA074783 to S Gallinger), Seattle Colorectal Cancer Family Registry (NCI/NIH grant number U01/U24 CA074794), and University of Hawaii Colorectal Cancer Family Registry (NCI/NIH grant number U01/U24 CA074806), Additional support for case ascertainment was provided from the Surveillance, Epidemiology and End Results (SEER) Program of the National Cancer Institute to Fred Hutchinson Cancer Research Center (Control Nos. N01-CN-67009 and N01-PC-35142, and Contract No. HHSN2612013000121), the Hawai‘i Department of Health (Control Nos. N01-PC- 67001 and N01-PC-35137, and Contract No. HHSN26120100037C, and the California Department of Public Health (contracts HHSN261201000035C awarded to the University of Southern California, and the following state cancer registries: AZ, CO, MN, NC, NH, and by the Victoria Cancer Registry and Ontario Cancer Registry. **Colorectal Cancer Transdisciplinary (CORECT) Study:** The CORECT Study was supported by the National Cancer Institute, National Institutes of Health (NCI/NIH), U.S. Department of Health and Human Services (grant numbers U19 CA148107, R01 CA81488, P30 CA014089, R01 CA197350,; P01 CA196569; R01 CA201407) and National Institutes of Environmental Health Sciences, National Institutes of Health (grant number T32 ES013678). **CPS-II:** The American Cancer Society funds the creation, maintenance, and updating of the Cancer Prevention Study-II (CPS-II) cohort. This study was conducted with Institutional Review Board approval. **DACHS:** This work was supported by the German Research Council (BR 1704/6-1, BR 1704/3, BR 1704/6-4, CH 117/1-1, HO 5117/2-1, HE 5998/2-1, KL 2354/3-1, RO 2270/8-1 and BR 1704/17-1), the Interdisciplinary Research Program of the National Center for Tumor Diseases (NCT), Germany, and the German Federal Ministry of Education and Research (01KH0404, 01ER0814, 01ER0815, 01ER1505A and 01ER1505B). **DALS:** National Institutes of Health (R01 CA48998 to M. L. Slattery). **Harvard cohorts (HPFS, NHS, PHS):** HPFS is supported by the National Institutes of Health (P01 CA055075, UM1 CA167552, U01 CA167552, R01 CA137178, R01 CA151993, R35 CA197735, K07 CA190673, and P50 CA127003), NHS by the National Institutes of Health (R01 CA137178, P01 CA087969, UM1 CA186107, R01 CA151993, R35 CA197735, K07 CA190673, and P50 CA127003) and PHS by the National Institutes of Health (R01 CA042182). **Kentucky:** This work was supported by the following grant support: Clinical Investigator Award from Damon Runyon Cancer Research Foundation (CI-8); NCI R01CA136726. **MCCS** cohort recruitment was funded by VicHealth and Cancer Council Victoria. The MCCS was further supported by Australian NHMRC grants 509348, 209057, 251553 and 504711 and by infrastructure provided by Cancer Council Victoria. Cases and their vital status were ascertained through the Victorian Cancer Registry (VCR) and the Australian Institute of Health and Welfare (AIHW), including the National Death Index and the Australian Cancer Database. **MEC:** National Institutes of Health (R37 CA54281, P01 CA033619, and R01 CA063464). **MECC:** This work was supported by the National Institutes of Health, U.S. Department of Health and Human Services (R01 CA81488 to SBG and GR). **NFCCR:** This work was supported by an Interdisciplinary Health Research Team award from the Canadian Institutes of Health Research (CRT 43821); the National Cancer Institute of Canada grants (18223 and 18226). The authors wish to acknowledge the contribution of Alexandre Belisle and the genotyping team of the McGill University and Génome Québec Innovation Centre, Montréal, Canada, for genotyping the Sequenom panel in the NFCCR samples. Funding was provided to Michael O. Woods by the Canadian Cancer Society Research Institute. **PMH-SCCFR:** National Cancer Institute, National Institutes of Health (grant numbers U01 CA167551, U01 CA074794 to J Potter and R01 CA076366 to P.A. Newcomb). **VITAL:** National Institutes of Health (K05 CA154337). **WHI:** The WHI program is funded by the National Heart, Lung, and Blood Institute, National Institutes of Health, U.S. Department of Health and Human Services through contracts HHSN268201100046C, HHSN268201100001C, HHSN268201100002C, HHSN268201100003C, HHSN268201100004C, and HHSN271201100004C.

*Acknowledgements:*

**ASTERISK:** We are very grateful to Dr. Bruno Buecher without whom this project would not have existed. We also thank all those who agreed to participate in this study, including the patients and the healthy control persons, as well as all the physicians, technicians and students. **CPS-II:** The authors thank the CPS-II participants and Study Management Group for their invaluable contributions to this research. The authors would also like to acknowledge the contribution to this study from central cancer registries supported through the Centers for Disease Control and Prevention National Program of Cancer Registries, and cancer registries supported by the National Cancer Institute Surveillance Epidemiology and End Results program. **DACHS:** We thank all participants and cooperating clinicians, and Ute Handte-Daub, Utz Benscheid, Muhabbet Celik and Ursula Eilber for excellent technical assistance. **Harvard cohorts (HPFS, NHS, PHS):** The study protocol was approved by the institutional review boards of the Brigham and Women’s Hospital and Harvard T.H. Chan School of Public Health, and those of participating registries as required. We would like to thank the participants and staff of the HPFS, NHS and PHS for their valuable contributions as well as the following state cancer registries for their help: AL, AZ, AR, CA, CO, CT, DE, FL, GA, ID, IL, IN, IA, KY, LA, ME, MD, MA, MI, NE, NH, NJ, NY, NC, ND, OH, OK, OR, PA, RI, SC, TN, TX, VA, WA, WY. The authors assume full responsibility for analyses and interpretation of these data. Kentucky: We would like to acknowledge the staff at the Kentucky Cancer Registry. **OFCCR:** Additional funding toward genetic analyses of OFCCR includes the Ontario Research Fund, the Canadian Institutes of Health Research, and the Ontario Institute for Cancer Research, through generous support from the Ontario Ministry of Research and Innovation. **PLCO:** The authors thank the PLCO Cancer Screening Trial screening center investigators and the staff from Information Management Services Inc and Westat Inc. Most importantly, we thank the study participants for their contributions that made this study possible. **WHI:** The authors thank the WHI investigators and staff for their dedication, and the study participants for making the program possible. A full listing of WHI investigators can be found at: <http://www.whi.org/researchers/Documents%20%20Write%20a%20Paper/WHI%20Investigator%20Short%20List.pdf>

**GenoMEL**

The melanoma meta-analysis makes use of data from two dbGap datasets (accession phs000187.v1.p1 and phs000519.v1.p1 ) **GenoMEL:** The GenoMEL study (http://www.genomel.org/) was funded by the European Commission under the 6th Framework Programme (contract no. LSHC-CT-2006-018702), by Cancer Research UK Programme Awards (C588/A4994 , C588/A10589 and C588/A19167), by a Cancer Research UK Project Grant (C8216/A6129) and by a grant from the US National Institutes of Health (NIH; CA83115). This research was also supported by the intramural Research Program of the NIH, National Cancer Institute (NCI), Division of Cancer Epidemiology and Genetics. This study makes use of data generated by the Wellcome Trust Case Control Consortium (http://www.wtccc.org.uk/). A full list of the investigators who contributed to the generation of the data is available from their website (see URLs). Funding for the project was provided by the Wellcome Trust under award 076113. Genotyping for the CIDRUK samples were provided by the Center for Inherited Disease Research (CIDR). CIDR is fully funded through a federal contract from the National Institutes of Health to The Johns Hopkins University, contract number HHSN268201200008I.

*Funding specific to particular centers is given below:*

**Stockholm:** Swedish Cancer Society, Karolinska Institutet Research Funds, Radiumhemmet Research Funds, Stockholm County Council Research Funding (ALF). **Lund:** Funding to be acknowledged; Swedish Cancer Society, Gunnar Nilsson Foundation, and European Research Council Advanced Grant (ERC-2011–294576). **Genoa:** Italian Ministry of Education, University and Research PRIN 2008, IMI and Mara Naum foundation. Italian association for cancer research (AIRC) IG 2014 (15460) to PG; IRCCS AOU San Martino-IST Istituto Nazionale per la Ricerca sul Cancro, 5% per la ricerca corrente, to PG and GBS. **Leiden:** Grant provided by European Biobanking and Biomolecular Resources Research Infrastructure (BBMRI) −Netherlands hub (CO18). **Spain:** The research at the Melanoma Unit in Barcelona is or was partially funded by Grants from Fondo de Investigaciones Sanitarias P.I. 09/01393 & 12/00840, Spain; by the CIBER de Enfermedades Raras of the Instituto de Salud Carlos III, Spain; by the AGAUR 2009 SGR 1337 and AGAUR 2014_SGR_603 of the Catalan Government, Spain; by a grant from “Fundació La Marató de TV3, 201331-30”, Catalonia, Spain; by the European Commission under the 6th Framework Programme, Contract no: LSHC-CT-2006-018702 (GenoMEL) and by the National Cancer Institute (NCI) of the US National Institute of Health (NIH) (CA83115). **Norway:** Grants from the Comprehensive Cancer Center, Oslo University Hospital (SE0728) and the Norwegian Cancer Society (71512-PR-2006-0356). **AMFS:** The AMFS was supported by the National Health and Medical Research Council of Australia (NHMRC) (project grants 566946, 107359, 211172 and program grant number 402761 to GJM and RFK); the Cancer Council New South Wales (project grant 77/00, 06/10), the Cancer Council Victoria and the Cancer Council Queensland (project grant 371); and the US National Institutes of Health (NIH RO1 grant CA-83115-01A2 and 2R01CA083115-11A1 to the international Melanoma Genetics Consortium - GenoMEL). Anne E. Cust is supported by fellowships from the Cancer Institute NSW and the NHMRC. We gratefully acknowledge all of the participants, and the work and dedication of the research coordinators, interviewers, examiners and data management staff. **WAMHS:** The WAMHS gratefully acknowledges all study participants for their time and contributions, and the Western Australian DNA Bank and the Ark at The University of Western Australia for biospecimen and bioinformatics related support. The Western Australian Cancer Registry, the WAMHS study team and the WAMHS Management Committee are also gratefully acknowledged for their assistance, as well as the Scott Kirkbride Melanoma Research Centre for funding received to establish the WAMHS resource and related salaries and PhD stipends. The Cancer Council Western Australia is also acknowledged for current salary support for Sarah Ward (Capacity Building and Collaboration grant). Genotyping services were provided by the Center for Inherited Disease Research (CIDR). CIDR is fully funded through a federal contract from the National Institutes of Health to The Johns Hopkins University, contract number HHSN268201200008I. **Q-MEGA cases and QTWINs controls (used in Q-MEGA_610k set):** Brisbane: Q-MEGA and QTWIN thanks A. Baxter, M. de Nooyer, I. Gardner, D. Statham, B. Haddon, M.J. Wright, J. Palmer, J. Symmons, B. Castellano, L. Bardsley, S. Smith, D. Smyth, L. Wallace, M.J. Campbell, A. Caracella, M. Kvaskoff, O. Zheng, B. Chapman and H. Beeby for their input in project management, sample processing and database development. We are grateful to the many research assistants and interviewers for assistance with the studies contributing to the QMEGA and QTWIN collections. The Q-MEGA/QTWIN study was supported by the Melanoma Research Alliance, the NIH NCI (CA88363, CA83115, CA122838, CA87969, CA055075, CA100264, CA133996 and CA49449), the National Health and Medical Research Council of Australia (NHMRC) (200071, 241944, 339462, 380385, 389927,389875, 389891, 389892,389938, 443036, 442915, 442981, 496610, 496675, 496739, 552485, 552498), the Cancer Councils New South Wales, Victoria and Queensland, the Cancer Institute New South Wales, the Cooperative Research Centre for Discovery of Genes for Common Human Diseases (CRC), Cerylid Biosciences (Melbourne), the Australian Cancer Research Foundation, The Wellcome Trust (WT084766/Z/08/Z) and donations from Neville and Shirley Hawkins. Stuart MacGregor acknowledges fellowship support from the Australian National Health and Medical Research Council and from an Australian Research Council Future Fellowship. **Endometriosis:** Contributors to the Endometriosis collection: Anjali K. Henders, S.H. Kennedy, S. Macgregor, N.G. Martin, S. Missmer, G.W. Montgomery, D.R. Nyholt, J.N. Painter, S.A. Treloar, L. Wallace, K.T. Zondervan. Acknowledgements: We acknowledge all the participants in the QIMR and endometriosis studies. We thank Anjali Henders, Leanne Wallace, and Lisa Bardsley for project management, sample processing and database development. We thank Endometriosis Associations for supporting study recruitment and S. Nicolaides and the Queensland Medical Laboratory for assistance with blood collection including pro bono collection and delivery of blood samples. Funding: This work was supported by the Cooperative Research Centre (CRC) for Discovery of Genes for Common. Human Diseases, Cerylid Biosciences (Melbourne), The Wellcome Trust and donations from Neville and Shirley Hawkins. Endometriosis sample genotyping was funded by a grant from the Wellcome Trust (WT084766/Z/08/Z) and NHMRC (496610). G.W.M. is supported by the NHMRC Fellowships scheme. D.R.N. was supported by the NHMRC Fellowship (613674) and Australian Research Council (ARC) Future Fellowship (FT0991022) schemes. **Study of Digestive Health:** Controls for use with the Q-MEGA_omni dataset were derived from the Study of Digestive Health group (SDH). The SDH was supported by grant number 5 RO1 CA 001833-02 from the National Cancer Institute. Its contents are solely the responsibility of the authors and do not necessarily represent the official views of the National Cancer Institute. We gratefully acknowledge the cooperation of the following institutions: Sullivan and Nicolaides Pathology (Brisbane); Queensland Medical Laboratory (Brisbane); Queensland Health Pathology Services (Brisbane); Institute of Medical and Veterinary Science (Adelaide); SouthPath (Adelaide). We also acknowledge the contribution of the study nurses and research assistants and would like to thank all of the people who participated in the study. DCW was supported by an NHMRC Research Fellowship (APP1058522). **Australian & New Zealand Registry of Advanced Glaucoma (ANZRAG)**: Support for recruitment of ANZRAG was provided by the Royal Australian and New Zealand College of Ophthalmology (RANZCO) Eye Foundation. Genotyping was funded by the National Health and Medical Research Council of Australia (#535074 and #1023911). This work was also supported by funding from the BrightFocus Foundation and a Ramaciotti Establishment Grant. The authors acknowledge the support of Ms. Bronwyn Usher‐Ridge in patient recruitment and data collection, and Dr Patrick Danoy and Dr Johanna Hadler for genotyping. **Inflammatory Bowel Disease (IBD)**: The authors express their thanks to Sullivan and Nicolaides Pathology, Queensland Medical Laboratories and the Queensland Health Pathology Service for identifying participants for this study. They are also grateful to Peter Schultz, Lauren Aoude, Loralie Parsonson, Stephen Walsh, Mitchell Stark, John Cardinal and Herlina Handoko for technical support. This research was supported by grant number CA 001833-03 from the United States National Cancer Institute. Its contents are solely the responsibility of the authors and do not necessarily represent the official views of the National Cancer Institute. PMW and DCW are Senior Research Fellows of the National Health and Medical Research Council of Australia. NP is supported by a PhD scholarship from the National Health and Medical Research Council of Australia. The funding bodies played no role in the design or conduct of the study, the collection, management, analysis, or interpretation of the data, or the preparation, review or approval of the manuscript. **M.D. Anderson:** Funding was provided by NCI grant P50CA093459, as well as by philanthropic contributions to the University of Texas MD Anderson Cancer Center Moon Shots Program, the Miriam and Jim Mulva Melanoma Research Fund and the Marit Peterson Fund for Melanoma Research. The authors thank the John Hopkins University Center for Inherited Disease Research for conducting high-throughput genotyping and University of Washington for the performance of quality control of the high-density SNP data. **MELARISK, Paris, France:** Grants from Institut National du Cancer (INCa-PL016 and INCa_5982) to F. Demenais, Ligue Nationale Contre Le Cancer (PRE 09/FD to F. Demenais and doctoral fellowship to M. Brossard), Fondation pour la Recherche Medicale (FDT20130928343), Programme Hospitalier de Recherche Clinique (AOM-07-195) to M.-F. Avril and F. Demenais. Ministère de l’Enseignement Supérieur et de la Recherche and Institut National du Cancer (INCa) to M. Lathrop. The authors thank the Epidemiological Study on the Genetics and Environment of Asthma (EGEA) cooperative group for giving access to data of the EGEA study (https://egeanet.vjf.inserm.fr). We acknowledge that the biological specimens of the French MELARISK study were obtained from the Institut Gustave Roussy and Fondation Jean Dausset–CEPH Biobanks. **Essen-Heidelberg**: The study was supported by a grant from Deutsche Forshungsgemeinschaft (GZ: SCHA 422/11-1) **Harvard:** Funding was provided by NIH grant R03CA167741. **Cambridge: SEARCH/MAPLES** We would like to thank Don Conroy, Craig Luccarini and Rebecca Mayes for their technical assistance. Funding: A.M. Dunning received funding from Cancer Research UK (Grant Numbers, C8197/A10123, C8197/A10865) and K.A. Pooley from Cancer Research UK (Grant Number C1287/A9540) and The Isaac Newton Trust. The Cambridge melanoma studies were supported by Cancer Research UK grants C490/A10124 (SEARCH) and C1287/A10118 (MAPLES). **Athens case control study:** The present work was funded by Aristeia Program. This Program is co-funded by the European Social Fund and National Resources (Project code:1094). The funders had no role in study design, data collection and analysis, decision to publish, or preparation of the manuscript. **Breakthrough Generations Study:** The Breakthrough Generations Study thank Breakthrough Breast Cancer and the Institute of Cancer Research (ICR) for support and funding of the Breakthrough Generations Study, and the Study participants, Study staff, and the doctors, nurses and other health care staff and data providers who have contributed to the Study. The ICR acknowledge NHS funding to the NIHR Biomedical Research Centre.

**SM** is supported by an Australian National Health and Medical Research Council Fellowship

**GICC**

Work at University of California, San Francisco was supported by the National Institutes of Health (grant numbers R01CA52689, P50CA097257, R01CA126831, R01CA139020, R25CA112355, and 1R01CA207360), as well as the loglio Collective, the Stanley D. Lewis and Virginia S. Lewis Endowed Chair in Brain Tumor Research, the Robert Magnin Newman Endowed Chair in Neuro-oncology, and by donations from families and friends of John Berardi, Helen Glaser, Elvera Olsen, Raymond E. Cooper, and William Martinusen. The collection of cancer incidence data used in this study was supported by the California Department of Public Health pursuant to California Health and Safety Code Section 103885; Centers for Disease Control and Prevention’s (CDC) National Program of Cancer Registries, under cooperative agreement 5NU58DP006344; the National Cancer Institute’s Surveillance, Epidemiology and End Results Program under contract HHSN261201800032I awarded to the University of California, San Francisco, contract HHSN261201800015I awarded to the University of Southern California, and contract HHSN261201800009I awarded to the Public Health Institute, Cancer Registry of Greater California. The ideas and opinions expressed herein are those of the author(s) and do not necessarily reflect the opinions of the State of California, Department of Public Health, the National Cancer Institute, and the Centers for Disease Control and Prevention or their Contractors and Subcontractors. This publication was supported by the National Center for Research Resources and the National Center for Advancing Translational Sciences, National Institutes of Health, through UCSF-CTSI Grant Number UL1 RR024131. Its contents are solely the responsibility of the authors and do not necessarily represent the official views of the NIH. The authors wish to acknowledge study participants, the clinicians and research staff at the participating medical centers, the UCSF Cancer Registry for providing vital status and other data, the UCSF Neurosurgery Tissue Bank for histo-pathology services and providing human biospecimens, and the UCSF Helen Diller Family Comprehensive Cancer Center Genome Analysis Core which is supported by a National Cancer Institute Cancer Center Support Grant (5P30CA082103). JKW was supported by the National Institutes of Health with grants R01CA207360, and P50CA097257 and by the Robert Magnin Newman Endowed Chair in Neurooncology.

**ILLCO/INTEGRAL**

NIH/NCI U19CA203654 and Cancer Prevention and Research Institute of Texas RR170048.

TRICL (U19 CA148127) and INTEGRAL (U19 CA203654).

**InterLymph**

*Support for individual studies:*

**ATBC** – The ATBC Study is supported by the Intramural Research Program of the U.S. National Cancer Institute, National Institutes of Health, and by U.S. Public Health Service contract HHSN261201500005C from the National Cancer Institute, Department of Health and Human Services. **BC** – Canadian Institutes for Health Research (CIHR); Canadian Cancer Society; Michael Smith Foundation for Health Research. **CPS-II** - The Cancer Prevention Study-II (CPS-II) Nutrition Cohort is supported by the American Cancer Society. Genotyping for all CPS-II samples were supported by the Intramural Research Program of the National Institutes of Health, NCI, Division of Cancer Epidemiology and Genetics. The authors would also like to acknowledge the contribution to this study from central cancer registries supported through the Centers for Disease Control and Prevention National Program of Cancer Registries, and cancer registries supported by the National Cancer Institute Surveillance Epidemiology and End Results program. **ELCCS** - Leukaemia & Lymphoma Research. **ENGELA** – Association pour la Recherche contre le Cancer (ARC), Institut National du Cancer (INCa), Fondation de France, Fondation contre la Leucémie, Agence nationale de sécurité sanitaire de l’alimentation, de l’environnement et du travail (ANSES). **EPIC** – Coordinated Action (Contract #006438, SP23-CT-2005-006438); HuGeF (Human Genetics Foundation), Torino, Italy; Cancer Research UK. **EpiLymph** – European Commission (grant references QLK4-CT-2000-00422 and FOOD-CT-2006-023103); the Spanish Ministry of Health (grant references CIBERESP, PI11/01810, PI14/01219, RCESP C03/09, RTICESP C03/10 and RTIC RD06/0020/0095), the Marató de TV3 Foundation (grant reference 051210), the Agència de Gestiód’AjutsUniversitarisi de Recerca – Generalitat de Catalunya (grant reference 2014SRG756) who had no role in the data collection, analysis or interpretation of the results; the NIH (contract NO1-CO-12400); the Compagnia di San Paolo—Programma Oncologia; the Federal Office for Radiation Protection grants StSch4261 and StSch4420, the José Carreras Leukemia Foundation grant DJCLS-R12/23, the German Federal Ministry for Education and Research (BMBF-01-EO-1303); the Health Research Board, Ireland and Cancer Research Ireland; Czech Republic supported by MH CZ – DRO (MMCI, 00209805) and MEYS - NPS I - LO1413; Fondation de France and Association de Recherche Contre le Cancer. **GEC/Mayo GWAS** - National Institutes of Health (CA118444, CA148690, CA92153). Intramural Research Program of the NIH, National Cancer Institute. Veterans Affairs Research Service. Data collection for Duke University was supported by a Leukemia & Lymphoma Society Career Development Award, the Bernstein Family Fund for Leukemia and Lymphoma Research, and the National Institutes of Health (K08CA134919), National Center for Advancing Translational Science (UL1 TR000135). *[HPFS acknowledgment for any GWAS project that includes HPFS data:]* **HPFS** (Walter C. Willet) – The HPFS was supported in part by National Institutes of Health grants UM1 CA167552, R01 CA149445, and R01 CA098122. We would like to thank the participants and staff of the Health Professionals Follow-up Study for their valuable contributions as well as the following state cancer registries for their help: AL, AZ, AR, CA, CO, CT, DE, FL, GA, ID, IL, IN, IA, KY, LA, ME, MD, MA, MI, NE, NH, NJ, NY, NC, ND, OH, OK, OR, PA, RI, SC, TN, TX, VA, WA, WY.  The authors assume full responsibility for analyses and interpretation of these data. The study protocol was approved by the institutional review boards of the Brigham and Women’s Hospital and Harvard T.H. Chan School of Public Health, and those of participating registries as required. **Iowa-Mayo SPORE** – NCI Specialized Programs of Research Excellence (SPORE) in Human Cancer (P50 CA97274); National Cancer Institute (P30 CA086862, P30 CA15083); Henry J. Predolin Foundation. **Italian GxE** - Italian Association for Cancer Research (AIRC, Investigator Grant 11855) (PC); Fondazione Banco di Sardegna 2010-2012, and Regione Autonoma della Sardegna (LR7 CRP-59812/2012) (MGE). **Mayo Clinic Case-Control** – National Institutes of Health (R01 CA92153); National Cancer Institute (P30 CA015083). **MCCS** – The Melbourne Collaborative Cohort Study recruitment was funded by VicHealth and Cancer Council Victoria. The MCCS was further supported by Australian NHMRC grants 209057, 251553 and 504711 and by infrastructure provided by Cancer Council Victoria. Cases and their vital status were ascertained through the Victorian Cancer Registry (VCR) and the Australian Institute of Health and Welfare (AIHW), including the National Death Index and the Australian Cancer Database. **MSKCC** – Geoffrey Beene Cancer Research Grant, Lymphoma Foundation (LF5541); Barbara K. Lipman Lymphoma Research Fund (74419); Robert and Kate Niehaus Clinical Cancer Genetics Research Initiative (57470); U01 HG007033; ENCODE; U01 HG007033. **NCI-SEER** – Intramural Research Program of the National Cancer Institute, National Institutes of Health, and Public Health Service (N01-PC-65064,N01-PC-67008, N01-PC-67009, N01-PC-67010, N02-PC-71105). **NHS** (Meir J. Stampfer) – The NHS was supported in part by National Institutes of Health grants UM1 CA186107, P01 CA87969, R01 CA49449, R01 CA149445 and R01 CA098122. We would like to thank the participants and staff of the Nurses' Health Study for their valuable contributions as well as the following state cancer registries for their help: AL, AZ, AR, CA, CO, CT, DE, FL, GA, ID, IL, IN, IA, KY, LA, ME, MD, MA, MI, NE, NH, NJ, NY, NC, ND, OH, OK, OR, PA, RI, SC, TN, TX, VA, WA, WY. The authors assume full responsibility for analyses and interpretation of these data. The study protocol was approved by the institutional review boards of the Brigham and Women’s Hospital and Harvard T.H. Chan School of Public Health, and those of participating registries as required. **NSW** - NSW was supported by grants from the Australian National Health and Medical Research Council (ID990920), the Cancer Council NSW, and the University of Sydney Faculty of Medicine. **NYU-WHS** - National Cancer Institute (R01 CA098661, P30 CA016087); National Institute of Environmental Health Sciences (ES000260). **PLCO** - This research was supported by the Intramural Research Program of the National Cancer Institute and by contracts from the Division of Cancer Prevention, National Cancer Institute, NIH, DHHS. **SCALE** – Swedish Cancer Society (2009/659). Stockholm County Council (20110209) and the Strategic Research Program in Epidemiology at Karolinska Institutet. Swedish Cancer Society grant (02 6661). National Institutes of Health (5R01 CA69669-02); Plan Denmark. **UCSF2** – The UCSF studies were supported by the NCI, National Institutes of Health, CA1046282 and CA154643. The collection of cancer incidence data used in this study was supported by the California Department of Health Services as part of the statewide cancer reporting program mandated by California Health and Safety Code Section 103885; the National Cancer Institute’s Surveillance, Epidemiology, and End Results Program under contract HHSN261201000140C awarded to the Cancer Prevention Institute of California, contract HHSN261201000035C awarded to the University of Southern California, and contract HHSN261201000034C awarded to the Public Health Institute; and the Centers for Disease Control and Prevention’s National Program of Cancer Registries, under agreement #1U58 DP000807-01 awarded to the Public Health Institute. The ideas and opinions expressed herein are those of the authors, and endorsement by the State of California, the California Department of Health Services, the National Cancer Institute, or the Centers for Disease Control and Prevention or their contractors and subcontractors is not intended nor should be inferred. **UTAH** - National Institutes of Health CA134674. Partial support for data collection at the Utah site was made possible by the Utah Population Database (UPDB) and the Utah Cancer Registry (UCR). Partial support for all datasets within the UPDB is provided by the Huntsman Cancer Institute (HCI) and the HCI Cancer Center Support grant, P30 CA42014. The UCR is supported in part by NIH contract HHSN261201000026C from the National Cancer Institute SEER Program with additional support from the Utah State Department of Health and the University of Utah. **WHI** – WHI investigators are: *Program Office* - (National Heart, Lung, and Blood Institute, Bethesda, Maryland) Jacques Rossouw, Shari Ludlam, Dale Burwen, Joan McGowan, Leslie Ford, and Nancy Geller; *Clinical Coordinating Center* - (Fred Hutchinson Cancer Research Center, Seattle, WA) Garnet Anderson, Ross Prentice, Andrea LaCroix, and Charles Kooperberg; *Investigators and Academic Centers -* (Brigham and Women's Hospital, Harvard Medical School, Boston, MA) JoAnn E. Manson; (MedStar Health Research Institute/Howard University, Washington, DC) Barbara V. Howard; (Stanford Prevention Research Center, Stanford, CA) Marcia L. Stefanick; (The Ohio State University, Columbus, OH) Rebecca Jackson; (University of Arizona, Tucson/Phoenix, AZ) Cynthia A. Thomson; (University at Buffalo, Buffalo, NY) Jean Wactawski-Wende; (University of Florida, Gainesville/Jacksonville, FL) Marian Limacher; (University of Iowa, Iowa City/Davenport, IA) Robert Wallace; (University of Pittsburgh, Pittsburgh, PA) Lewis Kuller; (Wake Forest University School of Medicine, Winston-Salem, NC) Sally Shumaker; *Women’s Health Initiative Memory Study* - (Wake Forest University School of Medicine, Winston-Salem, NC) Sally Shumaker. The WHI program is funded by the National Heart, Lung, and Blood Institute, National Institutes of Health, U.S. Department of Health and Human Services through contracts HHSN268201100046C, HHSN268201100001C, HHSN268201100002C, HHSN268201100003C, HHSN268201100004C, and HHSN271201100004C. **YALE** – National Cancer Institute (CA62006); National Cancer Institute (CA165923).

**OCAC**

*Funding:*

The Ovarian Cancer Association Consortium is supported by a grant from the Ovarian Cancer Research Fund thanks to donations by the family and friends of Kathryn Sladek Smith (PPD/RPCI.07). The scientific development and funding for this project were in part supported by the US National Cancer Institute GAME-ON Post-GWAS Initiative (U19-CA148112). This study made use of data generated by the Wellcome Trust Case Control consortium that was funded by the Wellcome Trust under award 076113. The results published here are in part based upon data generated by The Cancer Genome Atlas Pilot Project established by the National Cancer Institute and National Human Genome Research Institute (dbGap accession number phs000178.v8.p7).

The OCAC OncoArray genotyping project was funded through grants from the U.S. National Institutes of Health (CA1X01HG007491-01 (C.I.A.), U19-CA148112 (T.A.S.), R01-CA149429 (C.M.P.) and R01-CA058598 (M.T.G.); Canadian Institutes of Health Research (MOP-86727 (L.E.K.) and the Ovarian Cancer Research Fund (A.B.). The COGS project was funded through a European Commission's Seventh Framework Programme grant (agreement number 223175 - HEALTH-F2-2009-223175).

**B.M**. was supported by *grant* 17-44-020498, 17-29-06014 *of the Russian Foundation for Basic Research.*  D.P. was supported by *grant* 18-29-09129 *of the Russian Foundation for Basic Research.* ***E.K*** *was supported by*  the program for support the bioresource collections №007-030164/2 and study was performed as part of the assignment of the Ministry of Science and Higher Education of Russian Federation (№АААА-А16-116020350032-1).

*Funding for individual studies:*

**AAS:** National Institutes of Health (RO1-CA142081); **AUS:** The Australian Ovarian Cancer Study (AOCS) was supported by the U.S. Army Medical Research and Materiel Command (DAMD17-01-1-0729), National Health & Medical Research Council of Australia (199600, 400413 and 400281), Cancer Councils of New South Wales, Victoria, Queensland, South Australia and Tasmania and Cancer Foundation of Western Australia (Multi-State Applications 191, 211 and 182). AOCS gratefully acknowledges additional support from Ovarian Cancer Australia and the Peter MacCallum Foundation; **BAV:** ELAN Funds of the University of Erlangen-Nuremberg; **BEL:** National Kankerplan; **BGS:** Breast Cancer Now, Institute of Cancer Research; **BVU:** Vanderbilt University Medical Center’s BioVU is supported by the 1S10RR025141-01 instrumentation award and Vanderbilt CTSA grant from the National Institutes of Health (NIH)/National Center for Advancing Translational Sciences (NCATS) (ULTR000445); **CAM:** National Institutes of Health Research Cambridge Biomedical Research Centre and Cancer Research UK Cambridge Cancer Centre; **CHA:** Innovative Research Team in University (PCSIRT) in China (IRT1076); **CNI:** Instituto de Salud Carlos III (PI 12/01319); Ministerio de Economía y Competitividad (SAF2012); **DKE:** Ovarian Cancer Research Fund; **DOV:** National Institutes of Health R01-CA112523 and R01-CA87538; **EPC:** The coordination of EPIC is financially supported by the European Commission (DG-SANCO) and the International Agency for Research on Cancer. The national cohorts are supported by Danish Cancer Society (Denmark) (EMC 2014-6699); Ligue Contre le Cancer, Institut Gustave Roussy, Mutuelle Générale de l’Education Nationale, Institut National de la Santé et de la Recherche Médicale (INSERM) (France); German Cancer Aid, German Cancer Research Center (DKFZ), Federal Ministry of Education and Research (BMBF) (Germany); the Hellenic Health Foundation (Greece); Associazione Italiana per la Ricerca sul Cancro-AIRC-Italy and National Research Council (Italy); Dutch Ministry of Public Health, Welfare and Sports (VWS), Netherlands Cancer Registry (NKR), LK Research Funds, Dutch Prevention Funds, Dutch ZON (Zorg Onderzoek Nederland), World Cancer Research Fund (WCRF), Statistics Netherlands (The Netherlands); ERC-2009-AdG 232997 and Nordforsk, Nordic Centre of Excellence programme on Food, Nutrition and Health (Norway); Health Research Fund (FIS), PI13/00061 to Granada, PI13/01162 to EPIC-Murcia, Regional Governments of Andalucía, Asturias, Basque Country, Murcia and Navarra, ISCIII RETIC (RD06/0020) (Spain); Swedish Cancer Society, Swedish Research Council and County Councils of Skåne and Västerbotten (Sweden); Cancer Research UK (14136 to EPIC-Norfolk; C570/A16491 and C8221/A19170 to EPIC-Oxford), Medical Research Council (1000143 to EPIC-Norfolk, MR/M012190/1 to EPIC-Oxford) (United Kingdom); **GER:** German Federal Ministry of Education and Research, Programme of Clinical Biomedical Research (01 GB 9401) and the German Cancer Research Center (DKFZ); **GRC:** This research has been co-financed by the European Union (European Social Fund - ESF) and Greek national funds through the Operational Program "Education and Lifelong Learning" of the National Strategic Reference Framework (NSRF) - Research Funding Program of the General Secretariat for Research & Technology: SYN11_10_19 NBCA. Investing in knowledge society through the European Social Fund; **GRR:** Roswell Park Cancer Institute Alliance Foundation, P30 CA016056; **HAW:** U.S. National Institutes of Health (R01-CA58598, N01-CN-55424 and N01-PC-67001); **HJO:** Intramural funding; Rudolf-Bartling Foundation; **HMO:** Intramural funding; Rudolf-Bartling Foundation; **HOC:** Helsinki University Hospital Research Fund; **HOP:** University of Pittsburgh School of Medicine Dean’s Faculty Advancement Award (F. Modugno), Department of Defense (DAMD17-02-1-0669) and NCI (K07-CA080668, R01-CA95023, P50-CA159981 MO1-RR000056 R01-CA126841); **HUO:** Intramural funding; Rudolf-Bartling Foundation; **JPN:** Grant-in-Aid for the Third Term Comprehensive 10-Year Strategy for Cancer Control from the Ministry of Health, Labour and Welfare; **KRA:** This study (Ko-EVE) was supported by a grant from the Korea Health Technology R&D Project through the Korea Health Industry Development Institute (KHIDI), and the National R&D Program for Cancer Control, Ministry of Health & Welfare, Republic of Korea (HI16C1127; 0920010); **LAX:** American Cancer Society Early Detection Professorship (SIOP-06-258-01-COUN) and the National Center for Advancing Translational Sciences (NCATS), Grant UL1TR000124; **LUN:** ERC-2011-AdG 294576-risk factors cancer, Swedish Cancer Society, Swedish Research Council, Beta Kamprad Foundation; **MAC:** National Institutes of Health (R01-CA122443, P30-CA15083, P50-CA136393); Mayo Foundation; Minnesota Ovarian Cancer Alliance; Fred C. and Katherine B. Andersen Foundation; Fraternal Order of Eagles; **MAL:** Funding for this study was provided by research grant R01- CA61107 from the National Cancer Institute, Bethesda, MD, research grant 94 222 52 from the Danish Cancer Society, Copenhagen, Denmark; and the Mermaid I project; **MAS:** Malaysian Ministry of Higher Education (UM.C/HlR/MOHE/06) and Cancer Research Initiatives Foundation; **MAY:** National Institutes of Health (R01-CA122443, P30-CA15083, P50-CA136393); Mayo Foundation; Minnesota Ovarian Cancer Alliance; Fred C. and Katherine B. Andersen Foundation; **MCC:** MCCS cohort recruitment was funded by VicHealth and Cancer Council Victoria. Cancer Council Victoria, National Health and Medical Research Council of Australia (NHMRC) grants number 209057, 251533, 396414, and 504715; **MDA:** DOD Ovarian Cancer Research Program (W81XWH-07-0449); **MEC:** NIH (CA54281, CA164973, CA63464); **MOF:** Moffitt Cancer Center, Merck Pharmaceuticals, the state of Florida, Hillsborough County, and the city of Tampa; **NCO:** National Institutes of Health (R01-CA76016) and the Department of Defense (DAMD17-02-1-0666); **NEC:** National Institutes of Health R01-CA54419 and P50-CA105009 and Department of Defense W81XWH-10-1-02802; **NHS:** UM1 CA186107, P01 CA87969, R01 CA49449, R01-CA67262, UM1 CA176726; **NOR:** Helse Vest, The Norwegian Cancer Society, The Research Council of Norway; **NTH:** Radboud University Medical Centre; **OPL:** National Health and Medical Research Council (NHMRC) of Australia (APP1025142) and Brisbane Women's Club; **ORE:** OHSU Foundation; **OVA:** This work was supported by Canadian Institutes of Health Research grant (MOP-86727) and by NIH/NCI 1 R01CA160669-01A1; **PLC:** Intramural Research Program of the National Cancer Institute; **POC:** Pomeranian Medical University; **POL:** Intramural Research Program of the National Cancer Institute; **PVD:** Canadian Cancer Society and Cancer Research Society GRePEC Program; **RBH:** National Health and Medical Research Council of Australia; **RMH:** Cancer Research UK, Royal Marsden Hospital; **RPC:** National Institute of Health (P50 CA159981, R01CA126841); **SEA:** Cancer Research UK (C490/A10119 C490/A10124); UK National Institute for Health Research Biomedical Research Centres at the University of Cambridge; **SIS:** The Sister Study (SISTER) is supported by the Intramural Research Program of the NIH, National Institute of Environmental Health Sciences (Z01-ES044005 and Z01-ES049033); **SMC:** The Swedish Cancer Foundation and the Swedish Research Council (VR 2017-00644) grant for the Swedish Infrastructure for Medical Population-based Life-course Environmental Research (SIMPLER); **SRO:** Cancer Research UK (C536/A13086, C536/A6689) and Imperial Experimental Cancer Research Centre (C1312/A15589); **STA:** NIH grants U01 CA71966 and U01 CA69417; **SWH:** NIH (NCI) grant R37-CA070867; **TBO:** National Institutes of Health (R01-CA106414-A2), American Cancer Society (CRTG-00-196-01-CCE), Department of Defense (DAMD17-98-1-8659), Celma Mastery Ovarian Cancer Foundation; **TOR:** NIH grants R01 CA063678 and R01 CA063682; **UCI:** NIH R01-CA058860 and the Lon V Smith Foundation grant LVS-39420; **UHN:** Princess Margaret Cancer Centre Foundation-Bridge for the Cure; **UKO:** The UKOPS study was funded by The Eve Appeal (The Oak Foundation) and supported by the National Institute for Health Research University College London Hospitals Biomedical Research Centre; **UKR:** Cancer Research UK (C490/A6187), UK National Institute for Health Research Biomedical Research Centres at the University of Cambridge; **USC:** P01CA17054, P30CA14089, R01CA61132, N01PC67010, R03CA113148, R03CA115195, N01CN025403, and California Cancer Research Program (00-01389V-20170, 2II0200); **VAN:** BC Cancer Foundation, VGH & UBC Hospital Foundation; **VTL:** NIH K05-CA154337; **WMH:** National Health and Medical Research Council of Australia, Enabling Grants ID 310670 & ID 628903. Cancer Institute NSW Grants 12/RIG/1-17 & 15/RIG/1-16; **WOC:** National Science Centre (N N301 5645 40), The Maria Sklodowska-Curie Memorial Cancer Centre and Institute of Oncology, Warsaw, Poland.

*Acknowledgements:*

We are grateful to the family and friends of Kathryn Sladek Smith for their generous support of the Ovarian Cancer Association Consortium through their donations to the Ovarian Cancer Research Fund. The OncoArray and COGS genotyping projects would not have been possible without the contributions of the following: Per Hall (COGS); Kyriaki Michailidou, Manjeet K. Bolla, Qin Wang (BCAC), (OCAC), Rosalind A. Eeles, Ali Amin Al Olama, Zsofia Kote-Jarai, Sara Benlloch (PRACTICAL), Antonis Antoniou, Lesley McGuffog, Fergus Couch and Ken Offit (CIMBA), Alison M. Dunning, Andrew Lee, and Ed Dicks, Craig Luccarini and the staff of the Centre for Genetic Epidemiology Laboratory, Anna Gonzalez-Neira and the staff of the CNIO genotyping unit, Jacques Simard and Daniel C. Tessier, Francois Bacot, Daniel Vincent, Sylvie LaBoissière and Frederic Robidoux and the staff of the McGill University and Génome Québec Innovation Centre, Stig E. Bojesen, Sune F. Nielsen, Borge G. Nordestgaard, and the staff of the Copenhagen DNA laboratory, and Julie M. Cunningham, Sharon A. Windebank, Christopher A. Hilker, Jeffrey Meyer and the staff of Mayo Clinic Genotyping Core Facility. We pay special tribute to the contribution of Professor Brian Henderson to the GAME-ON consortium; to Olga M. Sinilnikova for her contribution to CIMBA and for her part in the initiation and coordination of GEMO until she sadly passed away on 30th June 2014 and to Catherine M. Phelan for her contribution to OCAC and the coordination of the OncoArray until she passed away on 22 September 2017. We thank the study participants, doctors, nurses, clinical and scientific collaborators, health care providers and health information sources who have contributed to the many studies contributing to this manuscript.

*Acknowledgements for individual studies:*

**AUS:** The AOCS also acknowledges the cooperation of the participating institutions in Australia, and the contribution of the study nurses, research assistants and all clinical and scientific collaborators. The complete AOCS Study Group can be found at [www.aocstudy.org](http://www.aocstudy.org). We would like to thank all of the women who participated in this research program; **BEL:** We would like to thank Gilian Peuteman, Thomas Van Brussel, Annick Van den Broeck and Joke De Roover for technical assistance; **BGS:** The Institute of Cancer Research (ICR) acknowledges NHS funding to the NIHR Biomedical Research Centre. We thank the Study staff, study participants , doctors, nurses, health care providers and health information sources who have contributed to the study; **BVU:** The dataset(s) used for the analyses described were obtained from Vanderbilt University Medical Center’s BioVU; **CHA:** Innovative Research Team in University (PCSIRT) in China (IRT1076); **CHN:** To thank all members of Department of Obstetrics and Gynaecology, Hebei Medical University, Fourth Hospital and Department of Molecular Biology, Hebei Medical University, Fourth Hospital; **EPC:** To thank all members and investigators of the Rotterdam Ovarian Cancer Study; **GER:** The German Ovarian Cancer Study (GER) thanks Ursula Eilber for competent technical assistance; **MAS:** We would like to thank Famida Zulkifli and Ms Moey for assistance in patient recruitment, data collection and sample preparation; **MCC:** Cases and their vital status were ascertained through the Victorian Cancer Registry (VCR) and the Australian Institute of Health and Welfare (AIHW), including the National Death Index and the Australian Cancer Database; **MOF:** the Total Cancer Care™ Protocol and the Collaborative Data Services and Tissue Core Facilities at the H. Lee Moffitt Cancer Center & Research Institute, an NCI designated Comprehensive Cancer Center (P30-CA076292), Merck Pharmaceuticals and the state of Florida; **NHS:** The NHS/NHSII studies thank the following state cancer registries for their help: AL, AZ, AR, CA, CO, CT, DE, FL, GA, ID, IL, IN, IA, KY, LA, ME, MD, MA, MI, NE, NH, NJ, NY, NC, ND, OH, OK, OR, PA, RI, SC, TN, TX, VA, WA, and WY; **OPL:** Members of the OPAL Study Group (<http://opalstudy.qimrberghofer.edu.au/>); **SEA:** SEARCH team, Craig Luccarini, Caroline Baynes, Don Conroy; **SRO:** To thank all members of Scottish Gynaecological Clinical Trails group and SCOTROC1 investigators; **SWH:** We thank the participants and the research staff of the Shanghai Women’s Health Study for making this study possible; **UHN:** Princess Margaret Cancer Centre Foundation-Bridge for the Cure; **UKO:** We particularly thank I. Jacobs, M.Widschwendter, E. Wozniak, A. Ryan, J. Ford and N. Balogun for their contribution to the study**; UKR:** Carole Pye; **VAN:** BC Cancer Foundation, VGH & UBC Hospital Foundation; **WMH:** We thank the Gynaecological Oncology Biobank at Westmead, a member of the Australasian Biospecimen Network-Oncology group.

**Oral Cancer GWAS**

Oral cancer GWAS Genotyping performed at the Center for Inherited Disease Research (CIDR) was funded through US National Institute of Dental and Craniofacial Research (NIDCR) grant 1X01HG007780-0.

**PANC4**

This work was supported by RO1 CA 154823. Genotyping Services were provided by the Center for Inherited Disease Research (CIDR).  CIDR is fully funded through a federal contract from the National Institutes of Health to the Johns Hopkins University, contract number HHSN268201100011I.

The **IARC/Central Europe study** was supported by a grant from the US National Cancer Institute at the National Institutes of Health (R03 CA123546-02) and grants from the Ministry of Health of the Czech Republic (NR 9029-4/2006, NR9422-3, NR9998-3, MH CZ-DRO-MMCI 00209805). The work at **Johns Hopkins University** was supported by the NCI Grants P50CA062924 and R01CA97075. The **Mayo Clinic Biospecimen Resource for Pancreas Research study** is supported by the Mayo Clinic SPORE in Pancreatic Cancer (P50 CA102701). The **Memorial Sloan Kettering Cancer Center Pancreatic Tumor Registry** is supported by P30CA008748, the Geoffrey Beene Foundation, the Arnold and Arlene Goldstein Family Foundation, and the Society of MSKCC. The **Queensland Pancreatic Cancer Study** was supported by a grant from the National Health and Medical Research Council of Australia (NHMRC) (Grant number 442302). RE Neale is supported by a NHMRC Senior Research Fellowship (#1060183). **SELECT:** Research reported in this publication was supported in part by the National Cancer Institute of the National Institutes of Health under Award Numbers U10 CA37429 (CD Blanke), and UM1 CA182883 (CM Tangen/IM Thompson). The content is solely the responsibility of the authors and does not necessarily represent the official views of the National Institutes of Health. The **UCSF pancreas study** was supported by NIH-NCI grants (R01CA1009767, R01CA109767-S1) and the Joan Rombauer Pancreatic Cancer Fund. Collection of cancer incidence data was supported by the California Department of Public Health as part of the statewide cancer reporting program; the NCI’s SEER Program under contract HHSN261201000140C awarded to CPIC; and the CDC’s National Program of Cancer Registries, under agreement #U58DP003862-01 awarded to the California Department of Public Health. The **Yale (CT) pancreas study** is supported by National Cancer Institute at the U.S. National Institutes of Health, grant 5R01CA098870. The cooperation of 30 Connecticut hospitals, including Stamford Hospital, in allowing patient access, is gratefully acknowledged.  The Connecticut Pancreas Cancer Study was approved by the State of Connecticut Department of Public Health Human Investigation Committee.  Certain data used in that study were obtained from the Connecticut Tumor Registry in the Connecticut Department of Public Health. The authors assume full responsibility for analyses and interpretation of these data.

**PanScan**

The work conducted at **NCI** was supported by the Intramural Research Program (IRP) of the Division of Cancer Epidemiology and Genetics, National Cancer Institute, US National Institutes of Health (NIH). The work conducted at the **Vanderbilt University Medical Center** was supported in part by R01CA188214 and K99CA218892. The **Melbourne Collaborative Cohort Study** cohort recruitment was funded by VicHealth and Cancer Council Victoria. The MCCS was further augmented by Australian National Health and Medical Research Council grants 209057, 396414 and 1074383 and by infrastructure provided by Cancer Council Victoria. Cases and their vital status were ascertained through the Victorian Cancer Registry and the Australian Institute of Health and Welfare, including the National Death Index and the Australian Cancer Database. The **WHI** program is funded by the National Heart, Lung, and Blood Institute, National Institutes of Health, U.S. Department of Health and Human Services through contracts HHSN268201600018C, HHSN268201600001C, HHSN268201600002C, HHSN268201600003C, and HHSN268201600004C. For a list of WHI investigators contributed to WHI science, please visit: <https://www.whi.org/researchers/SitePages/Principal%20Investigators.aspx>. Cancer incidence data for **CLUE** were provided by the Maryland Cancer Registry, Center for Cancer Surveillance and Control, Department of Health and Mental Hygiene, 201 W. Preston Street, Room 400, Baltimore, MD 21201, http://phpa.dhmh.maryland.gov/cancer, 410-767-4055. We acknowledge the State of Maryland, the Maryland Cigarette Restitution Fund, and the National Program of Cancer Registries of the Centers for Disease Control and Prevention for the funds that support the collection and availability of the cancer registry data.” We thank all the CLUE participants. The **NYU study** was funded by NIH R01 CA098661, UM1 CA182934 and center grants P30 CA016087 and P30 ES000260. The **Physicians' Health Study** was supported by research grants CA-097193, CA-34944, CA40360, HL-26490, and HL-34595 from the National Institutes of Health, Bethesda, MD USA. The **Women's Health Study** was supported by research grants CA-047988, HL-043851, HL080467, and HL-099355 from the National Institutes of Health, Bethesda, MD USA. **Health Professionals Follow-up Study** is supported by NIH grant UM1 CA167552 from the National Cancer Institute, Bethesda, MD USA. **Nurses’ Health Study** is supported by NIH grants UM1 CA186107, and R01 CA49449 from the National Cancer Institute, Bethesda, MD USA. The **PANKRAS II Study** in Spain was supported by research grants from Instituto de Salud Carlos III-FEDER, Spain: Fondo de Investigaciones Sanitarias (FIS) ((#PI95/0017, #PI12/00815, #PI13/00082 and #PI15/01573), Red Temática de Investigación Cooperativa en Cáncer (#RD12/0036/0050), and CIBER de Epidemiología (CIBERESP); Ministerio de Ciencia y Tecnología (CICYT SAF 2000-0097); Generalitat de Catalunya (CIRIT - SGR), Spain. The **IARC/Central Europe study** was supported by a grant from the US National Cancer

Institute at the National Institutes of Health (R03 CA123546-02) and grants from the Ministry of Health of the Czech Republic (NR 9029-4/2006, NR9422-3, NR9998-3, MH CZDRO-MMCI 00209805).

**PRACTICAL, CRUK, BPC3, CAPS, PEGASUS**

**CRUK and PRACTICAL consortium:**

This work was supported by the Canadian Institutes of Health Research, European Commission's Seventh Framework Programme grant agreement n° 223175 (HEALTH-F2-2009-223175), Cancer Research UK Grants C5047/A7357, C1287/A10118, C1287/A16563, C5047/A3354, C5047/A10692, C16913/A6135, and The National Institute of Health (NIH) Cancer Post-Cancer GWAS initiative grant: No. 1 U19 CA 148537-01 (the GAME-ON initiative).

We would also like to thank the following for funding support: The Institute of Cancer Research and The Everyman Campaign, The Prostate Cancer Research Foundation, Prostate Research Campaign UK (now PCUK), The Orchid Cancer Appeal, Rosetrees Trust, The National Cancer Research Network UK, The National Cancer Research Institute (NCRI) UK. We are grateful for support of NIHR funding to the NIHR Biomedical Research Centre at The Institute of Cancer Research and The Royal Marsden NHS Foundation Trust.

The Prostate Cancer Program of Cancer Council Victoria also acknowledge grant support from The National Health and Medical Research Council, Australia (126402, 209057, 251533, , 396414, 450104, 504700, 504702, 504715, 623204, 940394, 614296,), VicHealth, Cancer Council Victoria, The Prostate Cancer Foundation of Australia, The Whitten Foundation, PricewaterhouseCoopers, and Tattersall’s. EAO, DMK, and EMK acknowledge the Intramural Program of the National Human Genome Research Institute for their support.

Genotyping of the OncoArray was funded by the US National Institutes of Health (NIH) [U19 CA 148537 for ELucidating Loci Involved in Prostate cancer SuscEptibility (ELLIPSE) project and X01HG007492 to the Center for Inherited Disease Research (CIDR) under contract number HHSN268201200008I]. Additional analytic support was provided by NIH NCI U01 CA188392 (PI: Schumacher).

Funding for the iCOGS infrastructure came from: the European Community's Seventh Framework Programme under grant agreement n° 223175 (HEALTH-F2-2009-223175) (COGS), Cancer Research UK (C1287/A10118, C1287/A 10710, C12292/A11174, C1281/A12014, C5047/A8384, C5047/A15007, C5047/A10692, C8197/A16565), the National Institutes of Health (CA128978) and Post-Cancer GWAS initiative (1U19 CA148537, 1U19 CA148065 and 1U19 CA148112 - the GAME-ON initiative), the Department of Defence (W81XWH-10-1-0341), the Canadian Institutes of Health Research (CIHR) for the CIHR Team in Familial Risks of Breast Cancer, Komen Foundation for the Cure, the Breast Cancer Research Foundation, and the Ovarian Cancer Research Fund.

**BPC3:** The BPC3 was supported by the U.S. National Institutes of Health, National Cancer Institute (cooperative agreements U01-CA98233 to D.J.H., U01-CA98710 to S.M.G., U01-CA98216 toE.R., and U01-CA98758 to B.E.H., and Intramural Research Program of NIH/National Cancer Institute, Division of Cancer Epidemiology and Genetics).

**CAPS:** CAPS GWAS study was supported by the Cancer Risk Prediction Center (CRisP; www.crispcenter.org), a Linneus Centre (Contract ID 70867902) financed by the Swedish Research Council, (grant no K2010-70X-20430-04-3), the Swedish Cancer Foundation (grant no 09-0677), the Hedlund Foundation, the Soederberg Foundation, the Enqvist Foundation, ALF funds from the Stockholm County Council. Stiftelsen Johanna Hagstrand och Sigfrid Linner's Minne, Karlsson's Fund for urological and surgical research.

**PEGASUS:** PEGASUS was supported by the Intramural Research Program, Division of Cancer Epidemiology and Genetics, National Cancer Institute, National Institutes of Health.

**Renal Cancer GWAS**

**AHS:** This research was supported by the Intramural Research Program of the NIH, National Cancer

Institute, Division of Cancer Epidemiology and Genetics (Z01CP010119).**ATBC:** The ATBC Study was supported by funding provided by the Intramural Research Program of the NCI, NIH, and through U.S. Public Health Service contracts (N01-CN-45165, N01-RC-45035 and N01-RC-37004) from the NCI. **BioVU:** The dataset used in the analyses described were obtained from the Vanderbilt University Medical

Center resource BioVU, which is supported by institutional funding, the 1S10RR025141-01

instrumentation award, and by the Vanderbilt CTSA grant UL1 TR000445 from NCATS/NIH.

Sample processing and phenotyping algorithm development was supported by institutional

funding for TLE. **Center ‘Bioengineering’ of the Russian Academy of Sciences/Kurchatov Scientific Center:** The work conducted for this study was supported by the grant Russian Scientific Fund 14-14-

01202. **Centre National de Genotypage, France:** We thank Jean Guillaume Garnier and Delphine Bacq-Daian for their work on the IARC-2 scan. **ConFIRM/MCCS:** The ConFIRM study, also known as CARES, was supported by the Victorian Cancer Agency (PTCB08_05), the Australian National Health and Medical Research Council (Project Grant 1011626). We acknowledge the contribution of Professor Graham Giles in supporting this work and of Ms Olive Schmid and Ms Jennifer Walsh for the project management. The Melbourne Collaborative Cohort Study (MCCS) recruitment was funded by VicHealth and Cancer Council Victoria. The MCCS was further supported by Australian NHMRC grants 209057, 251553 and 504711 and by infrastructure provided by Cancer Council Victoria. **CPS-II:** The Cancer Prevention Study II Nutrition Cohort is supported by the American Cancer Society. The authors thank all of the men and women in the Cancer Prevention Study II Nutrition Cohort for their many years of dedicated participation in the study. **Health Professionals Follow-up Study (HPFS) and Nurses’ Health Study (NHS):** The HPFS is supported by National Institutes of Health, National Cancer Institute grant UM1CA167552. The NHS is supported by National Institutes of Health, National Cancer Institute grants UM1 CA186107 and P01 CA87969. We would like to thank the following state cancer registries for their help: AL, AZ, AR, CA, CO, CT, DE, FL, GA, ID, IL, IN, IA, KY, LA, ME, MD, MA, MI, NE, NH, NJ, NY, NC, ND, OH, OK, OR, PA, RI, SC, TN, TX, VA, WA, WY. The authors assume full responsibility for analyses and interpretation of these data. **K2 study:** This study was supported by the EU FP7 under grant agreement number 241669 (the CAGEKID project). In Czech Republic, this work was also supported by MH CZ—DRO (MMCI, 00209805) and by the project MEYS – NPS I – LO1413, Czech Republic. **Leeds Cohort:** The infrastructure support from Cancer Research UK as part of the Leeds Centre and Experimental Cancer Medicine Centre funding is gratefully acknowledged together with the Leeds Multidisciplinary Research Tissue Bank, all patients who consented to take part in the research studies and the staff of the Urology and Oncology Departments in Leeds Teaching Hospitals Trust. **Mayo Clinic:** This study was partially supported by National Institutes of Health: R21CA176422 (JEEP) and R01CA134466 (ASP). The authors acknowledge the Mayo Clinic Comprehensive Cancer Center Biospecimens Accessioning and Processing Shared Resource and the Pathology Research Core Shared Resource. **MD Anderson:** This work was supported in part by the NIH (grant R01 CA170298) and the Center for Translational and Public Health Genomics, Duncan Family Institute for Cancer Prevention and Risk Assessment, The University of Texas MD Anderson Cancer Center. **NCI/IARC RCC Study in Central Europe (CE):** This project was supported by the Intramural Research Program of the NIH and the National Cancer Institute. **Physicians’ Health Study (PHS):** This study was supported by grants CA 097193, CA 34944, CA 40360, HL 26490, and HL 34595 from the National Institutes of Health, Bethesda, MD. **PLCO:** This research was supported by the Intramural Research Program of the National Cancer Institute and by contracts from the Division of Cancer Prevention, National Cancer Institute, NIH, DHHS. **SEARCH:** This study is funded by Cancer Research UK (C490/A16651). **UK GWAS:** We acknowledge support from the Medical Research Council (MRC), Cancer Research UK, an educational grant from Bayer and NHS funding for the Royal Marsden Biomedical Research Centre and Cambridge University Health Partners. JL is supported by the NIHR RM/CR

Biomedical Research Centre for Cancer. **US Kidney Cancer Study:** The NCI United Stated Kidney Cancer Study was supported by the Intramural Research Program of the National Institutes of Health and the National Cancer Institute under the following contracts: N02-CP-10128 (Westat, Inc.), N02-CP-11004 (Wayne State University), and N02-CP- 11161 (University of Illinois at Chicago). **Van Andel Research Institute (VARI):** The authors would like to thank Dr. Kyle Furge for his role in this project as well as Drs. Anthony Avallone, John Ludlow and Philip Wise for contributing samples for this project. **Women’s Health Initiative (WHI):** The WHI program is funded by the National Heart, Lung, and Blood Institute, National Institutes of Health, U.S. Department of Health and Human Services through contracts HHSN268201100046C, HHSN268201100001C, HHSN268201100002C, HHSN268201100003C, HHSN268201100004C, and HHSN271201100004C. For a list of all the investigators who have contributed to WHI science, please visit: <https://www.whi.org/researchers/SitePages/Principal%20Investigators.aspx>. **Women’s Health Study (WHS):** The study is supported by grants CA-047988, HL-043851, HL-080467, HL-099355, and UM1CA182913 from the National Institutes of Health, Bethesda, MD.

**TECAC**

This work was supported by the Testicular Cancer Consortium (TECAC) funded by the National Cancer Institute under Grant no. R01 CA164947. The TECAC is an international collaborative effort between the National Cancer Institute (USA), The University of Pennsylvania (USA), Copenhagen University Hospital, Rigshospitalet (Denmark), Spanish National Cancer Research Center (Spain), University of Leeds (UK), University of Southern California (USA), University of Brescia (Italy), University Medical Center Groningen (The Netherlands), Cancer Registry of Norway (Norway), Princess Margaret Cancer Center (Canada), Harvard School of Public Health (USA), MD Anderson Cancer Center (USA), Moffitt Cancer Center (USA), Radboud University Medical Center, Nijmegen (The Netherlands), University Medical Center, Hamburg-Eppendorf (Germany), deCODE Genetics (Iceland), University of Turin (Italy), Fred Hutchinson Cancer Research Center (USA), Institute for Cancer Research (Norway), Institute of Cancer Research (UK), Karolinska Institute (Sweden),and Brown University (USA).

**3.3 CONSORTIUM-SPECIFIC COLLABORATORS**

**BCAC**

Thomas Ahearn^1^, Irene L. Andrulis^2,3^, Hoda Anton-Culver^4^, Natalia N. Antonenkova^5^, Volker Arndt^6^, Kristan J. Aronson^7^, Paul L. Auer^8,9^, Annelie Augustinsson^10^, Heiko Becher^11^, Matthias W. Beckmann^12^, Marina Bermisheva^13^, Carl Blomqvist^14,15^, Natalia V. Bogdanova^16,17,5^, Stig E. Bojesen^18,19,20^, Manjeet K. Bolla^21^, Bernardo Bonanni^22^, Hiltrud Brauch^23,24,25^, Hermann Brenner^26,27,25^, Annegien Broeks^28^, Sara Y. Brucker^29^, Thomas Brüning^30^, Barbara Burwinkel^31,32^, Federico Canzian^33^, Jose E. Castelao^34^, Christine L. Clarke^35^, Fergus J. Couch^36^, Kamila Czene^37^, Mary B. Daly^38^, Peter Devilee^39,40^, Thilo Dörk^17^, Isabel dos-Santos-Silva^41^, Alison M. Dunning^42^, Miriam Dwek^43^, Diana M. Eccles^44^, A. Heather Eliassen^45,46^, Peter A. Fasching^47,12^, Henrik Flyger^48^, Lin Fritschi^49^, Manuela Gago-Dominguez^50,51^, Susan M. Gapstur^52^, José A. García-Sáenz^53^, Mia M. Gaudet^52^, Graham G. Giles^54,55,56^, Mark S. Goldberg^57,58^, David E. Goldgar^59^, Pascal Guénel^60^, Eric Hahnen^61,62^, Niclas Håkansson^63^, Ute Hamann^64^, Steven N. Hart^65^, Bernadette A.M. Heemskerk-Gerritsen^66^, Peter Hillemanns^67^, Antoinette Hollestelle^66^, Maartje J. Hooning^66^, John L. Hopper^55^, David J. Hunter^68,46,69^, ABCTB Investigators^70^, Anna Jakubowska^71,72^, Wolfgang Janni^73^, Esther M. John^74^, Audrey Jung^75^, Rudolf Kaaks^75^, Pooja M. Kapoor^75,76^, Elza Khusnutdinova^77,13^, Veli-Matti Kosma^78,79,80^, Vessela N. Kristensen^81,82^, Katerina Kubelka-Sabit^83^, Allison W. Kurian^74,84^, Diether Lambrechts^85,86^, Loic Le Marchand^87^, Annika Lindblom^88,89^, Sibylle Loibl^90^, Jan Lubiński^71^, Michael P. Lux^91^, Arto Mannermaa^78,79,80^, Mehdi Manoochehri^64^, Sara Margolin^92,93^, Dimitrios Mavroudis^94^, Usha Menon^95^, Anna Marie. Mulligan^96,97^, NBCS Collaborators^98,99,100,101,102,103,104,105,106,107,108,109^, Susan L. Neuhausen^110^, Heli Nevanlinna^111^, Katie M. O'Brien^112^, Håkan Olsson^10^, Nick Orr^113^, Julian Peto^41^, Dijana Plaseska-Karanfilska^114^, Ross Prentice^8^, Nadege Presneau^43^, Brigitte Rack^73^, Paolo Radice^115^, Gad Rennert^116^, Hedy S. Rennert^116^, Atocha Romero^117^, Matthias Ruebner^91^, Emmanouil Saloustros^118^, Dale P. Sandler^112^, Rita K. Schmutzler^61,62^, Lukas Schwentner^73^, Christopher Scott^65^, Priyanka Sharma^119^, Xiao-Ou Shu^120^, Christof Sohn^121^, Melissa C. Southey^56,122,123^, John J. Spinelli^124,125^, Jennifer Stone^126,55^, Anthony J. Swerdlow^127,128^, Rulla M. Tamimi^45,46,68^, William J. Tapper^44^, Jack A. Taylor^112,129^, Mary Beth Terry^130^, Amanda E. Toland^131^, Thérèse Truong^60^, Michael Untch^132^, Celine M. Vachon^133^, Qin Wang^21^, Clarice R. Weinberg^134^, Hans Wildiers^135^, Alicja Wolk^63,136^, Xiaohong R. Yang^1^, Wei Zheng^120^, Argyrios Ziogas^4^

^1^Division of Cancer Epidemiology and Genetics, National Cancer Institute, National Institutes of Health, Department of Health and Human Services, Bethesda, MD, USA, ^2^Fred A. Litwin Center for Cancer Genetics, Lunenfeld-Tanenbaum Research Institute of Mount Sinai Hospital, Toronto, ON, Canada, ^3^Department of Molecular Genetics, University of Toronto, Toronto, ON, Canada, ^4^Department of Epidemiology, Genetic Epidemiology Research Institute, University of California Irvine, Irvine, CA, USA, ^5^N.N. Alexandrov Research Institute of Oncology and Medical Radiology, Minsk, Belarus, ^6^Division of Clinical Epidemiology and Aging Research, C070, German Cancer Research Center (DKFZ), Heidelberg, Germany, ^7^Department of Public Health Sciences, and Cancer Research Institute, Queen’s University, Kingston, ON, Canada, ^8^Cancer Prevention Program, Fred Hutchinson Cancer Research Center, Seattle, WA, USA, ^9^Zilber School of Public Health, University of Wisconsin-Milwaukee, Milwaukee, WI, USA, ^10^Department of Cancer Epidemiology, Clinical Sciences, Lund University, Lund, Sweden, ^11^Institute for Medical Biometrics and Epidemiology, University Medical Center Hamburg-Eppendorf, Hamburg, Germany, ^12^Department of Gynecology and Obstetrics, Comprehensive Cancer Center ER-EMN, University Hospital Erlangen, Friedrich-Alexander-University Erlangen-Nuremberg, Erlangen, Germany, ^13^Institute of Biochemistry and Genetics, Ufa Federal Research Centre of the Russian Academy of Sciences, Ufa, Russia, ^14^Department of Oncology, Helsinki University Hospital, University of Helsinki, Helsinki, Finland, ^15^Department of Oncology, Örebro University Hospital, Örebro, Sweden, ^16^Department of Radiation Oncology, Hannover Medical School, Hannover, Germany, ^17^Gynaecology Research Unit, Hannover Medical School, Hannover, Germany, ^18^Copenhagen General Population Study, Herlev and Gentofte Hospital, Copenhagen University Hospital, Herlev, Denmark, ^19^Department of Clinical Biochemistry, Herlev and Gentofte Hospital, Copenhagen University Hospital, Herlev, Denmark, ^20^Faculty of Health and Medical Sciences, University of Copenhagen, Copenhagen, Denmark, ^21^Centre for Cancer Genetic Epidemiology, Department of Public Health and Primary Care, University of Cambridge, Cambridge, UK, ^22^Division of Cancer Prevention and Genetics, IEO, European Institute of Oncology IRCCS, Milan, Italy, ^23^Dr. Margarete Fischer-Bosch-Institute of Clinical Pharmacology, Stuttgart, Germany, ^24^iFIT-Cluster of Excellence, Germany, University of Tuebingen, Tuebingen, Germany, ^25^German Cancer Consortium (DKTK), German Cancer Research Center (DKFZ), Heidelberg, Germany, ^26^Division of Clinical Epidemiology and Aging Research, German Cancer Research Center (DKFZ), Heidelberg, Germany, ^27^Division of Preventive Oncology, German Cancer Research Center (DKFZ) and National Center for Tumor Diseases (NCT), Heidelberg, Germany, ^28^Division of Molecular Pathology, The Netherlands Cancer Institute - Antoni van Leeuwenhoek Hospital, Amsterdam, The Netherlands, ^29^Department of Gynecology and Obstetrics, University of Tübingen, Tübingen, Germany, ^30^Institute for Prevention and Occupational Medicine of the German Social Accident Insurance, Institute of the Ruhr University Bochum (IPA), Bochum, Germany, ^31^Molecular Epidemiology Group, C080, German Cancer Research Center (DKFZ), Heidelberg, Germany, ^32^Molecular Biology of Breast Cancer, University Womens Clinic Heidelberg, University of Heidelberg, Heidelberg, Germany, ^33^Genomic Epidemiology Group, German Cancer Research Center (DKFZ), Heidelberg, Germany, ^34^Oncology and Genetics Unit, Instituto de Investigacion Sanitaria Galicia Sur (IISGS), Xerencia de Xestion Integrada de Vigo-SERGAS, Vigo, Spain, ^35^Westmead Institute for Medical Research, University of Sydney, Sydney, New South Wales, Australia, ^36^Department of Laboratory Medicine and Pathology, Mayo Clinic, Rochester, MN, USA, ^37^Department of Medical Epidemiology and Biostatistics, Karolinska Institutet, Stockholm, Sweden, ^38^Department of Clinical Genetics, Fox Chase Cancer Center, Philadelphia, PA, USA, ^39^Department of Pathology, Leiden University Medical Center, Leiden, The Netherlands, ^40^Department of Human Genetics, Leiden University Medical Center, Leiden, The Netherlands, ^41^Department of Non-Communicable Disease Epidemiology, London School of Hygiene and Tropical Medicine, London, UK, ^42^Department of Oncology, Centre for Cancer Genetic Epidemiology, University of Cambridge, Cambridge, UK, ^43^School of Life Sciences, University of Westminster, London, UK, ^44^Faculty of Medicine, University of Southampton, Southampton, UK, ^45^Channing Division of Network Medicine, Department of Medicine, Brigham and Women's Hospital and Harvard Medical School, Boston, MA, USA, ^46^Department of Epidemiology, Harvard T.H. Chan School of Public Health, Boston, MA, USA, ^47^David Geffen School of Medicine, Department of Medicine Division of Hematology and Oncology, University of California at Los Angeles, Los Angeles, CA, USA, ^48^Department of Breast Surgery, Herlev and Gentofte Hospital, Copenhagen University Hospital, Herlev, Denmark, ^49^School of Public Health, Curtin University, Perth, Western Australia, Australia, ^50^Genomic Medicine Group, Galician Foundation of Genomic Medicine, Instituto de Investigación Sanitaria de Santiago de Compostela (IDIS), Complejo Hospitalario Universitario de Santiago, SERGAS, Santiago de Compostela, Spain, ^51^Moores Cancer Center, University of California San Diego, La Jolla, CA, USA, ^52^Behavioral and Epidemiology Research Group, American Cancer Society, Atlanta, GA, USA, ^53^Medical Oncology Department, Hospital Clínico San Carlos, Instituto de Investigación Sanitaria San Carlos (IdISSC), Centro Investigación Biomédica en Red de Cáncer (CIBERONC), Madrid, Spain, ^54^Cancer Epidemiology Division, Cancer Council Victoria, Melbourne, Victoria, Australia, ^55^Centre for Epidemiology and Biostatistics, Melbourne School of Population and Global Health, The University of Melbourne, Melbourne, Victoria, Australia, ^56^Precision Medicine, School of Clinical Sciences at Monash Health, Monash University, Clayton, Victoria, Australia, ^57^Department of Medicine, McGill University, Montréal, QC, Canada, ^58^Division of Clinical Epidemiology, Royal Victoria Hospital, McGill University, Montréal, QC, Canada, ^59^Department of Dermatology, Huntsman Cancer Institute, University of Utah School of Medicine, Salt Lake City, UT, USA, ^60^Cancer & Environment Group, Center for Research in Epidemiology and Population Health (CESP), INSERM, University Paris-Sud, University Paris-Saclay, Villejuif, France, ^61^Center for Hereditary Breast and Ovarian Cancer, Faculty of Medicine and University Hospital Cologne, University of Cologne, Cologne, Germany, ^62^Center for Integrated Oncology (CIO), Faculty of Medicine and University Hospital Cologne, University of Cologne, Cologne, Germany, ^63^Institute of Environmental Medicine, Karolinska Institutet, Stockholm, Sweden, ^64^Molecular Genetics of Breast Cancer, German Cancer Research Center (DKFZ), Heidelberg, Germany, ^65^Department of Health Sciences Research, Mayo Clinic, Rochester, MN, USA, ^66^Department of Medical Oncology, Family Cancer Clinic, Erasmus MC Cancer Institute, Rotterdam, The Netherlands, ^67^Gynaecology Research Unit, Hannover Medical School, Hannover, Germany, ^68^Program in Genetic Epidemiology and Statistical Genetics, Harvard T.H. Chan School of Public Health, Boston, MA, USA, ^69^Nuffield Department of Population Health, University of Oxford, Oxford, UK, ^70^Australian Breast Cancer Tissue Bank, Westmead Institute for Medical Research, University of Sydney, Sydney, New South Wales, Australia, ^71^Department of Genetics and Pathology, Pomeranian Medical University, Szczecin, Poland, ^72^Independent Laboratory of Molecular Biology and Genetic Diagnostics, Pomeranian Medical University, Szczecin, Poland, ^73^Department of Gynaecology and Obstetrics, University Hospital Ulm, Ulm, Germany, ^74^Department of Medicine, Division of Oncology, Stanford Cancer Institute, Stanford University School of Medicine, Standford, CA, USA, ^75^Division of Cancer Epidemiology, German Cancer Research Center (DKFZ), Heidelberg, Germany, ^76^Faculty of Medicine, University of Heidelberg, Heidelberg, Germany, ^77^Department of Genetics and Fundamental Medicine, Bashkir State Medical University, Ufa, Russia, ^78^Translational Cancer Research Area, University of Eastern Finland, Kuopio, Finland, ^79^Institute of Clinical Medicine, Pathology and Forensic Medicine, University of Eastern Finland, Kuopio, Finland, ^80^Imaging Center, Department of Clinical Pathology, Kuopio University Hospital, Kuopio, Finland, ^81^Department of Cancer Genetics, Institute for Cancer Research, Oslo University Hospital-Radiumhospitalet, Oslo, Norway, ^82^Institute of Clinical Medicine, Faculty of Medicine, University of Oslo, Oslo, Norway, ^83^Department of Histopathology and Cytology, Clinical Hospital Acibadem Sistina, Skopje, Republic of North Macedonia, ^84^Department of Health Research and Policy - Epidemiology, Stanford University School of Medicine, Standford, CA, USA, ^85^VIB Center for Cancer Biology, VIB, Leuven, Belgium, ^86^Laboratory for Translational Genetics, Department of Human Genetics, University of Leuven, Leuven, Belgium, ^87^Epidemiology Program, University of Hawaii Cancer Center, Honolulu, HI, USA, ^88^Department of Molecular Medicine and Surgery, Karolinska Institutet, Stockholm, Sweden, ^89^Department of Clinical Genetics, Karolinska University Hospital, Stockholm, Sweden, ^90^German Breast Group, GmbH, Neu Isenburg, Germany, ^91^Department of Gynaecology and Obstetrics, University Hospital Erlangen, Friedrich-Alexander University Erlangen-Nuremberg, Comprehensive Cancer Center Erlangen-EMN, Erlangen, ^92^Department of Oncology, Södersjukhuset, Stockholm, Sweden, ^93^Department of Clinical Science and Education, Södersjukhuset, Karolinska Institutet, Stockholm, Sweden, ^94^Department of Medical Oncology, University Hospital of Heraklion, Heraklion, Greece, ^95^MRC Clinical Trials Unit at UCL, Institute of Clinical Trials & Methodology, University College London, London, UK, ^96^Department of Laboratory Medicine and Pathobiology, University of Toronto, Toronto, ON, Canada, ^97^Laboratory Medicine Program, University Health Network, Toronto, ON, Canada, ^98^Department of Cancer Genetics, Institute for Cancer Research, Oslo University Hospital-Radiumhospitalet, Oslo, ^99^Institute of Clinical Medicine, Faculty of Medicine, University of Oslo, Oslo, ^100^Department of Research, Vestre Viken Hospital, Drammen, Norway, ^101^Department of Cancer Genetics, Vestre Viken Hospital, Drammen, Norway, ^102^Section for Breast- and Endocrine Surgery, Department of Cancer, Division of Surgery, Cancer and Transplantation Medicine, Oslo University Hospital-Ullevål, Oslo, Norway, ^103^Department of Radiology and Nuclear Medicine, Oslo University Hospital, Oslo, Norway, ^104^Department of Pathology, Akershus University Hospital, Lørenskog, Norway, ^105^Department of Tumor Biology, Institute for Cancer Research, Oslo University Hospital, Oslo, Norway, ^106^Department of Oncology, Division of Surgery, Cancer and Transplantation Medicine, Oslo University Hospital-Radiumhospitalet, Oslo, Norway, ^107^National Advisory Unit on Late Effects after Cancer Treatment, Oslo University Hospital-Radiumhospitalet, Oslo, Norway, ^108^Department of Oncology, Akershus University Hospital, Lørenskog, Norway, ^109^Breast Cancer Research Consortium, Oslo University Hospital, Oslo, Norway, ^110^Department of Population Sciences, Beckman Research Institute of City of Hope, Duarte, CA, USA, ^111^Department of Obstetrics and Gynecology, Helsinki University Hospital, University of Helsinki, Helsinki, Finland, ^112^Epidemiology Branch, National Institute of Environmental Health Sciences, NIH, Research Triangle Park, NC, USA, ^113^Centre for Cancer Research and Cell Biology, Queen's University Belfast, Belfast, Ireland, ^114^Research Centre for Genetic Engineering and Biotechnology "Georgi D. Efremov", Macedonian Academy of Sciences and Arts, Skopje, Republic of North Macedonia, ^115^Unit of Molecular Bases of Genetic Risk and Genetic Testing, Department of Research, Fondazione IRCCS Istituto Nazionale dei Tumori (INT), Milan, Italy, ^116^Clalit National Cancer Control Center, Carmel Medical Center and Technion Faculty of Medicine, Haifa, Israel, ^117^Medical Oncology Department, Hospital Universitario Puerta de Hierro, Madrid, Spain, ^118^Department of Oncology, University Hospital of Larissa, Larissa, Greece, ^119^Department of Internal Medicine, Division of Medical Oncology, University of Kansas Medical Center, Westwood, KS, USA, ^120^Division of Epidemiology, Department of Medicine, Vanderbilt Epidemiology Center, Vanderbilt-Ingram Cancer Center, Vanderbilt University School of Medicine, Nashville, TN, USA, ^121^National Center for Tumor Diseases, University Hospital and German Cancer Research Center, Heidelberg, Germany, ^122^Department of Clinical Pathology, The University of Melbourne, Melbourne, Victoria, Australia, ^123^Cancer Epidemiology Centre, Cancer Council Victoria, Melbourne, Victoria, Australia, ^124^Population Oncology, BC Cancer, Vancouver, BC, Canada, ^125^School of Population and Public Health, University of British Columbia, Vancouver, BC, Canada, ^126^The Curtin UWA Centre for Genetic Origins of Health and Disease, Curtin University and University of Western Australia, Perth, Western Australia, Australia, ^127^Division of Genetics and Epidemiology, The Institute of Cancer Research, London, UK, ^128^Division of Breast Cancer Research, The Institute of Cancer Research, London, UK, ^129^Epigenetic and Stem Cell Biology Laboratory, National Institute of Environmental Health Sciences, NIH, Research Triangle Park, NC, USA, ^130^Department of Epidemiology, Mailman School of Public Health, Columbia University, New York, NY, USA, ^131^Department of Cancer Biology and Genetics, The Ohio State University, Columbus, OH, USA, ^132^Department of Gynecology and Obstetrics, Helios Clinics Berlin-Buch, Berlin, Germay, ^133^Department of Health Science Research, Division of Epidemiology, Mayo Clinic, Rochester, MN, USA, ^134^Biostatistics and Computational Biology Branch, National Institute of Environmental Health Sciences, NIH, Research Triangle Park, NC, USA, ^135^Leuven Multidisciplinary Breast Center, Department of Oncology, Leuven Cancer Institute, University Hospitals Leuven, Leuven, Belgium, ^136^Department of Surgical Sciences, Uppsala University, Uppsala, Sweden

**BEACON**

Lesley A. Anderson^1^, Leslie Bernstein^2^, Nigel C. Bird^3^, Wong-Ho Chow^4^, Doug A. Corley^5^, Rebecca C. Fitzgerald^6^, Marilie D. Gammon^7^, Laura J. Hardie^8^, Prasad G. Iyer^9^, Jesper Lagergren^10,11^, Geoffrey Liu^12^, Brian J. Reid^13^, Harvey A. Risch^14^, Nick J. Shaheen^15^, Tom L. Vaughan^16^, Anna H. Wu^17,18^, Weimin Ye^19^

^1^Centre for Public Health, School of Medicine, Dentistry and Biomedical Science, Queen's University Belfast, Belfast, UK, ^2^Division of Biomarkers of Early Detection and Prevention, Beckman Research Institute, City of Hope, Duarte, CA, USA, ^3^Department of Oncology, Medical School, University of Sheffield, Sheffield, UK, ^4^Department of Epidemiology, The University of Texas MD Anderson Cancer Center, Houston, TX, USA, ^5^Division of Research, Kaiser Permanente Northern California, Oakland, CA, USA, ^6^Medical Research Council Cancer Unit, Hutchison-MRC Research Centre, University of Cambridge, Cambridge, UK, ^7^Department of Epidemiology, University of North Carolina at Chapel Hill, Chapel Hill, NC, USA, ^8^Department of Clinical and Population Science, Leeds Institute of Cardiovascular and Metabolic Medicine, School of Medicine, University of Leeds, Leeds, UK, ^9^Division of Gastroenterology and Hepatology, Mayo Clinic, Rochester, MN, USA, ^10^Department of Molecular Medicine and Surgery, Upper Gastrointestinal Surgery, Karolinska Institutet, Karolinska University Hospital, Stockholm, Sweden, ^11^School of Cancer and Pharmaceutical Sciences, King's College London, London, UK, ^12^Princess Margaret Cancer Center, Toronto, ON, Canada, ^13^Divisions of Human Biology and Public Health Sciences, Fred Hutchinson Cancer Research Center, Seattle, WA, USA, ^14^Chronic Disease Epidemiology, Yale School of Public Health, New Haven, CT, USA, ^15^Division of Gastroenterology and Hepatology, University of North Carolina at Chapel Hill, Chapel Hill, NC, USA, ^16^Program in Epidemiology, Division of Public Health Sciences, Fred Hutchinson Cancer Research Center, Seattle, WA, USA, ^17^USC Norris Comprehensive Cancer Center,, University of Southern California, Los Angeles, CA, USA, ^18^Department of Preventive Medicine, Keck School of Medicine, University of Southern California, Los Angeles, CA, USA, ^19^Department of Medical Epidemiology and Biostatistics, Karolinska Institutet, Stockholm, Sweden

**ECAC**

Frederic Amant^1^, Daniela Annibali^1^, Katie Ashton^2,3,4^, John Attia^2,5^, Paul L. Auer^6,7^, Matthias W. Beckmann^8^, Amanda Black^9^, Louise Brinton^9^, Daniel D. Buchanan^10,11,12,13^, Stephen J. Chanock^9^, Chu Chen^14^, Maxine M. Chen^15^, Timothy H.T. Cheng^16^, Linda S. Cook^17,18^, Marta Crous-Bous^19,15^, Kamila Czene^20^, Jeroen Depreeuw^1,21,22^, Jennifer Anne Doherty^23^, Thilo Dörk^24^, Sean C. Dowdy^25^, Alison M. Dunning^26^, Matthias Dürst^27^, Douglas F. Easton^26,28^, Arif B. Ekici^29^, Peter A. Fasching^30,8^, Brooke L. Fridley^31^, Christine M. Friedenreich^18^, Montserrat García-Closas^9,32^, Mia M. Gaudet^33^, Graham G. Giles^34,11,35^, Dylan M. Glubb^36^, Ellen L. Goode^37^, Maggie Gorman^16^, Christopher A. Haiman^38^, Per Hall^20,39^, Susan E. Hankinson^19,40^, Catherine S. Healey^26^, Alexander Hein^8^, Peter Hillemanns^24^, Shirley Hodgson^41^, Erling Hoivik^42,43^, Elizabeth G. Holliday^2,5^, David J. Hunter^44,15,45^, Angela Jones^16^, Peter Kraft^44,15^, Camilla Krakstad^43,42^, Diether Lambrechts^46,22^, Loic Le Marchand^47^, Xiaolin Liang^48^, Annika Lindblom^49,50^, Jolanta Lissowska^51^, Jirong Long^52^, Lingeng Lu^53^, Anthony M. Magliocco^54^, Lynn Martin^55^, Mark McEvoy^5^, Roger L. Milne^34,11,35^, Miriam Mints^56^, Rami Nassir^57^, Irene Orlow^48^, Geoffrey Otton^58^, Claire Palles^16^, Paul DP. Pharoah^26,28^, Loreall Pooler^38^, Tony Proietto^58^, Timothy R. Rebbeck^59,60^, Stefan P. Renner^8^, Harvey A. Risch^53^, Matthias Rübner^8^, Ingo Runnebaum^27^, Carlotta Sacerdote^61,62^, Gloria E. Sarto^63^, Fredrick Schumacher^64^, Rodney J. Scott^65,4,2^, V. Wendy Setiawan^38^, Mitul Shah^26^, Xin Sheng^38^, Xiao-Ou Shu^52^, Melissa C. Southey^35,10^, Emma Tham^49,66^, Jone Trovik^42,43^, Constance Turman^15^, David Van Den Berg^38^, Adriaan Vanderstichele^67^, Zhaoming Wang^9^, Penelope M. Webb^68^, Nicolas Wentzensen^9^, Henrica MJ. Werner^42,43^, Stacey J. Winham^69^, Lucy Xia^38^, Yong-Bing Xiang^70^, Hannah P. Yang^9^, Herbert Yu^47^, Wei Zheng^52^

^1^Department of Obstetrics and Gynecology, Division of Gynecologic Oncology, University Hospitals KU Leuven, University of Leuven, Leuven, Belgium, ^2^Hunter Medical Research Institute, John Hunter Hospital, Newcastle, New South Wales, Australia, ^3^Centre for Information Based Medicine, University of Newcastle, Callaghan, New South Wales, Australia, ^4^Discipline of Medical Genetics, School of Biomedical Sciences and Pharmacy, Faculty of Health, University of Newcastle, Callaghan, New South Wales, Australia, ^5^Centre for Clinical Epidemiology and Biostatistics, School of Medicine and Public Health, University of Newcastle, Callaghan, New South Wales, Australia, ^6^Cancer Prevention Program, Fred Hutchinson Cancer Research Center, Seattle, WA, USA, ^7^Zilber School of Public Health, University of Wisconsin-Milwaukee, Milwaukee, WI, USA, ^8^Department of Gynecology and Obstetrics, Comprehensive Cancer Center ER-EMN, University Hospital Erlangen, Friedrich-Alexander-University Erlangen-Nuremberg, Erlangen, Germany, ^9^Division of Cancer Epidemiology and Genetics, National Cancer Institute, National Institutes of Health, Rockville, MD, USA, ^10^Department of Clinical Pathology, The University of Melbourne, Melbourne, Victoria, Australia, ^11^Centre for Epidemiology and Biostatistics, Melbourne School of Population and Global Health, The University of Melbourne, Melbourne, Victoria, Australia, ^12^Genetic Medicine and Family Cancer Clinic, Royal Melbourne Hospital, Parkville, Victoria, Australia, ^13^University of Melbourne Centre for Cancer Research, Victorian Comprehensive Cancer Centre, Parkville, Victoria, Australia, ^14^Epidemiology Program, Fred Hutchinson Cancer Research Center, Seattle, WA, USA, ^15^Department of Epidemiology, Harvard T.H. Chan School of Public Health, Boston, MA, USA, ^16^Wellcome Trust Centre for Human Genetics and Oxford NIHR Biomedical Research Centre, University of Oxford, Oxford, UK, ^17^University of New Mexico Health Sciences Center, University of New Mexico, Albuquerque, NM, USA, ^18^Department of Cancer Epidemiology and Prevention Research, Alberta Health Services, Calgary, AB, Canada, ^19^Department of Medicine, Channing Division of Network Medicine, Brigham and Women's Hospital and Harvard Medical School, Boston, MA, USA, ^20^Department of Medical Epidemiology and Biostatistics, Karolinska Institutet, Stockholm, Sweden, ^21^Vesalius Research Center, VIB, Leuven, Belgium, ^22^Department of Human Genetics, Laboratory for Translational Genetics, University of Leuven, Leuven, Belgium, ^23^Department of Population Health Sciences, Huntsman Cancer Institute, University of Utah, Salt Lake City, UT, USA, ^24^Gynaecology Research Unit, Hannover Medical School, Hannover, Germany, ^25^Department of Obstetrics and Gynecology, Division of Gynecologic Oncology, Mayo Clinic, Rochester, MN, USA, ^26^Department of Oncology, Centre for Cancer Genetic Epidemiology, University of Cambridge, Cambridge, UK, ^27^Department of Gynaecology, Jena University Hospital - Friedrich Schiller University, Jena, Germany, ^28^Department of Public Health and Primary Care, Centre for Cancer Genetic Epidemiology, University of Cambridge, Cambridge, UK, ^29^Institute of Human Genetics, University Hospital Erlangen, Friedrich-Alexander University Erlangen-Nuremberg, Comprehensive Cancer Center Erlangen-EMN, Erlangen, Germany, ^30^Department of Medicine Division of Hematology and Oncology, David Geffen School of Medicine, University of California at Los Angeles, Los Angeles, CA, USA, ^31^Department of Biostatistics, Kansas University Medical Center, Kansas City, KS, USA, ^32^Division of Genetics and Epidemiology, Institute of Cancer Research, London, UK, ^33^Behavioral and Epidemiology Research Group, American Cancer Society, Atlanta, GA, USA, ^34^Cancer Epidemiology Division, Cancer Council Victoria, Melbourne, Victoria, Australia, ^35^Precision Medicine, School of Clinical Sciences at Monash Health, Monash University, Clayton, Victoria, Australia, ^36^Department of Genetics and Computational Biology, QIMR Berghofer Medical Research Institute, Brisbane, Queensland, Australia, ^37^Department of Health Science Research, Division of Epidemiology, Mayo Clinic, Rochester, MN, USA, ^38^Department of Preventive Medicine, Keck School of Medicine, University of Southern California, Los Angeles, CA, USA, ^39^Department of Oncology, Södersjukhuset, Stockholm, Sweden, ^40^Department of Biostatistics & Epidemiology, University of Massachusetts, Amherst, Amherst, MA, USA, ^41^Department of Clinical Genetics, St George's, University of London, London, UK, ^42^Department of Clinical Science, Centre for Cancerbiomarkers, University of Bergen, Bergen, Norway, ^43^Department of Gynecology and Obstetrics, Haukeland University Hospital, Bergen, Norway, ^44^Program in Genetic Epidemiology and Statistical Genetics, Harvard T.H. Chan School of Public Health, Boston, MA, USA, ^45^Nuffield Department of Population Health, University of Oxford, Oxford, UK, ^46^VIB Center for Cancer Biology, VIB, Leuven, Belgium, ^47^Epidemiology Program, University of Hawaii Cancer Center, Honolulu, HI, USA, ^48^Department of Epidemiology and Biostatistics, Memorial Sloan-Kettering Cancer Center, New York, NY, USA, ^49^Department of Molecular Medicine and Surgery, Karolinska Institutet, Stockholm, Sweden, ^50^Department of Clinical Genetics, Karolinska University Hospital, Stockholm, Sweden, ^51^Department of Cancer Epidemiology and Prevention, M. Sklodowska-Curie Cancer Center, Oncology Institute, Warsaw, Poland, ^52^Department of Medicine, Vanderbilt Epidemiology Center, Vanderbilt-Ingram Cancer Center, Division of Epidemiology, Vanderbilt University School of Medicine, Nashville, TN, USA, ^53^Chronic Disease Epidemiology, Yale School of Public Health, New Haven, CT, USA, ^54^Department of Anatomic Pathology, Moffitt Cancer Center & Research Institute, Tampa, FL, USA, ^55^Institute of Cancer and Genomic Sciences, University of Birmingham, Birmingham, UK, ^56^Department of Women's and Children's Health, Karolinska Institutet, Stockholm, Sweden, ^57^Department of Biochemistry and Molecular Medicine, University of California Davis, Davis, CA, USA, ^58^School of Medicine and Public Health, University of Newcastle, Callaghan, New South Wales, Australia, ^59^Harvard T.H. Chan School of Public Health, Boston, MA, USA, ^60^Dana-Farber Cancer Institute, Boston, MA, USA, ^61^Center for Cancer Prevention (CPO-Peimonte), Turin, Italy, ^62^Human Genetics Foundation (HuGeF), Turino, Italy, ^63^Department of Obstetrics and Gynecology, School of Medicine and Public Health, University of Wisconsin, Madison, WI, USA, ^64^Department of Population and Quantitative Health Sciences, Case Western Reserve University, Cleveland, OH, USA, ^65^Pathology North, Division of Molecular Medicine, John Hunter Hospital, Newcastle, New South Wales, Australia, ^66^Clinical Genetics, Karolinska Institutet, Stockholm, Sweden, ^67^Department of Obstetrics and Gynaecology and Leuven Cancer Institute, Division of Gynecologic Oncology, University Hospitals Leuven, Leuven, Belgium, ^68^Population Health Department, QIMR Berghofer Medical Research Institute, Brisbane, Queensland, Australia, ^69^Department of Health Science Research, Division of Biomedical Statistics and Informatics, Mayo Clinic, Rochester, MN, USA, ^70^State Key Laboratory of Oncogene and Related Genes & Department of Epidemiology, Shanghai Cancer Institute, Renji Hospital, Shanghai Jiaotong University School of Medicine, Shanghai, China

**GECCO, CORECT, CCFR**

Goncalo R. Abecasis^1^, Yoon-Ok Ahn^2^, Barbara Banbury^3^, John A. Baron^4^, Sonja I. Berndt^5^, Stéphane Bézieau^6^, Stephanie A. Bien^3^, Hermann Brenner^7,8,9^, Daniel D. Buchanan^10,11,12^, Qiuyin Cai^13^, Andrew T. Chan^14,15,16^, Jenny Chang-Claude^17,18^, David V. Conti^19^, Keith R. Curtis^3^, Christopher K. Edlund^19^, Dallas R. English^20,21^, Jane Figueiredo^22^, Steven J. Gallinger^23^, Graham G. Giles^21,20^, Robert W. Haile^24^, Tabitha A. Harrison^3^, John L. Hopper^20,25^, Thomas J. Hudson^26^, David J. Hunter^27,28^, Jeroen R. Huyghe^3^, Jae Hwan Oh^2^, Sun Ha Jee^29^, Wei-Hua Jia^30^, Amit D. Joshi^27,16^, Keum Ji Jung^31^, Yoichiro Kamatani^32^, Dong-Hyun Kim^33^, Jeongseon Kim^34^, Charles Kooperberg^3^, Sébastien Küry^6^, Sun-Seog Kweon^2,35^, Loic Le Marchand^36^, Mathieu Lemire^37^, Li Li^38^, Yi Lin^3^, Noralane M. Lindor^39^, Jirong Long^13^, Yingchang Lu^13^, Koichi Matsuda^40^, Keitaro Matsuo^41,42^, Roger L. Milne^21,20^, Polly A. Newcomb^3,43^, Deborah A. Nickerson^44^, Shuji Ogino^45,46,27^, Isao Oze^41^, John D. Potter^3^, Conghui Qu^3^, Gad Rennert^47,48,49^, Hedy S. Rennert^47,48,49^, Lori C. Sakoda^50,3^, Robert E. Schoen^51^, Fredrick R. Schumacher^52^, Min-Ho Shin^2^, Aesun Shin^53^, Xiao-Ou Shu^54^, Martha L. Slattery^55^, Melissa C. Southey^56^, Stephen N. Thibodeau^57^, Emily White^3,58^, Michael O. Woods^59^, Yong-Bing Xiang^60^, Brent W. Zanke^61^, Yi-Xin Zeng^30^

^1^Department of Biostatistics and Center for Statistical Genetics, University of Michigan, Ann Arbor, MI, USA, ^2^Department of Preventive Medicine, Seoul National University College of Medicine, Seoul, South Korea, ^3^Public Health Sciences Division, Fred Hutchinson Cancer Research Center, Seattle, WA, USA, ^4^Department of Medicine, University of North Carolina School of Medicine, Chapel Hill, NC, USA, ^5^Division of Cancer Epidemiology and Genetics, National Cancer Institute, National Institutes of Health, Rockville, MD, USA, ^6^Service de Génétique Médicale, Centre Hospitalier Universitaire (CHU) Nantes France, Nantes, France, ^7^Division of Clinical Epidemiology and Aging Research, German Cancer Research Center (DKFZ), Heidelberg, Germany, ^8^Division of Preventive Oncology, German Cancer Research Center (DKFZ) and National Center for Tumor Diseases (NCT), Heidelberg, Germany, ^9^German Cancer Consortium (DKTK), German Cancer Research Center (DKFZ), Heidelberg, Germany, ^10^Colorectal Oncogenomics Group, Department of Clinical Pathology, The University of Melbourne, Parkville, Victoria, Australia, ^11^University of Melbourne Centre for Cancer Research, Victorian Comprehensive Cancer Centre, Parkville, Victoria, Australia, ^12^Genetic Medicine and Family Cancer Clinic, The Royal Melbourne Hospital, Parkville, Victoria, Australia, ^13^Division of Epidemiology, Department of Medicine, Vanderbilt-Ingram Cancer Center, Vanderbilt Epidemiology Center, Vanderbilt University School of Medicine, Nashville, TN, USA, ^14^Division of Gastroenterology, Massachusetts General Hospital and Harvard Medical School, Boston, MA, USA, ^15^Channing Division of Network Medicine, Brigham and Women's Hospital and Harvard Medical School, Boston, MA, USA, ^16^Clinical and Translational Epidemiology Unit, Massachusetts General Hospital and Harvard Medical School, Boston, MA, USA, ^17^Division of Cancer Epidemiology, German Cancer Research Center (DKFZ), Heidelberg, Germany, ^18^University Medical Centre Hamburg-Eppendorf, University Cancer Centre Hamburg (UCCH), Hamburg, Germany, ^19^Department of Preventive Medicine, USC Norris Comprehensive Cancer Center, Keck School of Medicine, University of Southern California, Los Angeles, CA, USA, ^20^Centre for Epidemiology and Biostatistics, Melbourne School of Population and Global Health, The University of Melbourne, Melbourne, Victoria, Australia, ^21^Cancer Epidemiology Division, Cancer Council Victoria, Melbourne, Victoria, Australia, ^22^Samuel Oschin Comprehensive Cancer Institute, Cedars-Sinai Medical Center, Los Angeles, CA, USA, ^23^Lunenfeld Tanenbaum Research Institute, Mount Sinai Hospital, University of Toronto, Toronto, Ontario, Canada, ^24^Department of Medicine, Division of Oncology, Stanford University, Stanford, CA, USA, ^25^Department of Epidemiology, School of Public Health and Institute of Health and Environment, Seoul National University, Seoul, South Korea, ^26^Ontario Institute for Cancer Research, Toronto, Ontario, Canada, ^27^Department of Epidemiology, Harvard T.H. Chan School of Public Health, Harvard University, Boston, MA, USA, ^28^Nuffield Department of Population Health, University of Oxford, Oxford, UK, ^29^Department of Epidemiology and Health Promotion, Graduate School of Public Health, Yonsei University, Seoul, South Korea, ^30^State Key Laboratory of Oncology in South China, Cancer Center, Sun Yat-sen University, Guangzhou, China, ^31^Institute for Health Promotion, Graduate School of Public Health, Yonsei University, Seoul, South Korea, ^32^Laboratory for Statistical Analysis, RIKEN Center for Integrative Medical Sciences, Kanagawa, Japan, ^33^Department of Social and Preventive Medicine, Hallym University College of Medicine, Okcheon-dong, South Korea, ^34^Department of Cancer Biomedical Science, Graduate School of Cancer Science and Policy, National Cancer Center, Gyeonggi-do, South Korea, ^35^Jeonnam Regional Cancer Center, Chonnam National University Hwasun Hospital, Hwasun, South Korea, ^36^University of Hawaii Cancer Center, University of Hawaii, Honolulu, HI, USA, ^37^PanCuRx Translational Research Initiative, Ontario Institute for Cancer Research, Toronto, Ontario, Canada, ^38^Department of Family Medicine, University of Virginia, Charlottesville, VA, USA, ^39^Department of Health Science Research, Mayo Clinic Arizona, Scottsdale, AZ, USA, ^40^Laboratory of Clinical Genome Sequencing, Department of Computational Biology and Medical Sciences, Graduate School of Frontier Sciences, University of Tokyo, Tokyp, Japan, ^41^Division of Molecular and Clinical Epidemiology, Aichi Cancer Center Research Institute, Nagoya, Japan, ^42^Department of Epidemiology, Nagoya University Graduate School of Medicine, Nagoya, Japan, ^43^School of Public Health, University of Washington, Seattle, WA, USA, ^44^Department of Genome Sciences, University of Washington, Seattle, WA, USA, ^45^Program in MPE Molecular Pathological Epidemiology, Department of Pathology, Brigham and Women's Hospital, Harvard Medical School, Boston, MA, USA, ^46^Department of Oncologic Pathology, Dana-Farber Cancer Institute, Boston, MA, USA, ^47^Department of Community Medicine and Epidemiology, Lady Davis Carmel Medical Center, Haifa, Israel, ^48^Ruth and Bruce Rappaport Faculty of Medicine, Technion-Israel Institute of Technology, Haifa, Israel, ^49^Clalit National Cancer Control Center, Haifa, Israel, ^50^Division of Research, Kaiser Permanente Northern California, Oakland, CA, USA, ^51^Department of Medicine and Epidemiology, University of Pittsburgh Medical Center, Pittsburgh, PA, USA, ^52^Department of Population and Quantitative Health Sciences, Case Western Reserve University, Cleveland, OH, USA, ^53^Department of Preventive Medicine, Seoul National University College of Medicine, Seoul National University Cancer Research Institute, Seoul, South Korea, ^54^Vanderbilt University Medical Center, Nashville, TN, USA, ^55^Department of Internal Medicine, University of Utah, Salt Lake City, Utah, USA, ^56^Genetic Epidemiology Laboratory, Department of Pathology, The University of Melbourne, Melbourne, Australia, ^57^Division of Laboratory Genetics, Department of Laboratory Medicine and Pathology, Mayo Clinic, Rochester, MN, USA, ^58^Department of Epidemiology, University of Washington School of Public Health, Seattle, WA, USA, ^59^Memorial University of Newfoundland, Discipline of Genetics, St. John's, Canada, ^60^State Key Laboratory of Oncogenes and Related Genes & Department of Epidemiology, Shanghai Cancer Institute, Renji Hospital, Shanghai Jiaotong University School of Medicine, Shanghai, China, ^61^University of Ottawa, Division of Hematology, Ottawa, Canada

**GenoMEL**

Lars A. Akslen^1^, Christopher I. Amos^2^, Per A. Andresen^3^, Marie-Françoise Avril^4^, Esther Azizi^5^, Jennifer H. Barrett^6^, Giovanna Bianchi Scarrà^7^, Myriam Brossard^8,9^, Kevin M. Brown^10^, Kathryn P. Burdon^11^, Wei V. Chen^12^, Jamie E. Craig^13^, Anne E. Cust^14^, Tadeusz Dębniak^15^, David L. Duffy^16^, Alison M. Dunning^17^, Douglas F. Easton^17,18^, David E. Elder^19^, Shenying Fang^20^, Eitan Friedman^21^, Pilar Galan^22^, Paola Ghiorzo^7^, Elizabeth M. Gillanders^23^, Alisa M. Goldstein^10^, Nelleke A. Gruis^24^, Jiali Han^25^, Johan Hansson^26^, Mark Harland^27^, Nicholas K. Hayward^28^, Per Helsing^29^, Marko Hočevar^30^, Veronica Höiom^26^, Christian Ingvar^31^, Peter A. Kanetsky^32^, Rajiv Kumar^33^, Katerina P. Kypreou^34^, Maria Teresa Landi^10^, Julie Lang^35^, G. Mark Lathrop^36^, Jeffrey E. Lee^20^, Jan Lubiński^15^, Rona M. Mackie^37^, Graham J. Mann^38^, Nicholas G. Martin^16^, Anders Molven^39^, Grant W. Montgomery^40^, Eric K. Moses^41^, Julia A. Newton Bishop^27^, Srdjan Novaković^42^, Dale R. Nyholt^43^, Håkan Olsson^44^, Nick Orr^45^, Paul D.P. Pharoah^17^, Karen A. Pooley^18^, Susana Puig^46^, Joan Anton Puig Butille^46^, Abrar A. Qureshi^47^, Graham L. Radford-Smith^48^, Juliette Randerson-Moor^27^, Dirk Schadendorf^49^, Hans-Joachim Schulze^50^, Lisa A. Simms^48^, Fengju Song^51^, Alexander J. Stratigos^34^, Anthony J. Swerdlow^52^, John C. Taylor^6^, Nienke van der Stoep^53^, Remco van Doorn^24^, David C. Whiteman^54^

^1^Centre for Cancer Biomarkers CCBIO, Department of Clinical Medicine, University of Bergen, Bergen, Norway, ^2^Department of Medicine, Baylor College of Medicine, Houston, TX, USA, ^3^Department of Pathology, Molecular Pathology, Oslo University Hospital, Rikshospitalet, Oslo, Norway, ^4^Assistance Publique–Hôpitaux de Paris, Hôpital Cochin, Service de Dermatologie, Université Paris Descartes, Paris, France, ^5^Department of Dermatology, Sheba Medical Center, Tel Hashomer, Sackler Faculty of Medicine, Tel Aviv, Israel, ^6^Division of Pathology and Data Analytics, Leeds Institute of Medical Research, University of Leeds, Leeds, UK, ^7^Department of Internal Medicine and Medical Specialties, University of Genoa, Genoa, Italy, ^8^Université de Paris, INSERM, UMR-1124, Paris, France, ^9^Division of Biotstatistics, Lunenfeld Tanenbaum Research Institute, Toronto, Canada, ^10^Division of Cancer Epidemiology and Genetics, National Cancer Institute, National Institutes of Health, Rockville, MD, USA, ^11^Menzies Institute for Medical Research, University of Tasmania, Hobart, Tasmania, Australia, ^12^Department of Genetics, The University of Texas MD Anderson Cancer Center, Houston, TX, USA, ^13^Department of Ophthalmology, Flinders University, Adelaide, Australia, ^14^Cancer Epidemiology and Services Research, Sydney School of Public Health, University of Sydney, Sydney, New South Wales, Australia, ^15^International Hereditary Cancer Center, Pomeranian Medical University, Szczecin, Poland, ^16^Genetic Epidemiology, QIMR Berghofer Medical Research Institute, Brisbane, Australia, ^17^Centre for Cancer Genetic Epidemiology, Department of Oncology, University of Cambridge, Cambridge, UK, ^18^Centre for Cancer Genetic Epidemiology, Department of Public Health and Primary Care, University of Cambridge, Cambridge, UK, ^19^Department of Pathology and Laboratory Medicine, Perelman School of Medicine at the University of Pennsylvania, Philadelphia, PA, USA, ^20^Department of Surgical Oncology, The University of Texas MD Anderson Cancer Center, Houston, TX, USA, ^21^Oncogenetics Unit, Sheba Medical Center, Tel Hashomer, Sackler Faculty of Medicine, Tel Aviv, Israel, ^22^Université Paris 13, Equipe de Recherche en Epidémiologie Nutritionnelle (EREN), Centre de Recherche en Epidémiologie et Biostatistiques de Sorbonne Paris Cité (CRESS), Institut National de la Santé et de la Recherche Médicale (INSERM U1153), Institut National de la Recherche Agronomique (INRA U1125), Conservatoire National des Arts et Métiers, Bobigny, France, ^23^Inherited Disease Research Branch, National Human Genome Research Institute, National Institutes of Health, Baltimore, MD, USA, ^24^Department of Dermatology, Leiden University Medical Centre, Leiden, The Netherlands, ^25^Department of Epidemiology, Richard M. Fairbanks School of Public Health, Melvin and Bren Simon Cancer Center, Indiana University, Indianapolis, IN, USA, ^26^Department of Oncology-Pathology, Karolinska Institutet, Karolinska University Hospital, Stockholm, Sweden, ^27^Division of Haematology and Immunology, Leeds Institute of Medical Research, University of Leeds, Leeds, UK, ^28^Oncogenomics, QIMR Berghofer Medical Research Institute, Brisbane, Australia, ^29^Department of Dermatology, Oslo University Hospital, Rikshospitalet, Oslo, Norway, ^30^Department of Surgical Oncology, Institute of Oncology Ljubljana, Ljubljana, Slovenia, ^31^Department of Surgery, Clinical Sciences, Lund University, Lund, Sweden, ^32^Department of Cancer Epidemiology, H. Lee Moffitt Cancer Center and Research Institute, Tampa, FL, USA, ^33^Division of Molecular Genetic Epidemiology, German Cancer Research Center, Heidelberg, Germany, ^34^Department of Dermatology, University of Athens School of Medicine, Andreas Sygros Hospital, Athens, Greece, ^35^Department of Medical Genetics, University of Glasgow, Glasgow, UK, ^36^McGill University, Montreal, Canada, ^37^Department of Public Health, University of Glasgow, Glasgow, UK, ^38^Centre for Cancer Research, University of Sydney at Westmead, Millennium Institute for Medical Research and Melanoma Institute Australia, Sydney, Australia, ^39^Department of Pathology, Haukeland University Hospital, Bergen, Norway, ^40^Molecular Biology, The University of Queensland, Brisbane, Australia, ^41^Centre for Genetic Origins of Health and Disease, Faculty of Medicine, Dentistry and Health Sciences, The University of Western Australia, Perth, Western Australia, Australia, ^42^Department of Molecular Diagnostics, Institute of Oncology Ljubljana, Ljubljana, Slovenia, ^43^Institute of Health and Biomedical Innovation, Queensland University of Technology, Brisbane, Queensland, Australia, ^44^Department of Oncology/Pathology, Clinical Sciences, Lund University, Lund, Sweden, ^45^Centre for Cancer Research and Cell Biology, Queen's University Belfast, Belfast, UK, ^46^Melanoma Unit, Dermatology Department & Biochemistry and Molecular Genetics Departments, Hospital Clinic, Institut de Investigacó Biomèdica August Pi Suñe, Barcelona, Spain, ^47^Department of Dermatology, The Warren Alpert Medical School of Brown University, Providence, RI, USA, ^48^Inflammatory Bowel Diseases, QIMR Berghofer Medical Research Institute, Brisbane, Australia, ^49^Department of Dermatology, University Hospital Essen, Essen, Germany, ^50^Department of Dermatology, Fachklinik Hornheide, Institute for Tumors of the Skin at the University of Münster, Münster, Germany, ^51^Departments of Epidemiology and Biostatistics, Key Laboratory of Cancer Prevention and Therapy, National Clinical Research Center of Cancer, Tianjin, P.R. China, ^52^Division of Genetics and Epidemiology, The Institute of Cancer Research, London, UK, ^53^Department of Clinical Genetics, Center of Human and Clinical Genetics, Leiden University Medical Centre, Leiden, The Netherlands, ^54^Cancer Control Group, QIMR Berghofer Medical Research Institute, Brisbane, Australia

**GICC**

Francis Ali-Osman^1^, Christopher I. Amos^2^, Georgina Armstrong^2^, Jonine Bernstein^3^, Elizabeth Claus^4^, Dora Il'yasova^5^, Robert Jenkins^6^, Christoffer Johansen^7^, Daniel Lachance^8^, Rose Lai^9^, Ryan Merrell^10^, Sara Olson^3^, Quinn Ostrom^2^, Siegal Sadetzki^11^, Michael Scheurer^12^, Joellen Schildkraut^13^, Sanjay Shete^14^

^1^Department of Surgery, Duke University Medical Center, Durham, NC, USA, ^2^Department of Medicine, Section of Epidemiology and Population Sciences, Baylor College of Medicine, Houston, TX, USA, ^3^Department of Epidemiology and Biostatistics, Memorial Sloan Kettering Cancer Center, New York, NY, USA, ^4^Department of Epidemiology and Public Health, Yale University School of Medicine, New Haven, CT, USA, ^5^Department of Epidemiology and Biostatistics, Georgia State University School of Public Health, Atlanta, GA, USA, ^6^Department of Laboratory Medicine and Pathology, Mayo Clinic Comprehensive Cancer Center, Rochester, MN, USA, ^7^Rigshospitalet and Survivorship Research Unit, The Danish Cancer Society Research Center, Copenhagen, Denmark, ^8^Department of Neurology, Mayo Clinic Comprehensive Cancer Center, Rochester, MN, USA, ^9^Departments of Neurology and Preventive Medicine, University of Southern California, Keck School of Medicine, Los Angeles, CA, USA, ^10^Department of Neurology, NorthShore University Health System, Evanston, IL, USA, ^11^Cancer and Radiation Epidemiology Unit, Gertner Institute, Chaim Sheba Medical Center, Tel Hashomer, Israel, ^12^Department of Pediatrics, Baylor College of Medicine, Houston, TX, USA, ^13^Department of Public Health Sciences, University of Virginia School of Medicine, Charlottesville, VA, USA, ^14^Department of Biostatistics, University of Texas MD Anderson Cancer Center, Houston, TX, USA

**ILCCO/Integral**

Demetrios Albanes^1^, Melinda C. Aldrich^2^, Christopher I. Amos^3^, Angeline S. Andrew^4^, Susanne M. Arnold^5^, Heike Bickeböller^6^, Stig E. Bojesen^7,8,9^, Paul Brennan^10^, Hans Brunnström^11^, Neil Caporaso^1^, Chu Chen^12^, David C. Christiani^13^, John K. Field^14^, Kjell Grankvist^15^, Rayjean J. Hung^16^, Mattias Johansson^10^, Mikael Johansson^17^, Lambertus A. Kiemeney^18^, Stephen Lam^19^, Maria Teresa Landi^1^, Philip Lazarus^20^, Geoffrey Liu^21^, Loic Le Marchand^22^, Olle Melander^23,24^, Gadi Rennert^25^, Angela Risch^26,27,28^, Matthew B. Schabath^29^, Hongbing Shen^30^, Sanjay S. Shete^31^, Adonina Tardon^32^, M. Dawn Teare^33^, H-Erich Wichmann^34,35,36^, Shan Zienolddiny^37^

^1^Division of Cancer Epidemiology and Genetics, National Cancer Institute, Rockville, MD, USA, ^2^Department of Thoracic Surgery, Division of Epidemiology, Vanderbilt University Medical Center, Nashville, TN, USA, ^3^Institute for Clinical and Translational Research, Dan L. Duncan Comprehensive Cancer Center, Baylor College of Medicine, Houston, TX, USA, ^4^Norris Cotton Cancer Center, Dartmouth Geisel School of Medicine, Lebanon, NH, USA, ^5^Markey Cancer Center, University of Kentucky, Lexington, KY, USA, ^6^University Medical Center Goettingen, Goettingen, Germany, ^7^Copenhagen General Population Study, Herlev and Gentofte Hospital, Herlev, Denmark, ^8^Department of Clinical Biochemistry, Herlev and Gentofte Hospital, Copenhagen University Hospital, Copenhagen, Denmark, ^9^Faculty of Health and Medical Sciences, University of Copenhagen, Copenhagen, Denmark, ^10^International Agency for Research on Cancer, Lyon, France, ^11^Pathology, Department of Clinical Sciences Lund, Laboratory Medicine Region Skåne, Lund University, Lund, Sweden, ^12^Public Health Sciences Division, Fred Hutchinson Cancer Research Center, Seattle, WA, USA, ^13^Department of Epidemiology, Harvard T.H. Chan School of Public Health, Boston, MA, USA, ^14^Department of Molecular and Clinical Cancer Medicine, University of Liverpool, Liverpool, UK, ^15^Department of Medical Biosciences, Umeå University, Umeå, Sweden, ^16^Lunenfeld-Tanenbuaum Research Institute, Sinai Health System, Toronto, ON, Canada, ^17^Department of Radiation Sciences, Umeå University, Umeå, Sweden, ^18^Radboud University Medical Center, Nijmegen, The Netherlands, ^19^British Columbia Cancer Agency, Vancouver, BC, Canada, ^20^College of Pharmacy, Washington State University, Spokane, WA, USA, ^21^Princess Margaret Cancer Center, Toronto, ON, Canada, ^22^Epidemiology Program, University of Hawaii Cancer Center, Honolulu, HI, USA, ^23^Department of Clinical Sciences Malmö, Lund University, Lund, Sweden, ^24^Department of Internal Medicine, Skåne University Hospital, Malmö, Sweden, ^25^Carmel Medical Center and Technion Faculty of Medicine, Haifa, Israel, ^26^University of Salzburg and Cancer Cluster, Salzburg, Germany, ^27^Translational Lung Research Center Heidelberg (TLRC-H), German Center for Lung Research (DZL), Heidelberg, Germany, ^28^German Cancer Research Center (DKFZ), Heidelberg, Germany, ^29^Department of Cancer Epidemiology, H. Lee Moffitt Cancer Center and Research Institute, Tampa, FL, USA, ^30^Department of Epidemiology and Biostatistics, Jiangsu Key Lab of Cancer Biomarkers, Prevention and Treatment, Collaborative Innovation Center for Cancer Personalized Medicine, School of Public Health, Nanjing, P.R. China, ^31^Department of Epidemiology, The University of Texas MD Anderson Cancer Center, Houston, TX, USA, ^32^Faculty of Medicine, IUOPA, University of Oviedo and CIBERESP, Oviedo, Spain, ^33^School of Health and Related Research (ScHARR), University Of Sheffield, Sheffield, UK, ^34^Institute of Medical Informatics, Biometry and Epidemiology, Chair of Epidemiology, Ludwig Maximilians University, Munich, Germany, ^35^Helmholtz Center Munich, Institute of Epidemiology, Neuherberg, Germany, ^36^Institute of Medical Statistics and Epidemiology, Technical University Munich, Munich, Germany, ^37^National Institute of Occupational Health (STAMI), Oslo, Norway

**InterLymph**

Demetrius Albanes^1^, Yolanda Benavente^2,3^, Paige M. Bracci^4^, Angela R. Brooks-Wilson^5,6^, Neil E. Caporaso^1^, James Cerhan^7^, Jacqueline Clavel^8,9^, Pierluigi Cocco^10^, Silvia de Sanjose^2,3^, Graham G. Giles^11,12^, Henrik Hjalgrim^13^, Rebecca D. Jackson^14^, Eleanor Kane^15^, Qing Lan^1^, Brian K. Link^16^, Alain Monnereau^8,9^, Alexandra Nieters^17^, Kari E. North^18,19^, Kenneth Offit^20^, Elio Riboli^21^, Christine F. Skibola^22^, Karin E. Smedby^23,24^, John J. Spinelli^25,26^, Lauren R. Teras^27^, Lesley F. Tinker^28^, Claire M. Vajdic^29^, Roel C.H. Vermeulen^30,31^, Joseph Vijai^20^, Paolo Vineis^32,33^, Anne Zeleniuch-Jacquotte^34,35,36^, Yawei Zhang^37^

^1^Division of Cancer Epidemiology and Genetics, National Cancer Institute, National Institutes of Health, Rockville, MD, USA, ^2^Cancer Epidemiology Research Programme, Catalan Institute of Oncology-IDIBELL, L'Hospitalet de Llobregat, Barcelona, Spain, ^3^CIBER de Epidemiología y Salud Pública (CIBERESP), Barcelona, Spain, ^4^Department of Epidemiology and Biostatistics, University of California San Francisco, San Francisco, CA, USA, ^5^Genome Sciences Centre, BC Cancer Agency, Vancouver, British Columbia, Canada, ^6^Department of Biomedical Physiology and Kinesiology, Simon Frazer University, Burnaby, British Columbia, Canada, ^7^Department of Internal Medicine, Mayo Clinic, Rochester, MN, USA, ^8^Epidemiology of Childhood and Adolescent Cancers Group, Inserm, Center of Research in Epidemiology and Statistics Sorbonne Paris Cité (CRESS), Paris, France, ^9^Université Paris Descartes, Paris, France, ^10^Department of Public Health, Clinical and Molecular Medicine, University of Cagliari, Monserrato, Cagliari, Italy, ^11^Cancer Epidemiology & Intelligence Division, Cancer Council Victoria, Melbourne, Victoria, Australia, ^12^Centre for Epidemiology and Biostatistics, Melbourne School of Population and Global Health,, University of Melbourne, Melbourne, Victoria, Australia, ^13^Department of Epidemiology Research, Division of Health Surveillance and Research, Statens Serum Institut, Copenhagen, Denmark, ^14^Division of Endocrinology, Diabetes and Metabolism, The Ohio State University, Columbus, OH, USA, ^15^Department of Health Sciences, University of York, York, UK, ^16^Department of Internal Medicine, Carver College of Medicine, The University of Iowa, Iowa City, IA, USA, ^17^Center for Chronic Immunodeficiency, University Medical Center Freiburg, Freiburg, Baden-Württemberg, Germany, ^18^Department of Epidemiology, University of North Carolina at Chapel Hill, Chapel Hill, NC, USA, ^19^Carolina Center for Genome Sciences, University of North Carolina at Chapel Hill, Chapel Hill, NC, USA, ^20^Department of Medicine, Memorial Sloan Kettering Cancer Center, New York, NY, USA, ^21^School of Public Health, Imperial College London, London, UK, ^22^Department of Hematology and Medical Oncology, Emory University School of Medicine, Atlanta, GA, USA, ^23^Department of Medicine, Solna, Karolinska Institutet, Stockholm, Sweden, ^24^Hematology Center, Karolinska University Hospital, Stockholm, Sweden, ^25^Cancer Control Research, BC Cancer Agency, Vancouver, British Columbia, Canada, ^26^School of Population and Public Health, University of British Columbia, Vancouver, British Columbia, Canada, ^27^Epidemiology Research Program, American Cancer Society, Atlanta, GA, USA, ^28^Division of Public Health Sciences, Fred Hutchinson Cancer Research Center, Seattle, WA, USA, ^29^Centre for Big Data Research in Health, University of New South Wales, Sydney, New South Wales, Australia, ^30^Institute for Risk Assessment Sciences, Utrecht University, Utrecht, The Netherlands, ^31^Julius Center for Health Sciences and Primary Care, University Medical Center Utrecht, Utrecht, The Netherlands, ^32^MRC-PHE Centre for Environment and Health, School of Public Health, Imperial College London, London, UK, ^33^Human Genetics Foundation, Turin, Italy, ^34^Department of Population Health, New York University School of Medicine, New York, NY, USA, ^35^Department of Environmental Medicine, New York University School of Medicine, New York, NY, USA, ^36^Perlmutter Cancer Center, NYU Langone Medical Center, New York, NY, USA, ^37^Department of Environmental Health Sciences, Yale School of Public Health, New Haven, CT, USA

**OCAC**

Katja K.H. Aben^1,2^, AOCS Group^3,4^, Hoda Anton-Culver^5^, Natalia N. Antonenkova^6^, Gerassimos Aravantinos^7^, Susana N. Banerjee^8^, Yukie Bean^9^, Matthias W. Beckmann^10^, Alicia Beeghly-Fadiel^11^, Javier Benitez^12,13^, Marina Bermisheva^14^, Marcus Q. Bernardini^15^, Line Bjorge^16,17^, Amanda Black^18^, Clara Bodelon^18^, Natalia V. Bogdanova^19,20,6^, James D. Brenton^21^, Louise Brinton^18^, Per Broberg^22^, Angela Brooks-Wilson^23,24^, Fiona Bruinsma^25^, Ralf Butzow^26^, Ian Campbell^27,28^, Rikki Cannioto^29^, Michael E. Carney^30^, Jenny Chang-Claude^31,32^, Stephen J. Chanock^18^, Xiao Qing Chen^4^, Georgia Chenevix-Trench^4^, Yoke-Eng Chiew^33,34^, Linda S. Cook^35^, Daniel W. Cramer^36,37^, Julie M. Cunningham^38^, Aimee A. D'Aloisio^39^, Agnieszka Dansonka-Mieszkowska^40^, Fanny Dao^41^, Anna deFazio^33,34^, Joe Dennis^42^, Ed Dicks^43^, Jennifer Anne Doherty^44^, Thilo Dörk^20^, Laure Dossus^45^, Matthias Dürst^46^, Diana M. Eccles^47^, Todd Edwards^48^, Arif B. Ekici^49^, Ailith Ewing^42^, Peter A. Fasching^50,10^, Sarah Ferguson^51^, James M. Flanagan^52^, Zachary C. Fogarty^53^, Renée T. Fortner^31^, Florentia Fostira^54^, George Fountzilas^55^, María J. García^56,57^, Aleksandra Gentry-Maharaj^58^, Graham G. Giles^59,60,61^, Rosalind Glasspool^62^, Marc T. Goodman^63^, Teodora Goranova^21^, Jacek Gronwald^64^, OPAL Study Group^65^, Christopher A. Haiman^66^, Niclas Håkansson^67^, Holly R. Harris^68,69^, Dennis Hazelett^70^, Alexander Hein^10^, Michelle A.T. Hildebrandt^71^, Peter Hillemanns^20^, Claus K. Høgdall^72^, Estrid Høgdall^73,74^, Helene Holland^4^, Karen Hosking^75^, Ruea-Yea Huang^76^, David G. Huntsman^77,78,79,80^, Tomasz Huzarski^64^, Liher Imaz^81,82^, Anna Jakubowska^64,83^, Allan Jensen^73^, Sharon Johnatty^4^, Michael E. Jones^84^, Pääivy Kannistö^85^, Siddhartha Kar^43^, Beth Y. Karlan^86,87^, Anthony Karnezis^88^, Linda E. Kelemen^89^, Catherine J. Kennedy^33,34^, Elza Khusnutdinova^90,14^, Lambertus A. Kiemeney^1^, Susanne K. Kjaer^73,91^, Martin Köbel^92^, Reidun K. Kopperud^16,17^, Jolanta Kupryjanczyk^40^, Diether Lambrechts^93,94^, Melissa C. Larson^53^, Kate Lawrenson^95^, Nhu D. Le^96^, Loic Le Marchand^97^, Shashikant B. Lele^98^, Jenny Lester^86,87^, Douglas A. Levine^41,99^, Dong Liang^100^, Clemens Liebrich^101^, Loren Lipworth^102^, Jolanta Lissowska^103^, Karen H. Lu^104^, Jan Lubiński^64^, Lene Lundvall^72^, Leon F.A.G. Massuger^105^, Taymaa May^106^, Jessica McAlpine^107^, Valerie McGuire^108^, John R. McLaughlin^109^, Iain A. McNeish^110,111^, Usha Menon^58^, Melissa Merritt^112,113^, Francesmary Modugno^114,115^, Melissa Moffitt^9,116^, Alvaro N. Monteiro^117^, Steven A. Narod^118^, Lotte Nedergaard^119^, Roberta B. Ness^120^, Heli Nevanlinna^121^, Kunle Odunsi^98^, Siel Olbrecht^122^, Håkan Olsson^22^, N. Charlotte Onland-Moret^123^, Nick Orr^124^, Sandra Orsulic^87^, Ana Osorio^12,56^, Domenico Palli^125^, Sue K. Park^126,127,128^, Tjoung-Won Park-Simon^20^, James Paul^129^, Tanja Pejovic^9,116^, Liisa M. Pelttari^130^, Jennifer B. Permuth^117^, Malcolm C. Pike^131,132^, Anna Piskorz^21^, Joanna Plisiecka-Halasa^40^, Darya Prokofyeva^90^, Susan J. Ramus^133,134^, Marjorie J. Riggan^135^, Cristina Rodriguez-Antona^13,57^, Mary Anne Rossing^68,69^, Joseph H. Rothstein^136,137^, Ingo Runnebaum^46^, Dale P. Sandler^138^, Minouk J. Schoemaker^84^, V. Wendy Setiawan^139^, Gianluca Severi^140,141,142,143^, Nadeem Siddiqui^144^, Weiva Sieh^137,136^, Honglin Song^43^, Melissa C. Southey^61,145^, Lara Sucheston^146^, Rebecca Sutphen^147^, Anthony J. Swerdlow^84,148^, Lukasz Szafron^149^, Jack A. Taylor^138,150^, Soo H. Teo^151,152^, Kathryn L. Terry^36,37^, Liv Cecilie Vestrheim Thomsen^16,17^, Anne Tinker^153^, Linda Titus^154^, Alicia Tone^106^, Britton Trabert^18^, Ruth Travis^155^, Antonia Trichopoulou^156,157^, Jonathan P. Tyrer^43^, Shelley S. Tworoger^117,36^, Anne M. van Altena^158^, David Van Den Berg^139^, Els Van Nieuwenhuysen^122^, Digna R. Velez Edwards^159^, Ignace Vergote^122^, Robert A. Vierkant^53^, Christine Walsh^87^, Shan Wang-Gohrke^160^, Penelope M. Webb^65^, Clarice R. Weinberg^161^, Nicolas Wentzensen^18^, Alice S. Whittemore^108,162^, Lynne R. Wilkens^163^, Stacey J. Winham^53^, Alicja Wolk^67,164^, Michelle Woo^165^, Xifeng Wu^71^, Anna H. Wu^139^, Hannah P. Yang^18^, Drakoulis Yannoukakos^54^, Argyrios Ziogas^5^

^1^Radboud Institute for Health Sciences, Radboud University Medical Center, Nijmegen, The Netherlands, ^2^Netherlands Comprehensive Cancer Organisation, Utrecht, The Netherlands, ^3^Research Department, Peter MacCallum Cancer Center, Melbourne, Victoria, Australia, ^4^Department of Genetics and Computational Biology, QIMR Berghofer Medical Research Institute, Brisbane, Queensland, Australia, ^5^Department of Epidemiology, Genetic Epidemiology Research Institute, University of California Irvine, Irvine, CA, USA, ^6^N.N. Alexandrov Research Institute of Oncology and Medical Radiology, Minsk, Belarus, ^7^"Agii Anargiri" Cancer Hospital, Athens, Greece, ^8^Gynaecology Unit, Royal Marsden Hospital, London, UK, ^9^Department of Obstetrics and Gynecology, Oregon Health & Science University, Portland, OR, USA, ^10^Department of Gynecology and Obstetrics, Comprehensive Cancer Center ER-EMN, University Hospital Erlangen, Friedrich-Alexander-University Erlangen-Nuremberg, Erlangen, Germany, ^11^Department of Medicine, Vanderbilt Epidemiology Center, Vanderbilt-Ingram Cancer Center, Division of Epidemiology, Vanderbilt University School of Medicine, Nashville, TN, USA, ^12^Centro de Investigación en Red de Enfermedades Raras (CIBERER), Madrid, Spain, ^13^Human Cancer Genetics Programme, Spanish National Cancer Research Centre (CNIO), Madrid, Spain, ^14^Institute of Biochemistry and Genetics, Ufa Federal Research Centre of the Russian Academy of Sciences, Ufa, Russia, ^15^University Health Network, Division of Gynecologic Oncology, Princess Margaret Hospital, Toronto, Ontario, Canada, ^16^Department of Obstetrics and Gynecology, Haukeland University Hospital, Bergen, Norway, ^17^Department of Clinical Science, Centre for Cancer Biomarkers CCBIO, University of Bergen, Bergen, Norway, ^18^Division of Cancer Epidemiology and Genetics, National Cancer Institute, National Institutes of Health, Rockville, MD, USA, ^19^Department of Radiation Oncology, Hannover Medical School, Hannover, Germany, ^20^Gynaecology Research Unit, Hannover Medical School, Hannover, Germany, ^21^Cancer Research UK Cambridge Institute, University of Cambridge, Cambridge, UK, ^22^Department of Cancer Epidemiology, Clinical Sciences, Lund University, Lund, Sweden, ^23^Genome Sciences Centre, BC Cancer Agency, Vancouver, BC, Canada, ^24^Department of Biomedical Physiology and Kinesiology, Simon Fraser University, Burnaby, BC, Canada, ^25^Cancer Epidemiology Division, Cancer Council Victoria, Melbourne, Victoria, Australia, ^26^Department of Pathology, Helsinki University Hospital, University of Helsinki, Helsinki, Finland, ^27^Research Department, Peter MacCallum Cancer Center, Melbourne, Victoria, Australia, ^28^Sir Peter MacCallum Department of Oncology, The University of Melbourne, Melbourne, Victoria, Australia, ^29^Cancer Pathology & Prevention, Division of Cancer Prevention and Population Sciences, Roswell Park Cancer Institute, Buffalo, NY, USA, ^30^Department of Obstetrics and Gynecology, John A. Burns School of Medicine, University of Hawaii, Honolulu, HI, USA, ^31^Division of Cancer Epidemiology, German Cancer Research Center (DKFZ), Heidelberg, Germany, ^32^Cancer Epidemiology Group, University Cancer Center Hamburg (UCCH), University Medical Center Hamburg-Eppendorf, Hamburg, Germany, ^33^Centre for Cancer Research, The Westmead Institute for Medical Research, The University of Sydney, Sydney, New South Wales, Australia, ^34^Department of Gynaecological Oncology, Westmead Hospital, Sydney, New South Wales, Australia, ^35^University of New Mexico Health Sciences Center, University of New Mexico, Albuquerque, NM, USA, ^36^Department of Epidemiology, Harvard T.H. Chan School of Public Health, Boston, MA, USA, ^37^Obstetrics and Gynecology Epidemiology Center, Brigham and Women's Hospital and Harvard Medical School, Boston, MA, USA, ^38^Department of Laboratory Medicine and Pathology, Mayo Clinic, Rochester, MN, USA, ^39^Social & Scientific Systems, Inc., Durham, NC, USA, ^40^Department of Pathology and Laboratory Medicine, Institute of Oncology and Maria Sklodowska-Curie Cancer Center, Warsaw, Poland, ^41^Department of Surgery, Gynecology Service, Memorial Sloan Kettering Cancer Center, New York, NY, USA, ^42^Department of Public Health and Primary Care, Centre for Cancer Genetic Epidemiology, University of Cambridge, Cambridge, UK, ^43^Department of Oncology, Centre for Cancer Genetic Epidemiology, University of Cambridge, Cambridge, UK, ^44^Department of Population Health Sciences, Huntsman Cancer Institute, University of Utah, Salt Lake City, UT, USA, ^45^Nutrition and Metabolism Section, International Agency for Research on Cancer (IARC-WHO), Lyon, France, ^46^Department of Gynaecology, Jena University Hospital - Friedrich Schiller University, Jena, Germany, ^47^Faculty of Medicine, University of Southampton, Southampton, UK, ^48^Department of Medicine, Division of Epidemiology, Center for Human Genetics Research, Vanderbilt University Medical Center, Nashville, TN, USA, ^49^Institute of Human Genetics, University Hospital Erlangen, Friedrich-Alexander University Erlangen-Nuremberg, Comprehensive Cancer Center Erlangen-EMN, Erlangen, Germany, ^50^Department of Medicine Division of Hematology and Oncology, David Geffen School of Medicine, University of California at Los Angeles, Los Angeles, CA, USA, ^51^Division of Gynecologic Oncology, University Health Network, Princess Margaret Hospital, Toronto, Ontario, Canada, ^52^Department of Surgery and Cancer, Division of Cancer and Ovarian Cancer Action Research Centre, Imperial College London, London, UK, ^53^Department of Health Science Research, Division of Biomedical Statistics and Informatics, Mayo Clinic, Rochester, MN, USA, ^54^Molecular Diagnostics Laboratory, INRASTES, National Centre for Scientific Research "Demokritos", Athens, Greece, ^55^Second Department of Medical Oncology, EUROMEDICA General Clinic of Thessaloniki, Aristotle University of Thessaloniki School of Medicine, Thessalonıki, Greece, ^56^Human Cancer Genetics Programme, Spanish National Cancer Research Centre (CNIO), Madrid, Spain, ^57^Biomedical Network on Rare Diseases (CIBERER), Madrid, Spain, ^58^MRC Clinical Trials Unit at UCL, Institute of Clinical Trials & Methodology, University College London, London, UK, ^59^Cancer Epidemiology Division, Cancer Council Victoria, Melbourne, Victoria, Australia, ^60^Centre for Epidemiology and Biostatistics, Melbourne School of Population and Global Health, The University of Melbourne, Melbourne, Victoria, Australia, ^61^Precision Medicine, School of Clinical Sciences at Monash Health, Monash University, Clayton, Victoria, Australia, ^62^Department of Medical Oncology, The Beatson West of Scotland Cancer Centre, Glasgow, UK, ^63^Samuel Oschin Comprehensive Cancer Institute, Cancer Prevention and Genetics Program, Cedars-Sinai Medical Center, Los Angeles, CA, USA, ^64^Department of Genetics and Pathology, Pomeranian Medical University, Szczecin, Poland, ^65^Population Health Department, QIMR Berghofer Medical Research Institute, Brisbane, Queensland, Australia, ^66^Department of Preventive Medicine, Keck School of Medicine, University of Southern California, Los Angeles, CA, USA, ^67^Institute of Environmental Medicine, Karolinska Institutet, Stockholm, Sweden, ^68^Program in Epidemiology, Division of Public Health Sciences, Fred Hutchinson Cancer Research Center, Seattle, WA, USA, ^69^Department of Epidemiology, University of Washington, Seattle, WA, USA, ^70^Center for Bioinformatics and Functional Biology, Samuel Oschin Comprehensive Cancer Institute, Cedars-Sinai Medical Center, Los Angeles, CA, USA, ^71^Department of Epidemiology, University of Texas MD Anderson Cancer Center, Houston, TX, USA, ^72^Department of Gynecology, Rigshospitalet, The Juliane Marie Centre, University of Copenhagen, Copenhagen, Denmark, ^73^Department of Virus, Lifestyle and Genes, Danish Cancer Society Research Center, Copenhagen, Denmark, ^74^Molecular Unit, Department of Pathology, Herlev Hospital, University of Copenhagen, Copenhagen, Denmark, ^75^Department of Oncology, University of Cambridge, Cambridge, UK, ^76^Center For Immunotherapy, Roswell Park Cancer Institute, Buffalo, NY, USA, ^77^Department of Molecular Oncology, BC Cancer Research Centre, Vancouver, BC, Canada, ^78^Department of Pathology and Laboratory Medicine, University of British Columbia, Vancouver, BC, Canada, ^79^Department of Obstetrics and Gynecology, University of British Columbia, Vancouver, BC, Canada, ^80^British Columbia's Ovarian Cancer Research (OVCARE) Program, Vancouver General Hospital, University of British Columbia, BC Cancer and Vancouver General Hospital, Vancouver, BC, Canada, ^81^Biodonostia Health Research Institute, Donostia-San Sebastian, Spain, ^82^Ministry of Health of the Basque Government Public Health Division of Gipuzkoa, Donostia-San Sebastian, Spain, Public Health Division of Gipuzkoa, Donostia-San Sebastian, Spain, ^83^Independent Laboratory of Molecular Biology and Genetic Diagnostics, Pomeranian Medical University, Szczecin, Poland, ^84^Division of Genetics and Epidemiology, The Institute of Cancer Research, London, UK, ^85^Department of Gynecology, University Hospital, Lund, Sweden, ^86^Department of Obstetrics and Gynecology, David Geffen School of Medicine, University of California at Los Angeles, Los Angeles, CA, USA, ^87^Women's Cancer Program at the Samuel Oschin Comprehensive Cancer Institute, Cedars-Sinai Medical Center, Los Angeles, CA, USA, ^88^Department of Pathology and Laboratory Medicine, UC Davis Medical Center, Sacramento, CA, USA, ^89^Hollings Cancer Center, Medical University of South Carolina, Charleston, SC, USA, ^90^Department of Genetics and Fundamental Medicine, Bashkir State Medical University, Ufa, Russia, ^91^Department of Gynaecology, Rigshospitalet, University of Copenhagen, Copenhagen, Denmark, ^92^Department of Pathology and Laboratory Medicine, University of Calgary, Foothills Medical Center, Calgary, AB, Canada, ^93^VIB Center for Cancer Biology, VIB, Leuven, Belgium, ^94^Laboratory for Translational Genetics, Department of Human Genetics, University of Leuven, Leuven, Belgium, ^95^Department of Obstetrics and Gynecology, Division of Gynecologic Oncology, Women’s Cancer Program at the Samuel Oschin Cancer Institute Cedars-Sinai Medical Center, Los Angeles, CA, USA, ^96^Cancer Control Research, BC Cancer Agency, Vancouver, BC, Canada, ^97^Epidemiology Program, University of Hawaii Cancer Center, Honolulu, HI, USA, ^98^Department of Gynecologic Oncology, Roswell Park Cancer Institute, Buffalo, NY, USA, ^99^Gynecologic Oncology, Laura and Isaac Pearlmutter Cancer Center, NYU Langone Medical Center, New York, NY, USA, ^100^College of Pharmacy and Health Sciences, Texas Southern University, Houston, TX, USA, ^101^Clinics of Gynaecology, Cancer Center Wolfsburg, Wolfsburg, Germany, ^102^Department of Medicine, Division of Epidemiology, Vanderbilt University Medical Center, Nashville, TN, USA, ^103^Department of Cancer Epidemiology and Prevention, M. Sklodowska-Curie Cancer Center, Oncology Institute, Warsaw, Poland, ^104^Department of Gynecologic Oncology and Clinical Cancer Genetics Program, University of Texas MD Anderson Cancer Center, Houston, TX, USA, ^105^Department of Gynaecology, Radboud Institute for Molecular Life Sciences, Radboud University Medical Center, Nijmegen, The Netherlands, ^106^Division of Gynecologic Oncology, University Health Network, Princess Margaret Hospital, Toronto, Ontario, Canada, ^107^Department of Gynecology, Division Gynecologic Oncology, University of British Columbia and BC Cancer Agency, Vancouver, BC, Canada, ^108^Department of Health Research and Policy - Epidemiology, Stanford University School of Medicine, Stanford, CA, USA, ^109^Public Health Ontario, Samuel Lunenfeld Research Institute, Toronto, Ontario, Canada, ^110^Department Surgery & Cancer, Division of Cancer and Ovarian Cancer Action Research Centre, Imperial College London, London, UK, ^111^Institute of Cancer Sciences, University of Glasgow, Glasgow, UK, ^112^Department of Epidemiology and Biostatistics, School of Public Health, Imperial College London, London, UK, ^113^Epidemiology Program, University of Hawaii Cancer Center, Honolulu, HI, USA, ^114^Womens Cancer Research Center, Magee-Womens Research Institute and Hillman Cancer Center, Pittsburgh, PA, USA, ^115^Department of Obstetrics, Gynecology and Reproductive Sciences, Division of Gynecologic Oncology, University of Pittsburgh School of Medicine, Pittsburgh, PA, USA, ^116^Knight Cancer Institute, Oregon Health & Science University, Portland, OR, USA, ^117^Department of Cancer Epidemiology, Moffitt Cancer Center, Tampa, FL, USA, ^118^Women's College Research Institute, University of Toronto, Toronto, Ontario, Canada, ^119^Department of Pathology, Rigshospitalet, University of Copenhagen, Copenhagen, Denmark, ^120^School of Public Health, University of Texas Health Science Center at Houston (UTHealth), Houston, TX, USA, ^121^Department of Obstetrics and Gynecology, Helsinki University Hospital, University of Helsinki, Helsinki, Finland, ^122^Department of Obstetrics and Gynaecology and Leuven Cancer Institute, Division of Gynecologic Oncology, University Hospitals Leuven, Leuven, Belgium, ^123^Julius Center for Health Sciences and Primary Care, University Utrecht, UMC Utrecht, Utrecht, The Netherlands, ^124^Centre for Cancer Research and Cell Biology, Queen's University Belfast, Belfast, UK, ^125^Cancer Risk Factors and Life-Style Epidemiology Unit, Institute for Cancer Research, Prevention and Clinical Network (ISPRO), Florence, Italy, ^126^Department of Preventive Medicine, Seoul National University College of Medicine, Seoul, Korea, ^127^Department of Biomedical Sciences, Seoul National University Graduate School, Seoul, Korea, ^128^Cancer Research Institute, Seoul National University, Seoul, Korea, ^129^Cancer Research UK Clinical Trials Unit, University of Glasgow, Glasgow, UK, ^130^Department of Obstetrics and Gynecology, University of Helsinki and Helsinki University Hospital, Helsinki, Finland, ^131^Department of Epidemiology and Biostatistics, Memorial Sloan-Kettering Cancer Center, New York, NY, USA, ^132^Department of Preventive Medicine, Keck School of Medicine, University of Southern California Norris Comprehensive Cancer Center, Los Angeles, CA, USA, ^133^School of Women's and Children's Health, Faculty of Medicine, University of NSW Sydney, Sydney, New South Wales, Australia, ^134^The Kinghorn Cancer Centre, Garvan Institute of Medical Research, Sydney, New South Wales, Australia, ^135^Department of Gynecologic Oncology, Duke University Medical Center, Durham, NC, USA, ^136^Department of Genetics and Genomic Sciences, Icahn School of Medicine at Mount Sinai, New York, NY, USA, ^137^Department of Health Science and Policy, Icahn School of Medicine at Mount Sinai, New York, NY, USA, ^138^Epidemiology Branch, National Institute of Environmental Health Sciences, NIH, Research Triangle Park, NC, USA, ^139^Department of Preventive Medicine, Keck School of Medicine, University of Southern California, Los Angeles, CA, USA, ^140^Human Genetics Foundation (HuGeF), Turino, Italy, ^141^Cancer Council Victoria and University of Melbourne, Melbourne, Victoria, Australia, ^142^Gustave Roussy, Villejuif, France, ^143^Université Paris-Saclay, Université Paris-Sud, Villejuif, France, ^144^Department of Gynaecological Oncology, Glasgow Royal Infirmary, Glasgow, UK, ^145^Department of Clinical Pathology, The University of Melbourne, Melbourne, Victoria, Australia, ^146^Division of Cancer Prevention and Control, Roswell Park Cancer Institute, Buffalo, NY, USA, ^147^Epidemiology Center, College of Medicine, University of South Florida, Tampa, FL, USA, ^148^Division of Breast Cancer Research, The Institute of Cancer Research, London, UK, ^149^Department of Immunology, the Maria Sklodowska-Curie Institute - Oncology Center, Warsaw, Poland, ^150^Epigenetic and Stem Cell Biology Laboratory, National Institute of Environmental Health Sciences, NIH, Research Triangle Park, NC, USA, ^151^Breast Cancer Research Programme, Cancer Research Malaysia, Subang Jaya, Selangor, Malaysia, ^152^Department of Surgery, Faculty of Medicine, University Malaya, Kuala Lumpur, Malaysia, ^153^British Columbia's Ovarian Cancer Research (OVCARE) Program - Cheryl Brown Ovarian Cancer Outcomes Unit (CBOCOU), BC Cancer Agency, Vancouver, BC, Canada, ^154^Geisel School of Medicine, Dartmouth College, Hanover, NH, USA, ^155^Cancer Epidemiology Unit, University of Oxford, Oxford, UK, ^156^Hellenic Health Foundation, Athens, Greece, ^157^WHO Collaborating Center for Nutrition and Health, Unit of Nutritional Epidemiology and Nutrition in Public Health, Dept. of Hygiene, Epidemiology and Medical Statistics, University of Athens Medical School, Athens, Greece, ^158^Department of Gynaecology, Radboud University Medical Center, Nijmegen, The Netherlands, ^159^Division of Quantitative Sciences, Department of Obstetrics and Gynecology, Department of Biomedical Informatics, Vanderbilt Epidemiology Center, Vanderbilt Genetics Institute, Vanderbilt University Medical Center, Nashville, TN, USA, ^160^Department of Gynaecology and Obstetrics, University Hospital Ulm, Ulm, Germany, ^161^Biostatistics and Computational Biology Branch, National Institute of Environmental Health Sciences, NIH, Research Triangle Park, NC, USA, ^162^Department of Biomedical Data Science, Stanford University School of Medicine, Stanford, CA, USA, ^163^Cancer Epidemiology Program, University of Hawaii Cancer Center, Honolulu, HI, USA, ^164^Department of Surgical Sciences, Uppsala University, Uppsala, Sweden, ^165^Department of Obstetrics and Gynecology, British Columbia's Ovarian Cancer Research (OVCARE) Program, Vancouver General Hospital, University of British Columbia, Vancouver, BC, Canada

**PanScan, PANC4**

Demetrius Albanes^1^, Gabriella Andreotti^1^, Alan A. Arslan^2,3,4^, Ana Babic^5^, Laura Beane-Freeman^1^, Julie Buring^6,7^, Federico Canzian^8^, Stephen J. Chanock^1^, Eric J. Duell^9^, Charles Fuchs^10^, J. Michael Gaziano^6,11,12^, Graham G. Giles^13,14,15^, Edward Giovannucci^5^, Gary E. Goodman^16^, Phyllis J. Goodman^17^, Patricia Hartge^1^, Robert Hoover^1^, Rudolf Kaaks^18^, Kay-Tee Khaw^19^, Eric A. Klein^20^, Manolis Kogevinas^21,22,23,24^, Charles Kooperberg^16^, Peter Kraft^7,25^, Loic Le Marchand^26^, Núria Malats^27^, Satu Männistö^28^, Olle Melander^29^, Roger Milne^13,14,30^, Kimmie Ng^5^, Domenico Palli^31^, Alpa V. Patel^32^, Ulrike Peters^16^, Miquel Porta^22,23^, Elio Riboli^33^, Maria-Jose Sanchez^34,35,22^, Howard D. Sesso^6,7^, Xiao-Ou Shu^36^, Mark D. Thornquist^16^, Anne Tjønneland^37,38^, Geoffrey S. Tobias^1^, Ruth C. Travis^39^, Antonia Trichopoulou^40^, Thérèse Truong^41^, Roel C.H. Vermeulen^42,43^, Kala Visvanathan^44^, Jean Wactawski-Wende^45^, Elisabete Weiderpass^46^, Emily White^16,47^, Lynne R. Wilkens^26^, Herbert Yu^26^, Kai Yu^1^, Chen Yuan^5^, Anne Zeleniuch-Jacquotte^3,48^, Wei Zheng^36^, Jun Zhong^49^

^1^Division of Cancer Epidemiology and Genetics, National Cancer Institute, National Institutes of Health, Rockville, MD, USA, ^2^Department of Obstetrics and Gynecology, New York University School of Medicine, New York, NY, USA, ^3^Department of Population Health, New York University School of Medicine, New York, NY, USA, ^4^Department of Environmental Medicine, New York University School of Medicine, New York, NY, USA, ^5^Department of Medical Oncology, Dana-Farber Cancer Institute, Boston, MA, USA, ^6^Division of Preventive Medicine, Brigham and Women’s Hospital, Boston, MA, USA, ^7^Department of Epidemiology, Harvard T.H. Chan School of Public Health, Boston, MA, USA, ^8^Genomic Epidemiology Group, German Cancer Research Center (DKFZ), Heidelberg, Germany, ^9^Unit of Nutrition and Cancer, Cancer Epidemiology Research Program, Bellvitge Biomedical Research Institute (IDIBELL) , Catalan Institute of Oncology (ICO), Barcelona, Spain, ^10^Yale Cancer Center, New Haven, CT, USA, ^11^Division of Aging, Brigham and Women’s Hospital, Boston, MA, USA, ^12^Boston VA Healthcare System, Boston, MA, USA, ^13^Cancer Epidemiology and Intelligence Division, Cancer Council Victoria, Melbourne, VIC, Australia, ^14^Centre for Epidemiology and Biostatistics, Melbourne School of Population and Global Health, The University of Melbourne, Parkville, VIC, Australia, ^15^Department of Epidemiology and Preventive Medicine, Monash University, Melbourne, VIC, Australia, ^16^Division of Public Health Sciences, Fred Hutchinson Cancer Research Center, Seattle, WA, USA, ^17^SWOG Statistical Center, Fred Hutchinson Cancer Research Center, Seattle, WA, USA, ^18^Division of Cancer Epidemiology, German Cancer Research Center (DKFZ), Heidelberg, Germany, ^19^Department of Public Health and Primary Care, Institute of Public Health, School of Clinical Medicine, University of Cambridge, Cambridge, UK, ^20^Glickman Urological and Kidney Institute, Cleveland Clinic, Cleveland, OH, USA, ^21^ISGlobal, Centre for Research in Environmental Epidemiology (CREAL), Barcelona, Spain, ^22^CIBER Epidemiología y Salud Pública (CIBERESP), Barcelona, Spain, ^23^Hospital del Mar Institute of Medical Research (IMIM), Universitat Autònoma de Barcelona, Barcelona, Spain, ^24^Universitat Pompeu Fabra (UPF), Barcelona, Spain, ^25^Department of Biostatistics, Harvard T.H. Chan School of Public Health, Boston, MA, USA, ^26^Cancer Epidemiology Program, University of Hawaii Cancer Center, Honolulu, HI, USA, ^27^Genetic and Molecular Epidemiology Group, Spanish National Cancer Research Center (CNIO), Madrid, Spain, ^28^Department of Public Health Solutions, National Institute for Health and Welfare, Helsinki, Finland, ^29^Department of Clinical Sciences Malmö, Lund University, Malmö, Sweden, ^30^Precision Medicine, School of Clinical Sciences at Monash Health, Monash University, Melbourne, VIC, Australia, ^31^Cancer Risk Factors and Life-Style Epidemiology Unit, Institute for Cancer Research, Prevention and Clinical Network - ISPRO, Villa delle Rose, Firenze, Italy, ^32^Epidemiology Research Program, American Cancer Society, Atlanta, GA, USA, ^33^Department of Epidemiology and Biostatistics, School of Public Health, Imperial College London, London, UK, ^34^Andalusian School of Public Health (EASP), Granada, Spain, ^35^Instituto de Investigación Biosanitaria de Granada (ibs.GRANADA), Universidad de Granada, Granada, Spain, ^36^Division of Epidemiology, Department of Medicine, Vanderbilt Epidemiology Center, Vanderbilt-Ingram Cancer Center, Vanderbilt University Medical Center, Nashville, TN, USA, ^37^Diet, Genes and Environment, Danish Cancer Society Research Center, Copenhagen, Denmark, ^38^Department of Public Health, Faculty of Health and Medical Sciences, University of Copenhagen, Copenhagen, Denmark, ^39^Cancer Epidemiology Unit, Nuffield Department of Health, University of Oxford, Oxford, UK, ^40^Hellenic Health Foundation, Athens, Greece, ^41^INSERM U1018 - Center for Research in Epidemiology and Population Health (CESP), Paris-Saclay University, Paris-Sud University, Villejuif, France, ^42^Institute for Risk Assessment Sciences (IRAS), Division of Environmental Epidemiology (EEPI), Utrecht University, Utrecht, The Netherlands, ^43^Julius Center for Health Sciences and Primary Care, University Medical Center Utrech, Utrecht, The Netherlands, ^44^Department of Epidemiology, Johns Hopkins Bloomberg School of Public Health, Baltimore, MD, USA, ^45^Department of Epidemiology and Environmental Health, University at Buffalo, Buffalo, NY, USA, ^46^International Agency for Research on Cancer, Lyon, France, ^47^Department of Epidemiology, University of Washington, Seattle, WA, USA, ^48^Perlmutter Cancer Center, New York University School of Medicine, New York, NY, USA, ^49^Laboratory of Translational Genomics, Division of Cancer Epidemiology and Genetics, National Cancer Institute, National Institutes of Health, Rockville, MD, USA

**PRACTICAL, CRUK, BPC3, CAPS, PEGASUS**

Demetrius Albanes^1^, Australian Prostate Cancer BioResource (APCB)^2^, Jyotsna Batra^2,3^, Sara Benlloch^4^, Sonja I. Berndt^1^, William J. Blot^5,6^, Hermann Brenner^7,8,9^, Géraldine Cancel-Tassin^10,11^, Lisa Cannon-Albright^12,13^, Stephen Chanock^1^, Frank Claessens^14^, Judith Clements^2,3^, David V. Conti^15^, Cezary Cybulski^16^, Kim De Ruyck^17^, Jenny L. Donovan^18^, Manuela Gago-Dominguez^19,20^, Susan M. Gapstur^21^, Graham G. Giles^22,23,24^, Eli Marie Grindedal^25^, Henrik Gronberg^26^, Freddie C. Hamdy^27,28^, Robert J. Hamilton^29,30^, Brian E. Henderson^15^, David J. Hunter^31^, Sue A. Ingles^15^, Esther M. John^32^, Radka Kaneva^33^, Kay-Tee Khaw^34^, Adam S. Kibel^35^, Jeri Kim^36^, Manolis Kogevinas^37,38,39,40^, Stella Koutros^1^, Peter Kraft^31^, Davor Lessel^41^, Yong-Jie Lu^42^, Christiane Maier^43^, Florence Menegaux^44^, Lorelei Mucci^45^, Kenneth Muir^46,47^, David E. Neal^48,49,50^, Susan L. Neuhausen^51^, Lisa F. Newcomb^52,53^, Børge G. Nordestgaard^54,55^, Hardev Pandha^56^, Jong Y. Park^57^, Nora Pashayan^58,59^, Kathryn L. Penney^60^, Azad Razack^61^, Elio Riboli^62^, Monique J. Roobol^63^, Barry S. Rosenstein^64,65^, Johanna Schleutker^66,67^, Karina Dalsgaard Sørensen^68,69^, Janet L. Stanford^52,70^, Victoria L. Stevens^21^, Catherine M. Tangen^71^, Manuel R. Teixeira^72,73^, Stephen N. Thibodeau^74^, Paul A. Townsend^75^, Ruth C. Travis^76^, Nawaid Usmani^77,78^, Ana Vega^79^, Stephanie Weinstein^1^, Catharine West^80^, Fredrik Wiklund^26^, Alicja Wolk^81,82^

^1^Division of Cancer Epidemiology and Genetics, National Cancer Institute, National Institutes of Health, Rockville, MD, USA, ^2^Australian Prostate Cancer Research Centre-Qld, Institute of Health and Biomedical Innovation and School of Biomedical Sciences, Queensland University of Technology, Brisbane, Queensland, Australia, ^3^Translational Research Institute, Brisbane, Queensland, Australia, ^4^Department of Public Health and Primary Care, Centre for Cancer Genetic Epidemiology, University of Cambridge, Cambridge, UK, ^5^Department of Medicine, Division of Epidemiology, Vanderbilt University Medical Center, Nashville, TN, USA, ^6^International Epidemiology Institute, Rockville, MD, USA, ^7^Division of Clinical Epidemiology and Aging Research, German Cancer Research Center (DKFZ), Heidelberg, Germany, ^8^Division of Preventive Oncology, German Cancer Research Center (DKFZ) and National Center for Tumor Diseases (NCT), Heidelberg, Germany, ^9^German Cancer Consortium (DKTK), German Cancer Research Center (DKFZ), Heidelberg, Germany, ^10^CeRePP, Tenon Hospital, Paris, France, ^11^Sorbonne Universite, GRC n°5, ONCOTYPE-URO, Tenon Hospital, Paris, France, ^12^Department of Medicine, Division of Genetic Epidemiology, University of Utah School of Medicine, Salt Lake City, UT, USA, ^13^George E. Wahlen Department of Veterans Affairs Medical Center, Salt Lake City, UT, USA, ^14^Department of Cellular and Molecular Medicine, Molecular Endocrinology Laboratory, Leuven, Belgium, ^15^Department of Preventive Medicine, Keck School of Medicine, University of Southern California/Norris Comprehensive Cancer Center, Los Angeles, CA, USA, ^16^Department of Genetics and Pathology, International Hereditary Cancer Center, Pomeranian Medical University, Szczecin, Poland, ^17^Faculty of Medicine and Health Sciences, Ghent University, Gent, Belgium, ^18^School of Social and Community Medicine, University of Bristol, Bristol, UK, ^19^Genomic Medicine Group, Galician Foundation of Genomic Medicine, Instituto de Investigacion Sanitaria de Santiago de Compostela (IDIS), Complejo Hospitalario Universitario de Santiago, Santiago de Compostela, Spain, ^20^Moores Cancer Center, University of California San Diego, San Diego, CA, USA, ^21^Epidemiology Research Program, American Cancer Society, Atlanta, GA, USA, ^22^Cancer Epidemiology Division, Cancer Council Victoria, Melbourne, Victoria, Australia, ^23^Centre for Epidemiology and Biostatistics, Melbourne School of Population and Global Health, The University of Melbourne, Melbourne, Victoria, Australia, ^24^Precision Medicine, School of Clinical Sciences at Monash Health, Monash University, Clayton, Victoria, Australia, ^25^Department of Medical Genetics, Oslo University Hospital, Oslo, Norway, ^26^Department of Medical Epidemiology and Biostatistics, Karolinska Institute, Stockholm, Sweden, ^27^Nuffield Department of Surgical Sciences, University of Oxford, Oxford, UK, ^28^Faculty of Medical Science, University of Oxford, John Radcliffe Hospital, Oxford, UK, ^29^Department of Surgical Oncology, Princess Margaret Cancer Centre, Toronto, Canada, ^30^Department of Surgery (Urology), University of Toronto, Toronto, Canada, ^31^Department of Epidemiology, Program in Genetic Epidemiology and Statistical Genetics, Harvard School of Public Health, Boston, MA, USA, ^32^Department of Medicine, Division of Oncology, Stanford Cancer Institute, Stanford University School of Medicine, Stanford, CA, USA, ^33^Department of Medical Chemistry and Biochemistry, Molecular Medicine Center, Medical University of Sofia, Sofia, Bulgaria, ^34^Clinical Gerontology Unit Department of Public Health and Primary Care, University of Cambridge, Cambridge, UK, ^35^Division of Urologic Surgery, Brigham and Womens Hospital, Boston, MA, USA, ^36^Department of Genitourinary Medical Oncology, The University of Texas MD Anderson Cancer Center, Houston, TX, USA, ^37^ISGlobal, Barcelona, Spain, ^38^IMIM (Hospital del Mar Medical Research Institute), Barcelona, Spain, ^39^Universitat Pompeu Fabra (UPF), Barcelona, Spain, ^40^CIBER Epidemiología y Salud Pública (CIBERESP), Madrid, Spain, ^41^Institute of Human Genetics, University Medical Center Hamburg-Eppendorf, Hamburg, Germany, ^42^Centre for Molecular Oncology - Barts Cancer Institute - Queen Mary University of London, London, UK, ^43^Humangenetik Tuebingen, Tuebingen, Germany, ^44^Cancer & Environment Group, Center for Research in Epidemiology and Population Health (CESP), Paris-Sud University, Villejuif Cédex, France, ^45^Department of Epidemiology, Harvard T. H. Chan School of Public Health, Boston, MA, USA, ^46^Division of Population Health, Health Services Research and Primary Care, University of Manchester, Manchester, UK, ^47^Warwick Medical School, University of Warwick, Coventry, UK, ^48^Nuffield Department of Surgical Sciences, University of Oxford, John Radcliffe Hospital, Oxford, UK, ^49^Department of Oncology, University of Cambridge, Addenbrooke's Hospital, Cambridge, UK, ^50^Cancer Research UK, Cambridge Research Institute, Li Ka Shing Centre, Cambridge, UK, ^51^Department of Population Sciences, Beckman Research Institute of the City of Hope, Duarte, CA, USA, ^52^Division of Public Health Sciences, Fred Hutchinson Cancer Research Center, Seattle, WA, USA, ^53^Department of Urology, University of Washington, Seattle, WA, USA, ^54^Faculty of Health and Medical Sciences, University of Copenhagen, Copenhagen, Denmark, ^55^Department of Clinical Biochemistry, Herlev and Gentofte Hospital, Copenhagen University Hospital, Copenhagen, Denmark, ^56^The University of Surrey, Guildford, UK, ^57^Department of Cancer Epidemiology, Moffitt Cancer Center, Tampa, FL, USA, ^58^Department of Applied Health Research, University College London, London, UK, ^59^Department of Public Health and Primary Care, Department of Oncology, Centre for Cancer Genetic Epidemiology, University of Cambridge, Strangeways Laboratory, Cambridge, UK, ^60^Department of Medicine, Channing Division of Network Medicine, Brigham and Women's Hospital/Harvard Medical School, Boston, MA, USA, ^61^Department of Surgery, Faculty of Medicine, University of Malaya, Kuala Lumpur, Malaysia, ^62^Department of Epidemiology and Biostatistics, School of Public Health, Imperial College London, London, UK, ^63^Department of Urology, Erasmus University Medical Center, Rotterdam, The Netherlands, ^64^Department of Radiation Oncology, Icahn School of Medicine at Mount Sinai, New York, NY, USA, ^65^Department of Genetics and Genomic Sciences, Icahn School of Medicine at Mount Sinai, New York, NY, USA, ^66^Institute of Biomedicine, University of Turku, Turku, Finland, ^67^Department of Medical Genetics, Laboratory Division, Turku University Hospital, Turku, Finland, ^68^Department of Molecular Medicine, Aarhus University Hospital, Aarhus, Denmark, ^69^Department of Clinical Medicine, Aarhus University, Aarhus, Denmark, ^70^Department of Epidemiology, The University of Washington School of Public Health, Seattle, WA, USA, ^71^SWOG Statistical Center, Fred Hutchinson Cancer Research Center, Seattle, WA, USA, ^72^Department of Genetics, Portuguese Oncology Institute of Porto, Porto, Portugal, ^73^Biomedical Sciences Institute (ICBAS), University of Porto, Porto, Portugal, ^74^Department of Laboratory Medicine and Pathology, Mayo Clinic, Rochester, MN, USA, ^75^Division of Cancer Sciences, NIHR Manchester Biomedical Research Centre, Manchester Academic Health Science Centre, University of Manchester, Manchester, UK, ^76^Nuffield Department of Population Health, Cancer Epidemiology Unit, University of Oxford, Oxford, UK, ^77^Department of Oncology, Cross Cancer Institute, University of Alberta, Edmonton, Alberta, Canada, ^78^Division of Radiation Oncology, Cross Cancer Institute, University of Alberta, Edmonton, Alberta, Canada, ^79^Fundación Pública Galega de Medicina Xenómica-SERGAS, Santiago de Compostela, Spain, ^80^Division of Cancer Sciences, Manchester Academic Health Science Centre, University of Manchester, The Christie Hospital NHS Foundation Trust, Manchester, UK, ^81^Division of Nutritional Epidemiology, Institute of Environmental Medicine, Karolinska Institutet, Stockholm, Sweden, ^82^Department of Surgical Sciences, Uppsala University, Uppsala, Sweden

**Renal Cancer GWAS**

Demetrius Albanes^1^, Garnet L. Anderson^2^, Gabriella Andreotti^1^, John G. Anema^3^, Rosamonde E. Banks^4^, Poulami Barman^5^, Laura E. Beane Freeman^1^, Vladimir Bencko^6^, Simone Benhamou^7^, Celine Besse^8^, Amanda Black^1^, Helene Blanche^9^, Anne Boland^8^, Kevin M. Brown^1^, Fiona Bruinsma^10^, H.B. Bueno-De-Mesquita^11^, Laurie Burdette^1^, Julie Buring^12^, Geraldine Cancel-Tassin^13^, Frederico Canzian^14^, Hallie Carol^15^, Robert Carreras-Torres^16^, Eunyoung Cho^17^, Toni K. Choueiri^15^, Wong-Ho Chow^18^, Peter E. Clark^19^, Leandro M. Colli^1^, Olivier Cussenot^13^, Cezary Cybulski^20^, Jean-Francois Deleuze^21,9^, Eric J. Duell^22^, Todd E. Edwards^19^, Timothy Eisen^23^, Eleonora Fabianova^24^, Tony Fletcher^25^, Matthieu Foll^16^, Lenka Foretova^26^, Matthew L. Freedman^15^, Neal D. Freedman^1^, Valerie Gaborieau^16^, Susan M. Gapstur^27^, J Michael. Gaziano^28^, Marc Henrion^29^, Jonathan N. Hofmann^1^, Ivana Holcatova^6^, Wen-Yi Huang^1^, Kristian Hveem^30^, Vladimir Janout^31^, Viorel Jinga^32^, Mattias Johansson^16^, Lisa Johnson^2^, Susan Jordan^33^, Richard J. Kahnoski^3^, Kvetoslava Koppova^24^, Stella Koutros^1^, Brian R. Lane^3^, James Larkin^34^, Susanna C. Larsson^35^, G Mark. Lathrop^36^, I-Min Lee^12^, Bradley C. Leibovich^5^, Peng Li^16^, Loren Lipworth^19^, Jolanta Lissowska^37^, Börje Ljungberg^38^, Jan Lubinski^20^, Juhua Luo^39^, Mitchell J. Machiela^1^, Satu Mannisto^40^, Dana Mates^41^, Mirjana Mijuskovic^42^, Lee E. Moore^1^, Anush Mukeriya^43^, Marie Navratilova^26^, Sabrina L. Noyes^44^, Miodrag Ognjanovic^45^, David Petillo^44^, Mark M. Pomerantz^15^, Mark A. Preston^28^, Egor Prokhortchouk^46^, Stefan Rascu^32^, Peter Rudnai^47^, Joshua N. Sampson^1^, Sharon A. Savage^1^, Peter J. Selby^4^, Howard S. Sesso^12^, Gianluca Severi^7^, Raviprakash T. Sitaram^38^, Konstantin G. Skryabin^46^, Victoria L. Stevens^27^, Neonila Szeszenia-Dabrowska^48^, Bin Tean Teh^44^, Lars J. Vatten^49^, Zhaoming Wang^50^, Stephanie Weinstein^1^, Emily White^2^, Kathryn M. Wilson^12^, Alicja Wolk^35^, Christopher Wood^18^, Yuanqing Ye^18^, Meredith Yeager^1^, David Zaridze^43^

^1^Division of Cancer Epidemiology and Genetics, National Cancer Institute, National Institutes of Health, Rockville, MD, USA, ^2^Fred Hutchinson Cancer Research Center, Seattle, WA, USA, ^3^Division of Urology, Spectrum Health, Grand Rapids, MI, USA, ^4^University of Leeds, Leeds, UK, ^5^Department of Urology, Mayo Clinic, Rochester, MN, USA, ^6^Charles University, Prague, Czech Republic, ^7^INSERM, Villejuif, France, ^8^Centre National de Genotypage, Evry, France, ^9^Fondation Jean Dausset-Centre d'Etude du Polymorphisme Humain, Paris, France, ^10^Cancer Epidemiology Division, Cancer Council Victoria, Melbourne, Australia, ^11^National Institute forPublic Health and the Environment, Bilthoven, The Netherlands, ^12^Harvard T.H. Chan School of Public Health, Boston, MA, USA, ^13^Groupe de Recherche GRC-UPMC n°5, Centre de Recherche sur les Pathologies Prostatiques et Urologiques (CeRePP), Paris, France, ^14^Genomic Epidemiology Group, German Cancer Research Center, Heidelberg, Germany, ^15^Dana-Farber Cancer Institute, Boston, MA, USA, ^16^International Agency for Research on Cancer (IARC), Lyon, France, ^17^Brown University, Providence, RI, USA, ^18^Department of Epidemiology, The University of Texas MD Anderson Cancer Center, Houston, TX, USA, ^19^Vanderbilt-Ingram Cancer Center, Nashville, TN, USA, ^20^Pomeranian Medical University, Szczecin, Poland, ^21^Centre National de Genotypage, Institut de Genomique, Centre de l'Energie Atomique et aux Energies Alternatives, Paris, France, ^22^Catalan Institute of Oncology, Barcelona, Spain, ^23^University of Cambridge, Cambridge, UK, ^24^Regional Authority of Public Health in Banska Bystrica, Banska Bystrica, Slovakia, ^25^London School of Hygiene and Tropical Medicine, University of London, London, UK, ^26^Department of Cancer Epidemiology and Genetics, Masaryk Memorial Cancer Institute, Brno, Czech Republic, ^27^American Cancer Society, Atlanta, GA, USA, ^28^Brigham and Women's Hospital, Boston, MA, USA, ^29^Icahn School of Medicine at Mount Sinai, New York, NY, USA, ^30^Norwegian University of Science and Technology, Levanger, Sweden, ^31^Palacky University, Olomouc, Czech Republic, ^32^Carol Davila University of Medicine and Pharmacy, Bucharest, Romania, ^33^QIMR Berghofer Medical Research Institute, Brisbane, Australia, ^34^The Institute for Cancer Research, London, UK, ^35^Institute of Environmental Medicine, Karolinska Institutet, Stockholm, Sweden, ^36^Genome Quebec Innovation Centre, McGill University, Montreal, Canada, ^37^The M Sklodowska-Curie Cancer Center and Institute of Oncology, Warsaw, Poland, ^38^Umeå University, Umeå, Sweden, ^39^Department of Epidemiology and Biostatistics, School of Public Health Indiana University, Bloomington, IN, USA, ^40^National Institute for Health and Welfare, Helsinki, Finland, ^41^National Institute of Public Health, Bucharest, Romania, ^42^Military Medical Academy, Belgrade, Serbia, ^43^Russian N.N. Blokhin Cancer Research Centre, Moscow, Russia, ^44^Van Andel Research Institute, Grand Rapids, MI, USA, ^45^International Organization for Cancer Prevention and Research, Belgrade, Serbia, ^46^Centre ‘Bioengineering’ of the Russian Academy of Sciences, Moscow, Russia, ^47^National Public Health Center, National Directorate of Environmental Health, Budapest, Hungary, ^48^Department of Epidemiology, Institute of Occupational Medicine, Lodz, Poland, ^49^Norwegian University of Science and Technology, Trondheim, Norway, ^50^St. Jude Children's Research Hospital, Memphis, TN, USA

**TECAC**

Victoria Cortessis^1^, Jourik A. Gietema^2^, Ramneek Gupta^3^, Trine B. Haugen^4^, Michelle A.T. Hildebrandt^5^, Robert Karlsson^6^, Kevin Litchfield^7^, Nandita Mitra^8^, Ewa Rajpert-De Meyts^9^, Stephen M. Schwartz^10^, Rolf I. Skotheim^11^, Saran Vardhanabhuti^12^, Zhaoming Wang^13^

^1^Department of Preventive Medicine, University of Southern California, Los Angeles, USA, ^2^Department of Medical Oncology, University Medical Center Groningen, Groningen, The Netherlands, ^3^Department of Health Technology, Technical University of Denmark, Lyngby, Denmark, ^4^Health Sciences, Oslo and Akershus University College of Applied Sciences, Oslo, Norway, ^5^Department of Epidemiology, MD Anderson Cancer Center, Houston, USA, ^6^Department of Medical Epidemiology and Biostatistics, Karolinska Institute, Solna, Sweden, ^7^Cancer Evolution and Genome Instability Laboratory, The Francis Crick Institute, London, UK, ^8^Department of Biostatistics, Epidemiology and Informatics, University of Pennsylvania, Philadelphia, USA, ^9^Department of Growth and Reproduction, Copenhagen University Hospital (Rigshospitalet), Copenhagen, Denmark, ^10^Epidemiology Program, Public Health Sciences Division, Fred Hutchinson Cancer Research Center, Seattle, USA, ^11^Department of Cancer Prevention, Genome Biology Group, Institute for Cancer Research, Oslo, Norway, ^12^Department of Biostatistics, Harvard T.H. Chan School of Public Health, Boston, USA, ^13^Department of Epidemiology and Cancer Control, St. Jude Children's Research Hospital, Memphis, USA
